# Effect of natural mutations of SARS-CoV-2 on spike structure, conformation and antigenicity

**DOI:** 10.1101/2021.03.11.435037

**Authors:** Sophie M-C. Gobeil, Katarzyna Janowska, Shana McDowell, Katayoun Mansouri, Robert Parks, Victoria Stalls, Megan F. Kopp, Kartik Manne, Kevin Saunders, Robert J Edwards, Barton F. Haynes, Rory C. Henderson, Priyamvada Acharya

**Author notes:** Correspondence (R.C.H.), (P.A.).

## Abstract

New SARS-CoV-2 variants that have accumulated multiple mutations in the spike (S) glycoprotein enable increased transmission and resistance to neutralizing antibodies. Here, we study the antigenic and structural impacts of the S protein mutations from four variants, one that was involved in transmission between minks and humans, and three that rapidly spread in human populations and originated in the United Kingdom, Brazil or South Africa. All variants either retained or improved binding to the ACE2 receptor. The B.1.1.7 (UK) and B.1.1.28 (Brazil) spike variants showed reduced binding to neutralizing NTD and RBD antibodies, respectively, while the B.1.351 (SA) variant showed reduced binding to both NTD- and RBD-directed antibodies. Cryo-EM structural analyses revealed allosteric effects of the mutations on spike conformations and revealed mechanistic differences that either drive inter-species transmission or promotes viral escape from dominant neutralizing epitopes.

**Highlights:**

- Cryo-EM structures reveal changes in SARS-CoV-2 S protein during inter-species transmission or immune evasion.
- Adaptation to mink resulted in increased ACE2 binding and spike destabilization.
- B.1.1.7 S mutations reveal an intricate balance of stabilizing and destabilizing effects that impact receptor and antibody binding.
- E484K mutation in B.1.351 and B.1.1.28 S proteins drives immune evasion by altering RBD conformation.
- S protein uses different mechanisms to converge upon similar solutions for altering RBD up/down positioning.

## Introduction

The emergence of rapidly-spreading variants of the severe acute respiratory syndrome coronavirus 2 (SARS-CoV-2), the causative agent for coronavirus disease 2019 (COVID-19), threatens to prolong an already devastating pandemic with unprecedented global health and economic consequences. First identified in December 2019 in Wuhan, China, declared a Public Health Emergency of International Concern in January 2020, and a pandemic in March 2020, COVID-19 has claimed more than 2.5 million lives and infected more than 117 million people world-wide (https://coronavirus.jhu.edu). Several vaccines are being deployed worldwide to gain control of the pandemic (Baden et al., 2021; Chung et al., 2020; Polack et al., 2020), although appearance of highly transmissible variants have caused concern (Galloway et al., 2021; Leung et al., 2021) (https://www.cdc.gov/coronavirus/2019-ncov/cases-updates/variant-surveillance/variant-info.html). Some variants have exhibited resistance to neutralization by antibodies and plasma from convalescent or vaccinated individuals, which together with their fast spread have raised concerns that their resistance may render current vaccines ineffective (Wang et al., 2021b; Wibmer et al., 2021). Additionally, transmission of SARS-CoV-2 between humans and animals has been observed, most notably in mink farms leading to the culling of large populations of minks in Denmark and other countries to prevent establishment of a non-human reservoir of SARS-CoV-2 variants (Oude Munnink et al., 2021). Multiple synergizing mutations in the spike (S) glycoprotein (Ke et al., 2020; Turonova et al., 2020) in these variants are under scrutiny due to the S protein’s central role in engaging the angiotensin-converting enzyme 2 (ACE2) receptor to mediate cellular entry (Hoffmann et al., 2020a), and being the primary target of neutralizing antibodies elicited either by vaccination or during natural infection (Corbett et al., 2020; Sempowski et al., 2020).

The prefusion SARS-CoV-2 S protein trimer is composed of two subunits, S1 and S2, separated by a furin cleavage site (**Figure 1**). The S1 subunit contains the N-terminal domain (NTD), the ACE2 receptor binding domain (RBD), and two subdomains (SD1 and SD2). Both the NTD and RBD are dominant targets for neutralizing antibodies (Barnes et al., 2020a; Barnes et al., 2020b; Li et al., 2021; Yan et al., 2020). The RBD transitions between a “closed” or “down” conformation that is inaccessible to ACE2 receptor and “open” or “up” conformation that allows for recognition and binding to the host cell ACE2 receptor (Gui et al., 2017; Shang et al., 2020; Yuan et al., 2017). We and others have previously shown that mutations in distal regions of the S protein can have allosteric effects on the RBD up/down disposition (Gobeil et al., 2021; Henderson et al., 2020; Yurkovetskiy et al., 2020; Zhou et al., 2020). We have further shown that SD1 and SD2 play essential roles in modulating spike allostery (Gobeil et al., 2021). The S2 subunit contains a second protease cleavage site (S2’) for the protease TMPRSS2, followed by the fusion peptide (FP), heptad repeat 1 (HR1), the central helix (CH), the connector domain (CD), heptad repeat 2 (HR2), the transmembrane domain (TM) and a cytoplasmic tail (CT) (**Figure 1**). After binding the ACE2-receptor, and protease cleavage at the furin and TMPRSS2 cleavage sites, the S protein undergoes large conformational changes leading to cellular entry (Bestle et al., 2020; Hoffmann et al., 2020a; Hoffmann et al., 2020b; Matsuyama et al., 2020).

**Figure 1.**
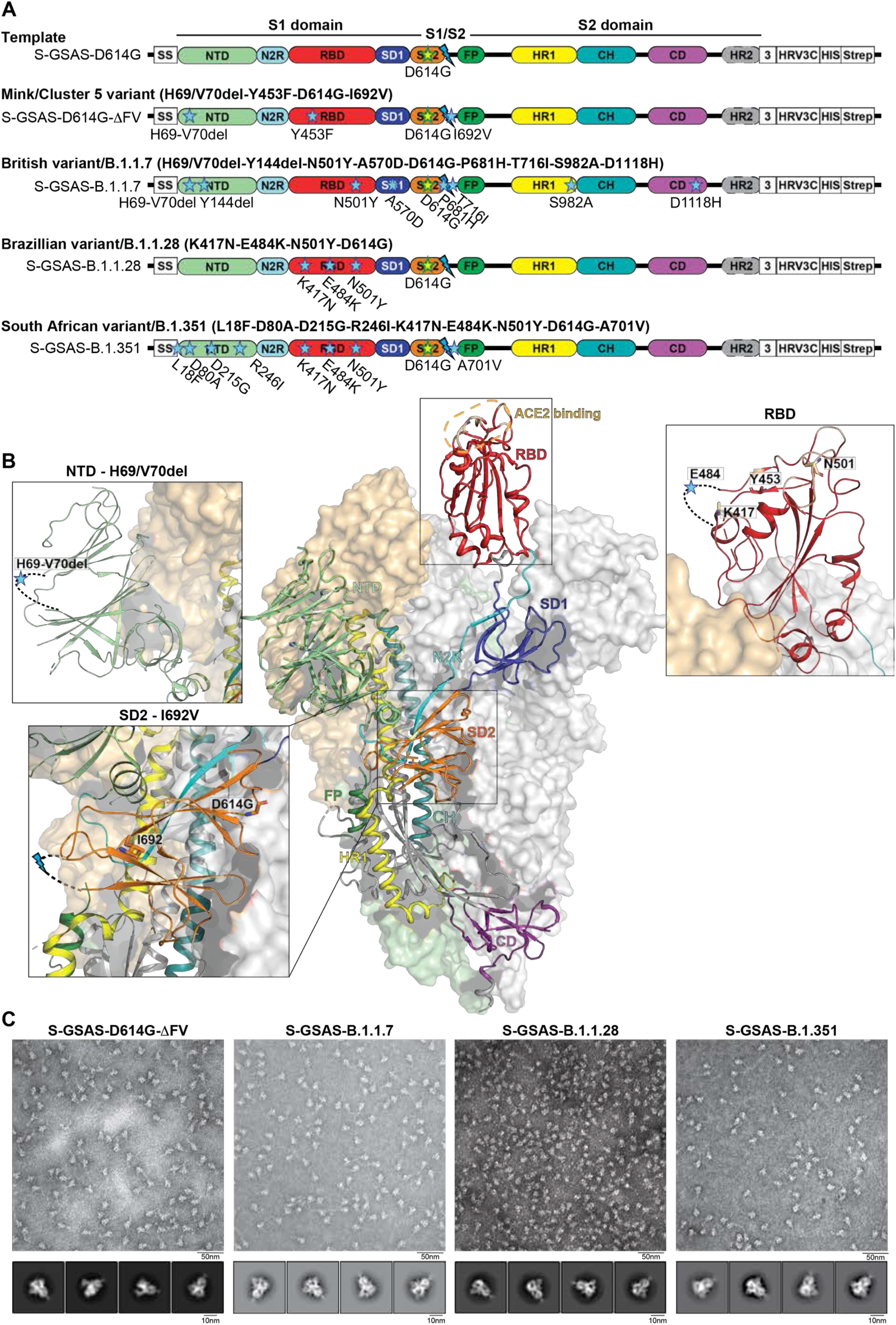
SARS-CoV-2 spike (S) protein ectodomains for characterizing structures and antigenicity of S protein variants. **A.** Domain architecture of the SARS-CoV-2 spike protomer. The S1 subunit contains a signal sequence (SS), the NTD (N-terminal domain, pale green), N2R (NTD-to-RBD linker, cyan), RBD (receptor-binding domain, red), SD1 and SD2 (subdomain 1 and 2, dark blue and orange) subdomains. The S2 subunit contains the FP (fusion peptide, dark green), HR1 (heptad repeat 1, yellow), CH (central helix, teal), CD (connector domain, purple) and HR2 (heptad repeat 2, grey) subdomains. The transmembrane domain (TM) and cytoplasmic tail (CT) have been truncated and replaced by a foldon trimerization sequence (3), an HRV3C cleavage site (HRV3C), a his-tag (His) and strep-tag (Strep). The D614G mutation is in the SD2 domain (yellow star, green contour). The S1/S2 furin cleavage site (RRAR) has been mutated to GSAS (blue lightning). **B.** Representation of the trimeric SARS-CoV-2 spike ectodomain with one RBD-up in a prefusion conformation (PDB ID 7KDL). The S1 domain on an RBD-down protomer is shown as pale orange molecular surface while the S2 domain is shown in pale green. The subdomains on an RBD-up protomer are colored according to panel **A** on a ribbon diagram. Each inset correspond to the spike regions harboring mutations included in this study. **C.** Representative NSEM micrographs and 2D class averages of S protein variants.

Fall 2020 has been marked by the appearance of several neutralization-resistant and/or highly infectious variants of SARS-CoV-2 with mutations in the S protein accumulating in the background of the D614G variant that arose early in the pandemic and quickly became dominant worldwide (Korber et al., 2020). Some S mutations recur in variants that originated independently in different parts of the world, suggesting these mutations confer selective advantages such as escape from vaccine induced or convalescent neutralizing antibodies. Here, we determined structures and antigenicity of (1) a variant that was implicated in the transmission between humans and minks (Koopmans, 2021), (2) the B.1.1.7 variant (20I/501Y.V1) that was first detected in the United Kingdom (UK) and shares the N501Y mutation with the B.1.351 and B.1.1.28 variants (Galloway et al., 2021; Leung et al., 2021), (3) the B.1.1.28 variant, first reported in Japan in four travelers from Brazil and was a dominant driver of the early epidemic phase in multiple Southeastern Brazilian states (Paiva et al., 2020), and (4) the B.1.351 variant (20H/501Y.V2) that arose independently in South Africa, and shares three RBD mutations, K417N, E484K, and N501Y, with the B.1.1.28 variant (Mwenda et al., 2021) (**Figure 1**). The P.1 lineage (20J/501Y.V3) branched off the B.1.1.28 lineage, and has been identified in an outbreak in a region in Brazil that had already seen approximately 75% of the population infected with SARS-CoV-2 as of October 2020 (Sabino et al., 2021), thus raising concerns about increase in propensity for SARS-CoV-2 re-infection of individuals. Here, we study the impact of these SARS-CoV-2 mutations on the antigenicity and structure of the variant spike proteins, and elucidate the structural mechanisms underlying the effects of these spike mutations on transmissibility and immune evasion.

## Results

### SARS-CoV-2 S protein variant constructs, protein production and quality control

For the S ectodomain constructs described in this study, we used the previously described S-GSAS-D614G S ectodomain template (**Figure 1**) (Gobeil et al., 2021). This construct includes residues 1 to 1208 of SARS-CoV-2 S, a “RRAR” to “GSAS” substitution that renders the furin cleavage site at the junction of the S1 and S2 subunits inactive, a foldon trimerization motif appended at the C-terminus of the spike sequence that ensures formation of native-like spike trimers in the absence of the transmembrane region, and a C-terminal TwinStrep tag for efficient purification and S protein immobilization on solid support.

The S protein variants described in this study were expressed, purified and assessed similarly, and as we have described previously (Edwards et al., 2021). All the purified S proteins showed similar migration profiles on SDS-PAGE and SEC, with high-quality spike preparations confirmed by negative stain electron microscopy (NSEM) (**Figure 1**, **Supplemental Item 1)**. NSEM reported between 78% and 91% prefusion spike trimers; the remaining particle picks were classified as junk.

### SARS-CoV-2 S protein variant with mink-associated cluster 5 mutations

Spillover of SARS-CoV-2 from humans to minks, and then from minks to humans was first reported in April 2020 in the Netherlands, and since has also been independently reported in Denmark, Spain, Italy, USA, Sweden, and Greece (Koopmans, 2021). This issue came into public view in the fall 2020 when a decision to cull all (∼ 17 million) minks in Denmark was announced. Five S protein mutations were observed in SARS-CoV-2 transmitted from humans to mink, and back to humans. This variant, termed “cluster 5”, includes a H69 (H69Δ) and V70 (V70Δ) deletion in the NTD, and amino acid mutations Y453F in the RBD, I692V located downstream of the furin motif in SD2, and M1229I in the transmembrane domain. In the SARS-CoV-2 S ectodomain context, we included the H69Δ, V70Δ, Y453F and I692V mutations, referred to here as S-GSAS-D614G-ΔFV (**Figure 1**). The fifth “cluster 5” mutation (M1229I) in the transmembrane region is not present in the ectodomain construct.

#### Receptor binding and antigenicity of the S-GSAS-D614G-ΔFV ectodomain

To understand the effect of the acquired mutations in the S ectodomain on its binding to human ACE2 receptor, we measured binding of the S-GSAS-D614G-ΔFV ectodomain to the recombinant human ACE2 protein by ELISA and SPR (**Figure 2A**, **Supplemental Items 2-4**). By ELISA, ACE2 showed higher levels of binding to S-GSAS-D614G-ΔFV than to the S-GSAS-D614G ectodomain **(Supplemental Item 2A)**, and SPR showed a slower off-rate of binding **(Supplemental Item 2B)**.

**Figure 2.**
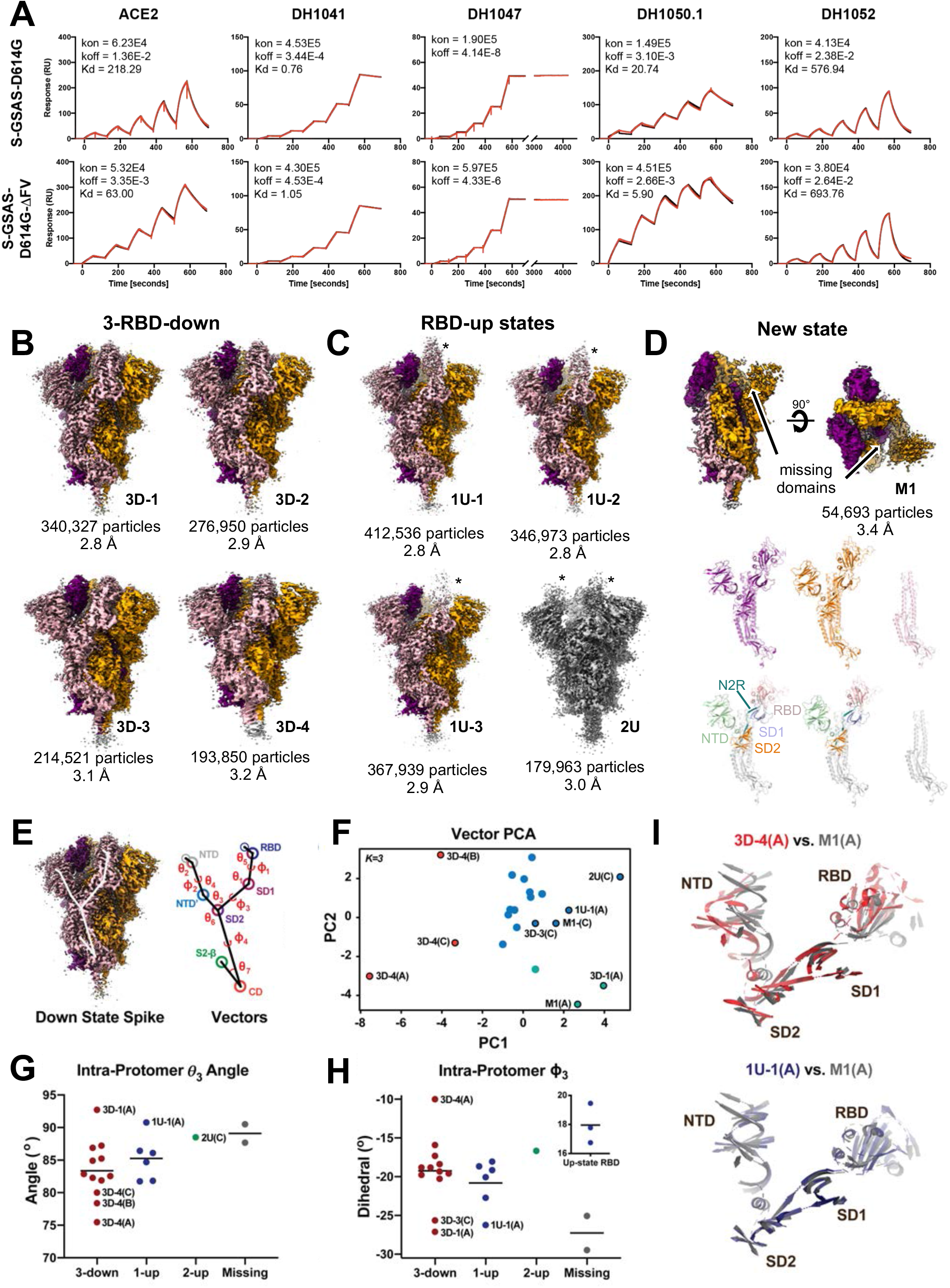
Structures and antigenicity of mink-associated S-GSAS-ΔFV ectodomain. **A.** Binding of ACE2 receptor ectodomain (RBD-directed), and antibodies DH1041 and DH1047 (RBD-directed, neutralizing), DH1050.1 (NTD-directed, neutralizing) and DH1052 (NTD-directed, non-neutralizing). to S-GSAS-D614G (top) and S-GSAS-B.1.1.28 (bottom) measured by SPR using single-cycle kinetics. The red lines are the binding sensorgrams and the black lines show fits of the data to a 1:1 Langmuir binding model. The on-rate (k_on_, M^-1^s^-1^), off-rate (k_off_, s^-1^) and affinity (K_D_, nM) for each interaction are indicated in the insets. Binding of DH1047 was too tight for accurate measurement of K_D_. **B-D.** Cryo-EM reconstructions of S-GSAS-ΔFV ectodomain colored by protomer chains **B.** 3-RBD-down states; 3D-1 (EMDB: 23549, PDB: 7LWL), 3D-2 (EMDB: 23548, PDB: 7LWK), 3D-3 (EMDB: 23546, PDB: 7LWI), 3D-4 (EMDB: 23547, PDB: 7LWJ), **C.** RBD-up states, including 3 1-RBD-up states: 1U-1 (EMDB: 23550, PDB: 7LWM), 1U-2 (EMDB: 23551, PDB: 7LWN), 1U-3 (EMDB: 23552, PDB: 7LWO), and a 2-RBD-up state (EMDB: 23553, PDB: 7LWP), **D.** A state, named M1 (EMDB: 23554, PDB: 7LWQ) lacking the S1 subunit and SD2 subdomain of one of the three protomers (EMDB: 23554, PDB 7LWQ). Top panel shows two views of the cryo-EM reconstruction rotated by 90°, middle panel shows the individual protomers colored to match the colors in the top panel, bottom panel shows the protomers with RBDs colored salmon, NTDs green, SD1 blue, SD2 orange, and the S2 subunit grey. **E-I.** Vector analysis defining changes in intra-protomer domain disposition. **E.** Schematic showing angles and dihedrals between different structural elements in the SARS-CoV-2 S protein shown in the context of a 3-RBD-down spike. **F.** Principal components analysis of the intra-protomer vector magnitudes, angles, and dihedrals colored according to placement in 3-down (red), 1-up (blue), 2-up (green), or S1 missing (grey) spike trimers. **G.** Intra-protomer ɸ_3_ dihedral angles of the NTD’ relative to the RBD about a vector connecting SD2 and SD1 **H.** Intra-protomer !_3_ angles between the NTD’, SD2, and SD1. **I.** Chain A of the M1 protomer aligned to the chain A of 3D-4 (top) and chain A of 1U-1 (bottom). The protomers were aligned on SD2.

Using single cycle kinetics, we measured a ∼3.5-fold tighter affinity for ACE2 binding to S-GSAS-D614G-ΔFV than to S-GSAS-D614G, and confirmed that the improved affinity was contributed primarily by a decrease in off-rate (**Figure 2A**, **Supplementary Table 1**). To deconvolute the effect of the different spike mutations, we measured ACE2 binding to S-GSAS-D614G-H69/V70Δ, S-GSAS-D614G-Y453F and S-GSAS-D614G-I692V **(Supplemental Item 4)**. Of these, S-GSAS-D614G-Y453F showed a similar kinetics and affinity profile for ACE2 binding as the S-GSAS-D614G-ΔFV spike, thus implicating the Y453F mutation in the enhanced ACE2 binding affinity of S-GSAS-D614G-ΔFV.

To understand the effect of the mutations on spike antigenicity, we measured binding affinity and kinetics of S-GSAS-D614G-ΔFV to a panel of antibodies targeting different regions of the spike (**Figure 2A**, **Supplementary Table 1, Supplemental Items 3-4**) (Acharya et al., 2020; Li et al., 2021; Yuan et al., 2017). S-GSAS-D614G-ΔFV retained robust binding to RBD-directed neutralizing antibodies, DH1041, DH1043 and DH1047. We observed improved binding affinity of S-GSAS-D614G-ΔFV compared to S-GSAS-D614G to the neutralizing NTD-directed antibodies DH1050.1 and DH1050.2 by 3.5 and 2.6-fold, respectively. This was primarily a result of an increased on-rate, with the dominant contribution coming from the H69/V70Δ mutation (**Figure 2A**, **Supplemental Item 4)**. Thus, our binding studies showed that inter-species adaptation involved enhancement of receptor binding affinity of the S protein without evidence of escape from the dominant neutralization epitopes.

#### Cryo-EM structures of the S-GSAS-D614G-ΔFV ectodomain

To understand how the mutations acquired during interspecies transmission between mink and humans affected S protein conformation, we determined cryo-EM structures of the S-GSAS-D614G-ΔFV ectodomain. From the cryo-EM dataset, we classified and refined four 3-RBD-down populations to overall resolutions of 2.8-3.2 Å, without imposing 3-fold symmetry, and named them 3D-1, 3D-2, 3D-3 and 3D-4 (PDB 7LWK, 7LWL, 7LWI and 7LWJ respectively) (**Figure 2B**). We also refined three 1-RBD-up populations, 1U-1, 1U-2 and 1U-3 (PDB 7LWM, 7LWN and 7LWO respectively), to resolutions of 2.8-2.9 Å, and one 2-RBD-up population (2U; PDB 7LWP) to 3.0 Å (**Figure 2C**, **Supplementary Table 2, Supplemental Items 5 and 6**). We identified a spike population, named M1, with two RBDs in “down” position, and no density visible for the entire S1 subunit of the third protomer, both in the 3.2 Å resolution reconstruction and the gaussian filtered map (**Figure 2D**, **Supplementary Table 2, Supplemental Items 5 and 6**). Since SDS-PAGE did not show any evidence for proteolysis, the absence of the S1 subunit density for one protomer suggests either shearing of this region from the rest of the spike during cryo-EM specimen vitrification, or unfolding of this region.

To examine details of the structural variability, the refined maps were fitted with coordinates. Overlay of the 3-RBD-down structures, using S2 residues 908-1035 of the HR1-CH region for the superposition, revealed that in addition to the expected variation in the S1 subunits (Gobeil et al., 2021), there was considerable variability in S2, including the region near the HR1-CH hinge **(Supplemental Item 7A)**. This variability was especially pronounced for the 3D-4 structure (**Figure 2B and Supplemental Item 7)**. By contrast, superposition of the three 1-RBD-up structures using residues 908-1035 yielded good structural alignment in the S2 subunit, with the S1subunit showing variability, especially in mobile RBD and NTD regions (**Supplemental Item 8**).

To obtain residue-level visualization of the differences between the structures, we performed difference distance matrices (DDM) analysis (Richards and Kundrot, 1988). DDM analyses provide superposition-free comparisons between a pair of structures by calculating the differences between the distances of each pair of C*α* atoms in a structure and the corresponding pair of C*α* atoms in a second structure. In our analysis, we first compared each protomer with the other two protomers within the same spike, and next, compared each protomer to the protomers from the other 3-RBD-down structures (**Figure 2B and Supplemental Item 7**). The DDM analysis revealed that 3D-4 was the most asymmetric of the four 3-RBD-down structures, with large variations in the S1 subunit between its three protomer, especially in the mobile NTD and RBD regions. In addition, 3D-4 was also most different from the other three 3-RBD-down structure with large differences in the S2 region. The variation in the S2 region between different pre-fusion structures was unexpected since in prior studies the S2 subunit had appeared relatively invariable, with variability mostly contained within the S1 subunit. By contrast, the DDM analysis of the 1-up structures showed large movements in the S1 subunits, but little change in the S2 subunit, similar to prior observations with other 1-RBD-up structures, including of the S-GSAS-D614G spike (**Supplemental Item 8**) (Gobeil et al., 2021).

#### Vector Analysis of the S-GSAS-D614G-ΔFV ectodomain structures

We next analyzed the S1 and S2 domain dispositions within each protomer via calculation of our previously described vector relations (**Figure 2E-I**) (Henderson et al., 2020). Principal components analysis (PCA) of the intra-protomer angles and distances for the down state protomers revealed protomers from the 3D-4 structure formed a distinct cluster, consistent with the DDM analysis that showed 3D-4 to be a distinct structure, different from the other 3-RBD-down structures (**Figure 2F**). The two protomers from the M1 structure, with their RBDs in the down position, displayed similarity to the 3D-1 protomers along the primary component, while the 3D-3 chain C protomer closely matched the M1 chain C protomer and the 3D-1 chain A protomer closely matched the M1 chain A structure.

The I692V mutation in the S-GSAS-D614G-ΔFV ectodomain occurs in SD2, a region we had previously shown to be a conformational anchor separating the mobile NTD and RBD regions of a protomer (**Figure 1**). Small changes in SD2 can translate to large changes in the NTD and RBD regions (Gobeil et al., 2021). The I692 residue in the S-GSAS-D614G structures contacts residue P600, and loss of the methyl group at residue 692 due to the I692V substitution resulted in P600 and V692 being placed farther apart (**Supplemental Item 9**). While the map density in this region was well-defined for three of the 3-RBD-down structures (3D-1, 3D-2 and 3D-3), the 3D-4 cryo-EM map showed disorder in this region, as well as the largest separation between P600 and V692. Thus, given the critical role of the SD2 subdomain, and potential destabilization in this region due to the I692V mutation in the down-state protomers, we inspected angular disposition of the NTD′ and RBD about a vector connecting SD2 and SD1 (ɸ3) and the angle formed by the NTD′, SD2, and SD1 centroids (*θ*3; **Figure 2G-H**). These were determined for the down state protomers in the 3-down, 1-up, and 2-up structures. Consistent with the DDM and PCA results, the 3D-4 protomers occupied a distinct cluster in the ɸ3 and *θ*3 angles; in particular the 3D-4 chain A protomer ɸ3 dihedral differed markedly from the primary cluster in a direction toward that observed in up-state protomers (**Figure 2H**, **inset).** Comparison of the *θ*3 angles indicated 3D-1 chain A, 1U-1 chain A, as well as the 2-up 2U-1 chain C displayed similarity to the M1 protomers. The 3D-3 chain C and 3D-1 chain A, and 1U-1 chain A protomers displayed similar ɸ3 dihedrals to the M1 protomers (**Figure 2G-H**). Comparing the 3D-4 chain A S1 subunit structure to that of the M1 chain A demonstrates the marked differences between these structures while alignment of M1 chain A S1 subunit to 1U-1 chain A shows their similarity (**Figure 2I**). Together, this analysis suggests that loss of a single protomer’s S1 density in M1 allows the trimer to relax into a more stable 1-up like state. This is likely a result of the conformational changes indicated by the ɸ3 and *θ*3 angles emanating from internal SD2 domain differences mediated by the I692V mutation.

#### Summary

Overall, the binding and structural data on the S-GSAS-D614G-ΔFV S ectodomain suggest that the interspecies adaptation involved improved receptor binding affinity mediated primarily by the Y453F substitution in the receptor binding motif (RBM), as well as an increased propensity for open states of the S protein. We detected intriguing variability in the 3-RBD-down states, with conformational changes detected in the S2 subunit that had typically been known to remain invariant in prefusion SARS-CoV-2 S structures. Finally, we identified a new state that was missing the S1 subunit of one of its three protomer. The presence of this state and the unusual variability in the 3-RBD-down states, suggests destabilization of the pre-fusion spike structure in the S-GSAS-D614G-ΔFV S protein.

### The SARS-CoV-2 S protein B.1.1.7 variant

The B.1.1.7 variant emerged in South East England in September 2020 and rapidly became dominant in the UK. This variant has now spread to over 50 countries with reports of increased transmissibility, virulence and mortality (Davies et al., 2021). The B.1.1.7 variant contains 8 mutations in the S protein (in addition to the D614G mutation). The mutations spanned the NTD (ΔH69/V70 and ΔY144 deletions), the receptor binding motif (RBM) in the RBD (N501Y), the SD1 subdomain (A570D), the SD2 subdomain (P681H, proximal to the furin cleavage site), the T716I substitution upstream of the fusion peptide, the HR1 region (S982A), and the CD domain (D1118H). The NTD ΔH69/V70 deletion is shared with the mink-associated mutation. The B.1.1.7 variant has been shown to be susceptible to neutralizing antibodies elicited by current vaccines, as well as to RBD-directed antibodies DH1041, DH1043 and DH1047 (Li et al., 2021; Shen et al., 2021). The B.1.1.7 variant shows increased resistance to NTD-directed antibodies including 4A8 (PDB: 7C2L), 5-24, 4-8 and DH1050.1 (PDB: 7LCN) (Li et al., 2021; Liu et al., 2020; Yan et al., 2020), and this was attributed to the ΔY144 deletion that is part of a loop forming an antigenic supersite in the NTD (Wang et al., 2021a).

#### Receptor binding and antigenicity of the S-GSAS-B.1.1.7 S ectodomain

To elucidate the antigenic and structural impacts of the S protein mutations, we constructed an S ectodomain with all 8 mutations of the B.1.1.7 variant (**Figure 1A**). Consistent with previous reports, we measured a substantial drop in the binding level of the NTD-directed antibody DH1050.1 both by ELISA and SPR (**Figure 3A**, **Supplemental Item 2-4**). Although much reduced, the binding was not completely knocked out, and the SPR profile showed retention of high affinity binding (nM) with a similar kinetic profile as for S-GSAS-D614G (**Figure 3A**, **Supplementary Table 1)**. Binding of the B.1.1.7 construct to ACE2 showed ∼5-fold improvement in affinity relative to the S-GSAS-D614G, contributed primarily by the N501Y substitution in the receptor binding motif of the RBD. This is consistent with reports identifying residue 501 as a hotspot for mutagenic modulation of ACE2 binding affinity with the N501F substitution increasing ACE2 binding in deep mutational scanning studies (Starr et al., 2020). Adaptation of SARS-CoV-2 in BALB/c mice for vaccine efficacy testing resulted in N501Y selection, further supporting a functional role for this mutation (Gu et al., 2020). The N501Y substitution, either on its own or in combination with the NTD H69/V70 deletion or the SD2 P681H mutation, does not substantially affect serum neutralization elicited by current vaccines (Wang et al., 2021b; Wu et al., 2021; Xie et al., 2021). These results, together with the observed resistance to NTD Abs, and robust binding of RBD antibodies, suggest that, although the NTD antigenic supersite has been rendered ineffective in the B.1.1.7 variant, RBD-directed antibodies elicited by the current vaccines remain active against this variant.

**Figure 3.**
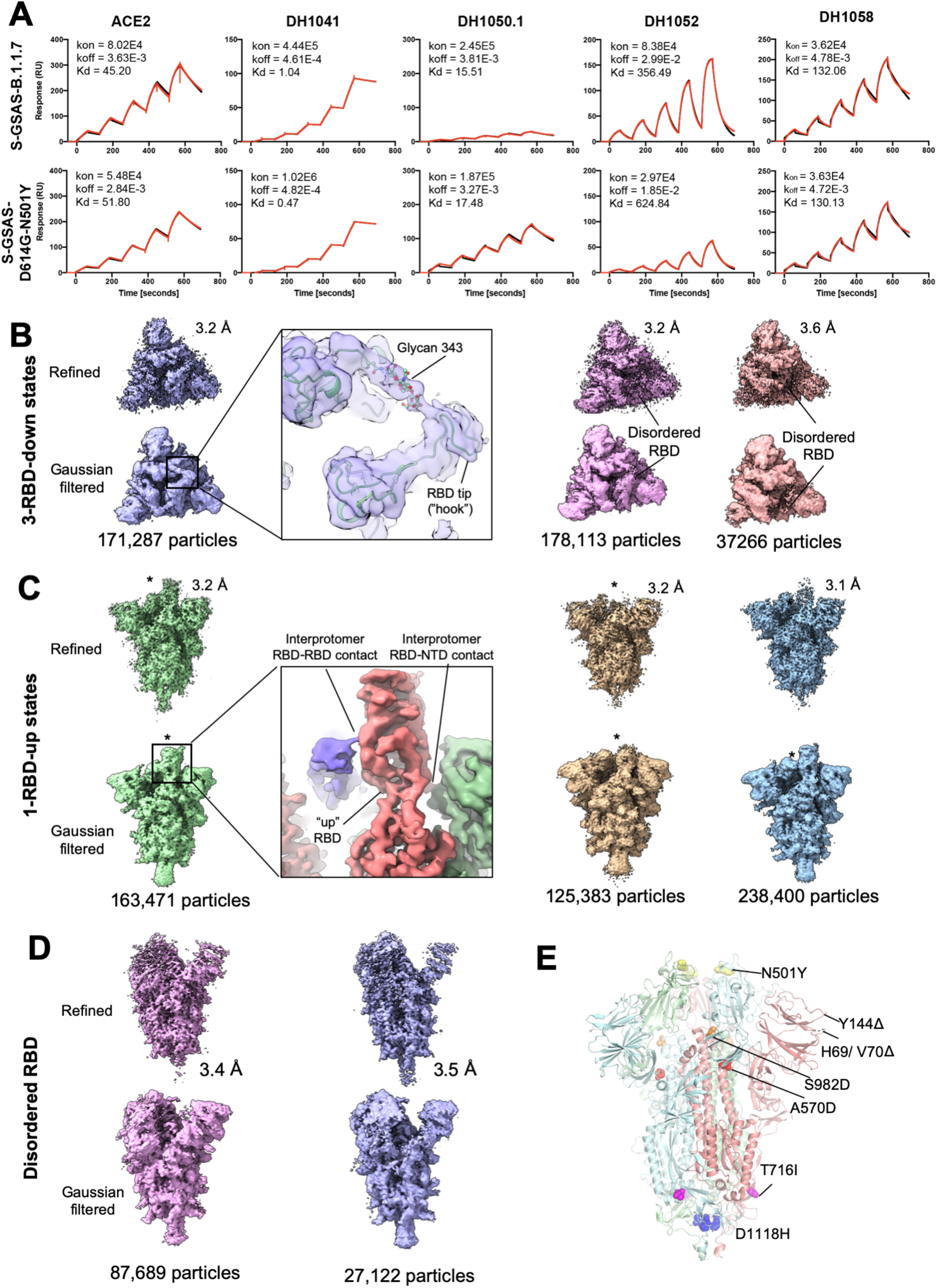
Antigenicity and structures of the S-GSAS-B.1.1.7 ectodomain. **A.** Binding of ACE2 receptor ectodomain (RBD-directed), and antibodies DH1041 and DH1047 (RBD-directed, neutralizing), DH1050.1 (NTD-directed, neutralizing) and DH1052 (NTD-directed, non-neutralizing). to S-GSAS-B.1.1.7 (top) and S-GSAS-N501Y (bottom) measured by SPR using single-cycle kinetics. The red lines are the binding sensorgrams and the black lines show fits of the data to a 1:1 Langmuir binding model. The on-rate (k_on_, M^-1^s^-1^), off-rate (k_off_, s^-1^) and affinity (K_D_, nM) for each interaction are indicated in the insets. Binding of DH1047 was too tight for accurate measurement of K_D_. **B-E.** Cryo-EM reconstructions of **B.** 3-RBD-down states, **C.** 1-RBD-up states, **D.** 1-RBD-up states with disordered RBD, **E.** Fitted coordinates into the S-GSAS-B.1.1.7 3-RBD-down cryo-EM reconstruction (PDB:7LWS), with mutations shown in spheres.

#### Cryo-EM structures of the S-GSAS-B.1.1.7 S ectodomain

To visualize the impact of the B.1.1.7 mutations on spike conformation, we solved cryo-EM structures of the S-GSAS-B.1.1.7 S ectodomain (**Figure 3B-D**, **Supplementary Table 2, Supplemental Items 10-12**). We performed 3D-classification of the dataset to yield multiple populations of the 3-RBD-down and RBD-up states. We performed asymmetric reconstruction of three populations of the 3-RBD-down state that were refined to 3.2-3.6 Å (**Figure 3B**, **Supplementary Table 2, Supplemental Item 10-11**). Each of these states revealed asymmetry in their three RBDs, with one of the RBDs showing weaker density than the other two (**Figure 3B**), indicative of enhanced mobility. An RBD in its down state makes interprotomer contacts with an adjacent NTD and with RBD glycan 343 of the same protomer. (**Figure 3B**, **inset**) (Sztain et al., 2021). Transition from the “down” to “up” state results in replacement of these contacts with differing RBD-to-NTD and RBD-to-RBD contacts (**Figure 3C**, **inset**). The apparent increase in RBD mobility in the 3-down-state of the B.1.1.7 mutant suggested a reduced barrier to the transition to the “up” state due to a weakening of the down state contacts. Consistent with this observation, the S-GSAS-B.1.1.7 cryo-EM dataset revealed a higher proportion of RBD-up state particles compared to the S-GSAS-D614G dataset (∼1.8:1 for RBD-up/RBD-down for S-GSAS-B.1.1.7 versus ∼1:1 for S-GSAS-D614G) (Gobeil et al., 2021). The ΔH69/V70 and ΔY144 deletions may contribute, in part, to this result via modification of the NTD position. In addition to RBD/NTD disorder in the 3-RBD-down states, and the identification of populations that were refined to yield typical 1-RBD-up structures (**Figure 3C)**, we also isolated two 1-RBD-up populations in which the up state RBD density was comparatively weak (**Figure 3D)**. This weak density was accompanied by considerable mobility in its adjacent NTD as inferred by its weaker density. We also identified states with 2-or 3-RBD up (**Supplemental Item 10G**) but limitations of particle numbers and preferred orientations precluded high resolution reconstructions of these populations. Unlike the mink-associated S-GSAS-D614G-ΔFV ectodomain structures, DDM analysis of the S-GSAS-B.1.1.7 structures did not show variability in the S2 subunit **(Supplemental Item 12)**.

Taken together, our results from the antigenicity assays and the cryo-EM structures are consistent with other studies that have reported impairment of the NTD antigenic supersite in the B.1.1.7 variant (Wang et al., 2021a), while retaining binding to most RBD-directed antibodies. Binding to the cross-reactive fusion peptide directed antibody DH1058 was also retained (**Figure 3A) (Li et al., 2021)**. Our results are also consistent with the reported enhancement of ACE2 binding via the N501Y mutation (Starr et al., 2020). We observed higher propensity for RBD-up states in the cryo-EM dataset, evidenced both by an increased percentage of the total particles adopting RBD-up state, as well as appearance of 2-up and 3-up states that were not detected in the S-GSAS-D614G dataset (Gobeil et al., 2021).

We next sought to understand the role of the B.1.1.7 variant mutations distal from the mobile RBD/NTD region. These mutations spanned multiple domains including the SD1 (A570D), SD2 (P681H), HR1 (S982A) and CD (D1118H) and the linker region between SD2 and fusion peptide (T716I) (**Figure 1A**). The P681H mutation, which is located in the SD2 subdomain and proximal to the furin cleavage site, could not be visualized due to the disorder in that region of the density. We found that the D1118H mutation resulted in the formation of a symmetric histidine triad near the base of the spike (**Figure 3E**, **4A-B**). Although the histidines from the three protomers were positioned farther from each other than what would enable direct hydrogen bonding, water mediated interactions would be feasible at this separation. Moreover, the cryo-EM reconstructions showed evidence for alternate conformations (**Supplemental Item 11**) that could potentially place the histidines from the different protomers into closer proximity; thus, the D1118H mutation appears to be a stabilizing mutation. In contrast, the T716I mutation appears destabilizing, resulting in the loss of a hydrogen bond between theThr716 side chain makes and Gln1071 main chain carbonyl (**Figure 4C-D**). We next examined the two mutations, A570D and S982A (**Figure 4E-I**), which appeared to be counterposing. In the S-GSAS-D614G spike, the SD1 loop containing A570 stacks against the hydrophobic face of a HR1 helix (**Figure 4F**), and the S982 residue in the HR1 domain hydrogen bonds with the side chain of T547 in SD1 (**Figure 4H**). Mutating A570 to Asp reinforced the stacking between the loop and the HR1 helix with the D570 side chain forming a hydrogen bond with the side chain of N856 (**Figure 4F**). The S982A mutation, in contrast, results in the loss of a hydrogen bond that the side chain of S982 makes with the SD1 residues T457 (**Figure 4G-H**). Comparing this region in the “down” (PDB: 7KDH) and “up” (PDB: 7KDL) protomers of S-GSAS-D614G showed a ∼5 Å shift in the position of this loop with the loop in the “up” protomer moving farther away from, and no longer within hydrogen bonding distance of S982 (**Figure 4H**). Thus, the S982A mutation is akin to disabling a latch that modulates the “up” and “down” RBD dispositions, thereby increasing the propensity of the RBD to adopt the up-state position. (**Figure 4I**).

**Figure 4.**
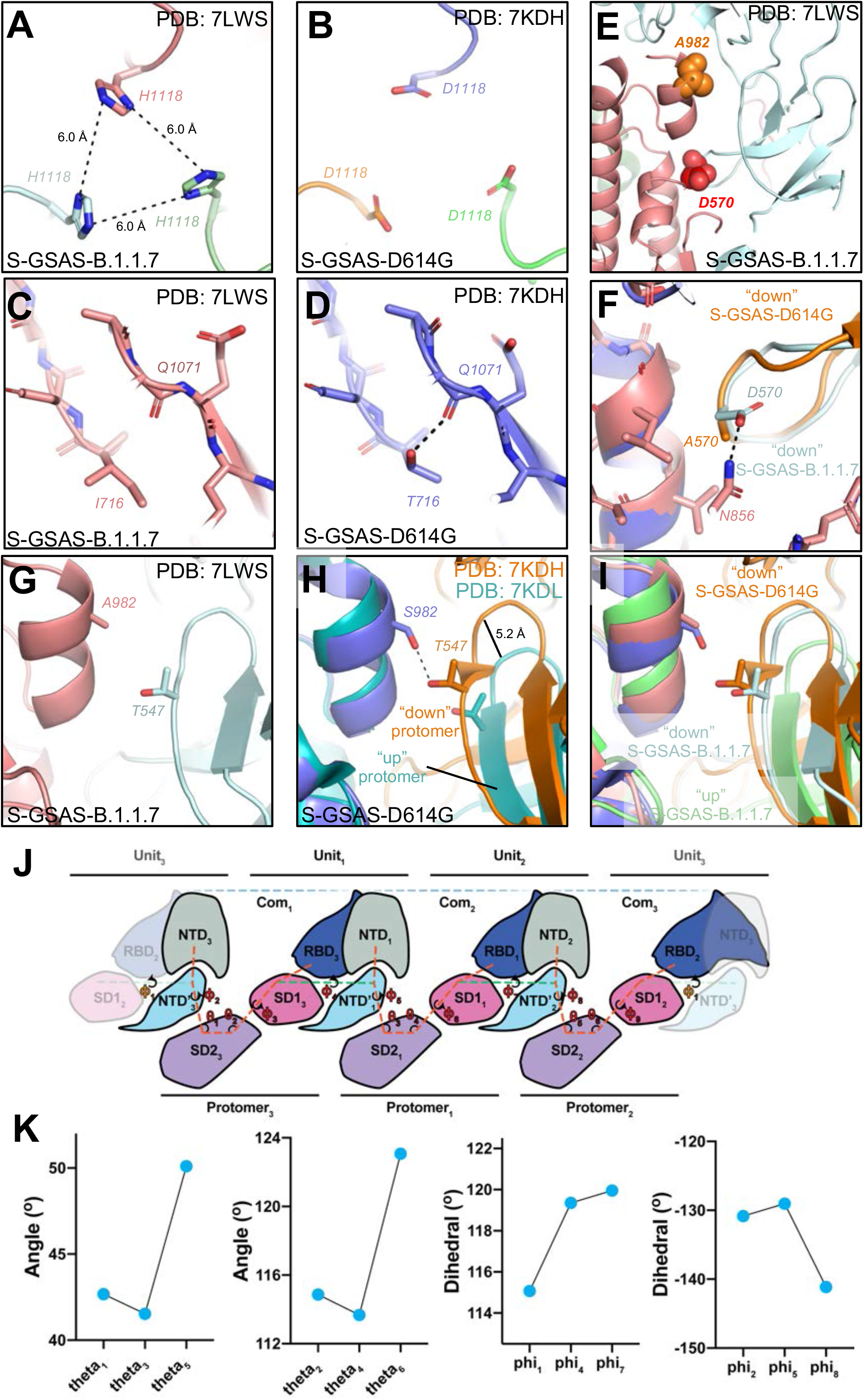
Details of S-GSAS-B.1.1.7 mutations. **A.** H1118 in S-GSAS-B.1.1.7 (PDB: 7LWS). **B.** D1118 in S-GSAS-D614G (PDB:7DKH). **C.** I716 in S-GSAS-B.1.1.7, **D.** T716 in S-GSAS-D614G; dotted line shows H-bond with backbone carbonyl OF Q1071. **E.** Zoomed-in view of the region of the A570D (red spheres) and S982A (orange spheres) mutations in S-GSAS-B.1.1.7. S protein protomers are colored light cyan and salmon. **F.** Overlay of 3-RBD-down structures of S-GSAS-D614G (PDB: 7DKH) and S-GSAS-B.1.1.7 (PDB:7LWS). **G.** S-GSAS-B.1.1.7 (PDB:7LWS) showing zoomed-in view of region around S982A mutation. Two protomers are shown in salmon and pale cyan. Residues A982 and T547 are shown in sticks. **H.** Overlay of 3-RBD-down (PDB: 7KDH) and 1-RBD-up (PDB:7KDL) structures of S-GSAS-D614G Zoomed-in view showing loss in H-bond between T547 and S982 on transition from “down” (PDB: 7KDH) to “up” (PDB: 7KDL) state. **I.** Overlay of 3-RBD-down structures of S-GSAS-D614G (PDB: 7DKH) and S-GSAS-B.1.1.7 (PDB:7LWS), and 1-RBD-up structure of S-GSAS-B.1.1.7 (PDB:7LWV). Residues 908-1035 were used for the overlays. **J.** Vector network connecting the protomer NTD’, SD2, and SD1 domains. Domains are identified as Units splitting inter-protomer SD1/RBD to NTD/NTD’ pairs. **K.** Angular measures for the inter-protomer network. (left) SD2 to SD1 angles, (middle left) NTD’ to SD2 angles, (middle right) SD1 to NTD’ dihedrals, (right) NTD’ to SD2 dihedrals.

#### Vector analysis of the S-GSAS-B.1.1.7 S ectodomain

We next examined residue contact and domain position changes in the S-GSAS-B.1.1.7 S ectodomain relative to S-GSAS-D614G. Loss of the S2 S982-T547 latch hydrogen bond resulted in a ∼2 Å Cα-Cα shift in each protomer; visual inspection of each SD1 revealed the A570D containing loop occupies distinct positions (**Figure 4H-I**). This was reminiscent of u1S2q, an up-state stabilized construct we reported earlier, where a similar shift in the position of the A570 loop was implicated in increased up state propensity (Henderson et al., 2020). The marked asymmetry of the C1 reconstruction, in conjunction with the close coupling between SD1 and NTD′, suggested that these shifts might act to release constraints on the down state via reduced NTD to RBD coupling. Further, relative shifts in the orientation of a single SD1 should propagate throughout the trimer due to close contact between a SD1 loop (residues 557-569) contact with the adjacent protomer’s NTD′ loops (residues 38-45 and 281-284) as well as contact between RBDs. In order to examine the relative configurations of these domains, we determined angular dispositions between spatially paired SD1, NTD′ and SD2 domains via a vector network spanning the trimer (**Figure 4J-K**). Examination of the angle between the mobile RBD adjacent NTD′, and SD2 revealed a marked increase in angle relative to the other two protomers with a concomitant shift in the SD2 to SD1 angle. These angular changes were accompanied by a rotation of the NTD′ relative to the SD2 as well as a compensatory rotation of the distal protomer SD1 to NTD′ disposition. This compensatory shift explains the observed differences in the A570D loop positions. With the SD2 disposition relative to S2 largely similar to that of the other protomers, these movements can be ascribed to the S982A and A570D induced movements of SD1. Together, these changes results in disengagement of the NTD from the adjacent RBD, explaining the apparent increase in RBD mobility. This indicates the S982A and A570D pairing act as an allosteric switch through coupling of domain movements.

#### Summary

Taken together, our structural analysis of the S-GSAS-B.1.1.7 ectodomain highlights how allosteric effect of mutations in distal regions alter its RBD disposition. Moreover, the evolution of the B.1.1.7 variant appears to have balanced mutations that destabilizes the RBD-down or “closed” state to favor the RBD-up or “open” conformation, with other mutations that stabilize the pre-fusion spike conformation.

### The SARS-CoV-2 S protein B.1.351 and B.1.1.28 variants

The B.1.351 variant (20H/501Y.V2) harbors eight S protein mutations relative to the Wuhan-1 D614G mutant virus. All but one are located in the NTD and the RBD (**Figure 1A**). Accumulation of mutations in these immunodominant regions of the spike suggested selection pressure due to immune evasion. This is supported by reports that demonstrated resistance of the B.1.351 variant to RBD and NTD-directed antibodies, as well as to convalescent sera (Wibmer et al., 2021). B.1.351 has three mutations in the RBD - K417N, E484K and N501Y. These mutations also arose independently in Brazil in the B.1.1.28 variant. The B.1.1.28 variant subsequently acquired additional mutations in later variants, including the P.1 variant that was responsible for a resurgent COVID-19 outbreak in Manaus, Brazil (Sabino et al., 2021). We and others have shown that these three mutations have a strong deleterious effect on the activity of some RBD-directed antibodies. The E484K mutation is of particular concern, and has been shown to reduce or eliminate binding to many Class 1 RBD-directed antibodies, including antibody DH1041 when measured in an RBD only construct (Saunders et al., 2021; Wibmer et al., 2021). To study the effect of the B.1.351 mutations on the structure and the antigenicity of the S protein, we prepared an S ectodomain construct, S-GSAS-B.1.351, containing the eight mutations (**Figure 1A**). Additionally, we prepared an S ectodomain construct, S-GSAS-B.1.1.28, containing only the B.1.351 RBD mutations.

#### Receptor binding and antigenicity of the S-GSAS-B.1.351 and S-GSAS-B.1.1.28 ectodomains

We measured binding of the S-GSAS-B.1.351 and S-GSAS-B.1.1.28 constructs to the ACE2 receptor and to antibodies targeting different S epitopes by ELISA and SPR (**Figure 5**, **Supplemental Items 3-5 and Supplementary Table 1**). The ACE2 binding affinity to both variant S ectodomains improved relative to S-GSAS-D614G, an effect attributed to the N501Y mutation, also shared by the S-GSAS-B.1.1.7 variant (**Figures 3A and 5A**). The binding to the neutralizing NTD-directed antibody DH1050.1 was unaffected by the E484K mutation or by the triple RBD mutant S-GSAS-B.1.1.28, but was, however, impacted in the S-GSAS-B.1.351 construct that has multiple mutations in the NTD (**Figure 5A**, **Supplemental Item 2B)**. Although much reduced, DH1050.1 binding to S-GSAS-B.1.351 showed similar kinetics profile as S-GSAS-D614G (**Figure 5A and Supplemental Item 4)**.

**Figure 5.**
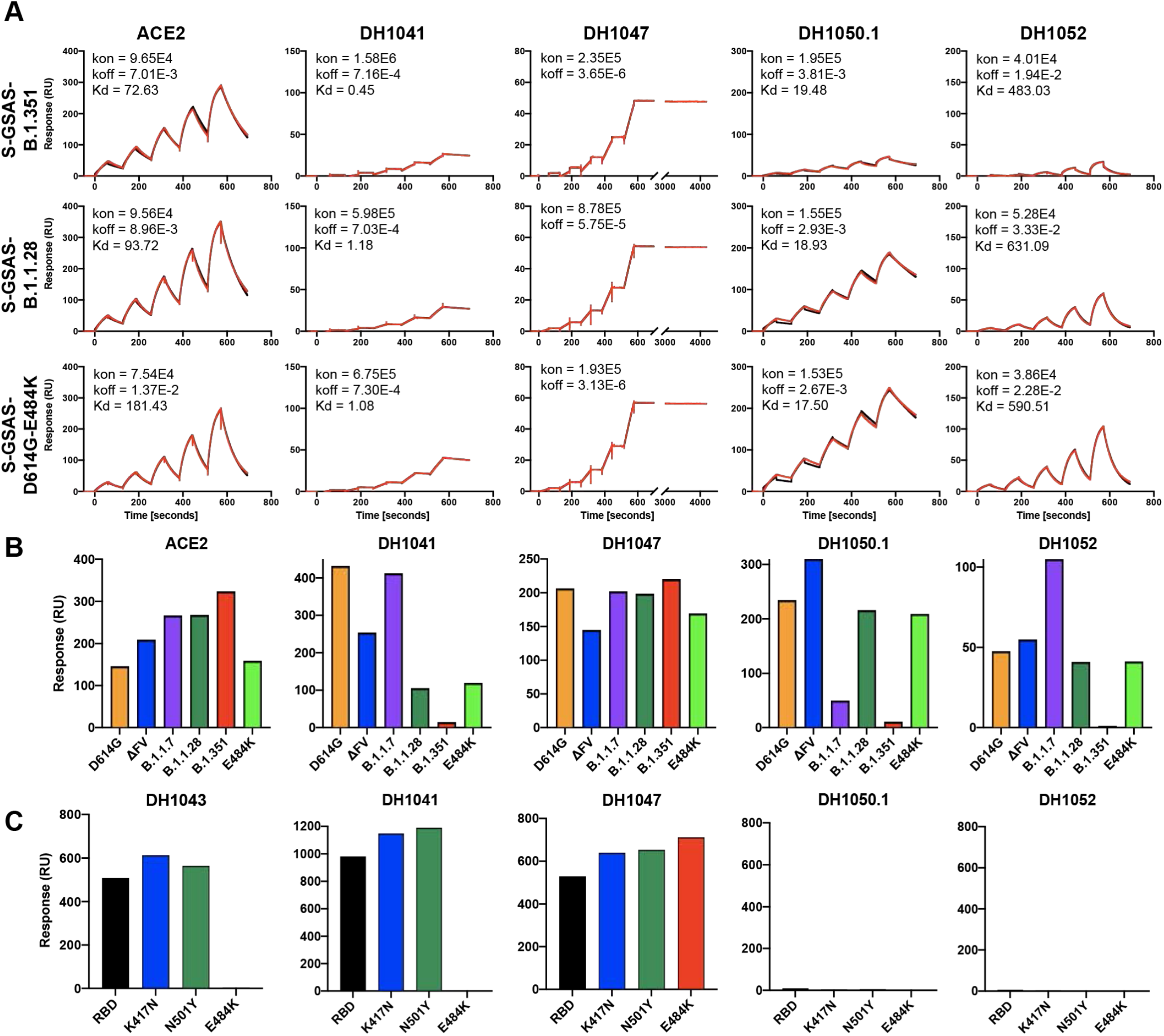
Antigenicity of of S-GSAS-B.1.351 and S-GSAS-B.1.1.28 ectodomains. **A.** Binding of ACE2 receptor ectodomain (RBD-directed), and antibodies DH1041 and DH1047 (RBD-directed, neutralizing), DH1050.1 (NTD-directed, neutralizing) and DH1052 (NTD-directed, non-neutralizing). to S-GSAS-B.1.351 (top), S-GSAS-B.1.1.28 (middle) and S-GSAS-E484K (bottom) measured by SPR using single-cycle kinetics. The red lines are the binding sensorgrams and the black lines show fits of the data to a 1:1 Langmuir binding model. The on-rate (k_on_, M^-1^s^-1^), off-rate (k_off_, s^-1^) and affinity (K_D_, nM) for each interaction are indicated in the insets. Binding of DH1047 was too tight for accurate measurement of K_D_. **B.** Binding of ACE2, RBD-directed antibodies DH1041 and DH1047, and NTD-directed antibodies DH1050.1 and DH1052 to spike variants, measured by SPR using the single injection format **C.** Binding of RBD-directed antibodies DH1041, DH1043 and DH1047, and NTD-directed antibodies DH1050.1 and DH1052 to WT RBD, RBD-K417N, RBD-N501Y and RBD-E484K, measured by SPR (see Supplemental Item 2).

The RBD-directed antibodies DH1041 and DH1043 bound robustly to the S-GSAS-D614G-E484K and S-GSAS-B.1.1.28 spikes by ELISA, although the E484K mutation is located at the epitope of these antibodies **(Supplemental Item 2 and 3)** (Li et al., 2021). By SPR, even though we observed decrease in the levels of binding of the DH1041 and DH1043 Fab to S-GSAS-D614G-E484K and S-GSAS-B.1.1.28 relative to S-GSAS-D614G, the binding profile was indicative of stable binding to the spikes (**Figure 5A and Supplemental Item 2B)**. The observation of robust and stable binding of the S-GSAS-D614G-E484K and S-GSAS-B.1.1.28 spikes to DH1041 and DH1043 despite the E484 residue being at the binding interface suggested accommodation of the E484K mutation. The binding of DH1041 and DH1043 to S-GSAS-B.1.351 was, however, dramatically reduced relative to S-GSAS-B.1.1.28, demonstrating an allosteric effect of the NTD mutations in S-GSAS-B.1.351 on the binding of antibodies DH1041 and DH1043. The binding data showing ability of antibodies DH1041 to accommodate the E484K mutation when binding to the spike, was is in contrast to its binding to an RBD-only construct, where the E484K mutation was shown to result in complete knockout of binding by ELISA (Saunders et al., 2021). We also measured dramatic loss of binding to antibodies DH1041 and DH1043 that target the RBD “hook” by SPR (**Figure 5B and Supplemental Item 13**). At the same time, binding to cross-reactive RBD-directed antibody DH1047 remained unaltered whether to the mutant spikes or to RBD-only constructs (**Figure 5C)** (Saunders et al., 2021). The binding to the fusion peptide-directed antibody DH1058 also remained unaffected by the spike mutations acquired by the variants **(Supplemental Items 2 and 4)**.

#### Cryo-EM structures of the S-GSAS-B.1.351 and S-GSAS-B.1.1.28 ectodomains

To visualize the impact of the mutations on the S protein conformation, we determined cryo-EM structures of S-GSAS-B.1.351 and S-GSAS-B.1.1.28 S ectodomains (**Figure 6**, **Supplementary Table 2, Supplemental Items 14-17**). We identified 3-RBD-down, 1-RBD-up and 2-RBD-up states in the S-GSAS-B.1.351 dataset, with total particle numbers of each state at 212,753, 1,047,277 and 200,301 respectively, thus resulting in ∼6:1 ratio of up to down state RBDs. The reduction in the fraction of closed or 3-RBD-down states is in contrast to the roughly 1:1 ratio reported for the S-GSAS-D614G ectodomain. A “consensus” 3-RBD-down state with 212,753 particles was refined to 3.7 Å, and displayed remarkably weak RBD density in one of the 3 RBDs that also appeared detached from its interprotomer contacting NTD (**Figure 6A**). Further classification of this consensus state yielded substates that showed similar features as the consensus structure, including a 65,713-particle state that we refined to 3.7 Å **(Supplemental Item 14)**. A 2-RBD-up population comprised of 200,301 particles was refined to overall resolution of 3.7 Å. Several populations of spike with one RBD in the “up” position were identified, which yielded typical 1-RBD-up structures with well-aligned S2 regions and variability in the S1 region due to NTD/RBD motion, and additional mobility of the “up” RBD (compared to a “down” RBD) as it oscillated between its interprotomer contacts with an NTD and an adjacent RBD (**Figure 6A and Supplemental Item 17**).

**Figure 6.**
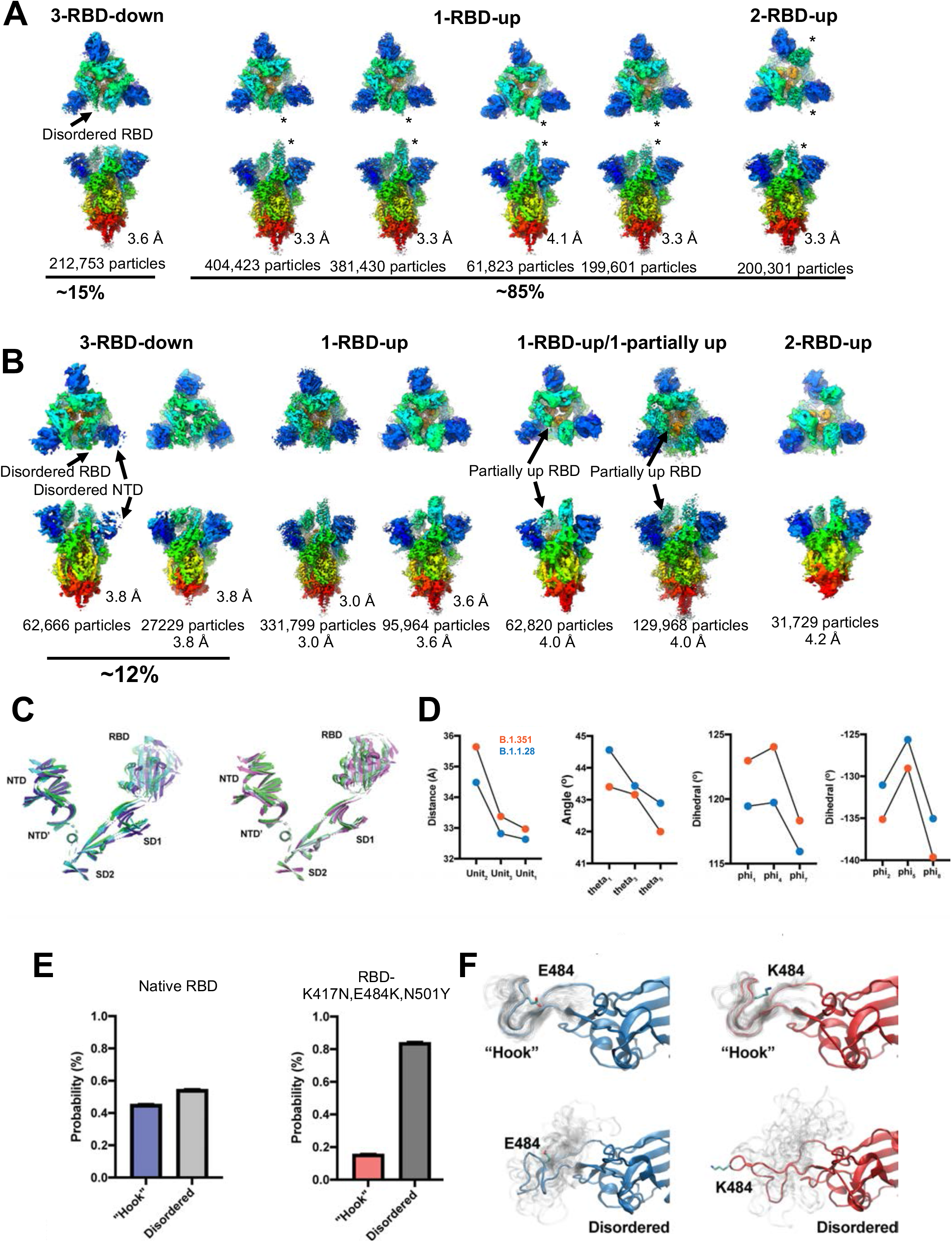
Structural analysis of the S-GSAS-B.1.1.28 and S-GSAS-B.1.351 ectodomains. Cryo-EM reconstructions of **A.** B.1.351. and **B.** B.1.1.28, in rainbow colors. **C.** Cartoon helix and sheet secondary structure elements of the (left) B.1.351 variant SD2 aligned S1 protomers. (right) B.1.1.28 variant SD2 aligned S1 protomers. **D.** Angle and dihedral measures for inter-protomer SD2-SD1-NTD’ network. (left) RBD to adjacent NTD distance, (middle left) NTD’ to SD2 angle, (middle right) SD1 to NTD’ dihedral, and (right) NTD’ to SD2 dihedral. **E.** State probabilities from (left) the WT RBD and (right) the B.1.1.28 variant RBD Markov model stationary distribution. Error bars indicate the 95% confidence interval. **F.** (top left) The WT “Hook” state of the RBD with 25 configurations shown in translucent grey. (bottom left)) The WT Disordered state of the RBD with 25 configurations shown in translucent grey. (top right) The WT “Hook” state of the RBD with 25 configurations shown in translucent grey. (bottom right) The WT Disordered state of the RBD with 25 configurations shown in translucent grey. Residue 484 is depicted in stick representation.

The S-GSAS-B.1.1.28 cryo-EM dataset mirrored the dramatic shift in the populations seen in the S-GSAS-B.1.351 dataset, with more heterogeneity and greater disorder in the smaller fraction of closed (or 3-RBD-down) structures and increase in the “up” or open populations. We identified multiple populations of “up” RBD states in the S-GSAS-B.1.1.28 dataset, including intermediate states that showed 1 RBD in the “up” position and another partially up. 3-RBD-down states accounted for ∼12% of the total spike population, and the cryo-EM reconstructions of these states showed considerable disorder in the RBDs. This disorder was more pronounced for one out of the three RBDs and its contacting NTD (**Figure 6B**). These observations implicate the 3 RBD mutations – K417N, E484K and N501Y – in disordering the RBDs in the 3-RBD-down state, and increasing the propensity to adopt an “up” configuration.

#### Vector analysis of the S-GSAS-B.1.351 and S-GSAS-B.1.1.28 ectodomain structures

Unlike the B.1.1.7 and mink-associated cluster 5 variants, the B.1.351 and B.1.1.28 variants include no S1 to S2 or interprotomer S1 mutations outside the RBD. The only mutation in the B.1.351 variant that is outside of the NTD and RBD regions is A701V that is located in the region between the SD2 and the fusion peptide, and is not involved in interdomain contacts. Visualization of the S1 interprotomer interfaces and alignment of each SD2 indicates each RBD and SD1 occupy distinct positions in both variants (**Figure 6C**). Unlike spikes harboring SD2 or SD1 mutations, the SD1 rotations in the B.1.351 and B.1.1.28 variants occur about an axis nearer the residue 570 loop, limiting this loop’s movement. These SD1 rotations result in large SD2 to SD1 angular differences and concomitant changes in the SD2 to SD1, SD1 to NTD′, NTD′ to SD2 dihedral dispositions with relatively muted differences in the NTD′ to SD2 angles (**Figure 6D**). Examination of the intra-protomer angles and dihedrals involving the RBD shows marked differences as well. In particular the protomer3 RBD orientation stands out resulting in its shift away from the adjacent NTD. Considering the location of the B.1.351 and B.1.1.28 mutations at the RBD-to-RBD interfaces and the lack of inter- and intra-protomer S1 mutations elsewhere, the observed differences in domain dispositions likely propagate through the RBD to SD1. Where the previously discussed variant structures appear to use SD2 or SD1 mediated NTD disengagement to facilitate enhanced RBD exposure, the B.1.351 and B.1.1.28 variants may instead utilize modification of the previously identified N343-glycan gate (Sztain et al., 2021) that contacts the adjacent RBD tip at a location near the E484K mutation to achieve a similar result.

#### Molecular dynamics simulations of RBD

The loss in binding of the RBD tip targeting DH1041 and DH1043 to the RBD-only construct harboring the K417N, E484K and N501Y mutations, was in contrast to their ability to bind the spike harboring the same three mutations in S-GSAS-B.1.1.28, where substantial binding was retained in ELISA and SPR assays (**Figure 5A-C**, **Supplemental Items 2, 4 and 15**). This suggested that disruption of intermolecular interactions between the spike and antibody by the E484K substitution is not solely responsible for the differences in interaction and that there might, therefore, be a conformational component to this observation. The cryo-EM structures of the down state spike of the B.1.1.28 variant displayed considerable disorder, especially at the RBD tip. Since the E484K substitution occurs at an interdomain interface where two adjacent RBDs contact via the N343-glycan of one with the RBD tip/hook region of the other (**Figure 3B**), we probed the conformational effect of the E484K substitution on the conformation of that region using molecular dynamics simulations. We built Markov state models of transitions between conformational states from large ensembles of short molecular dynamics simulations of both an unmutated RBD and a RBD harboring the K417N, E484K and N501Y mutations. (**Supplemental Items 18-20;** ∼260 µs total simulation time each). The Markov models for both RBD constructs were characterized by an RBD folded tip, “Hook” state, consistent with the conformation observed in binding mode 1 and 2 (Yuan et al., 2020) of RBD tip targeting Fab-RBD complex x-ray crystal structures, and a highly dynamic “Disordered” state in which the tip cycles between a variety of conformations (**Figures 6E-F**, **Supplemental Items 18 and 19, C and E**). While the native RBD displays a nearly even proportion of “Hook” vs. “Disordered” states, the triple mutant RBD spike shows a dramatic increase in the “Disordered” state population with a concomitant reduction in the “Hook” state (**Figure 6E**). These population differences are a consequence of an increased transition rate to the “Disordered” state from the “Hook” state in combination with a reduced transition rate back to the “Hook” state in the triple mutant compared to the native RBD (**Supplemental Items 18 and 19F**). Monitoring of the interactions between residue 484 sidechain in each model indicate the native E484 hydrogen bonds with the F490 backbone in particular, stabilizing the “Hook” state (**Supplemental Item 20**). Conversely, in the “Disordered” state, the K484 side chain forms fewer interactions across the RBD compared to E484, which primarily forms “Hook” region interactions (**Supplemental Item 20B**). Together, these results are consistent with a loss in DH1041 and DH1043 binding in the RBD-only context and indicate that the E484K mutation acts to destabilize the mode 1 and 2 SARS-CoV-2 neutralizing Mab-preferred conformation of the RBD tip. E484K mutation enhanced conformational disorder in the RBD “hook” may also be the source of the increased RBD up state population observed in both the B.1.351 and B.1.1.28 variants due to weakened RBD to RBD coupling. In the context of the spike, interprotomer interactions made by the RBD in its up state, as well as secondary contacts that the bound antibody makes to adjacent RBDs may play a role in stabilizing antibody binding to the E484K mutant **(Supplemental Figure 3)**.

### The SARS-CoV-2 S protein variant conformational comparison

Collectively, the structural results presented here indicate the primary consequence of 3-down state variant conformational adjustments is increased exposure of the RBD. A prime source of increased RBD up state propensity appears to be destabilization of one or more down RBDs in the 3-down state. In order to compare and contrast the disparate approaches toward destabilization, we generated an additional vector set describing RBD dispositions relative to adjacent RBDs and NTDs (**Figure 7**). Using an asymmetric refinement of the u1S2q design (Henderson et al., 2020) and four of our previously published 3-down state S-GSAS-D614G reconstructions (Gobeil et al., 2021), we first examined PCA clustering to identify structurally similar sets (**Figure 7A**). The B.1.351 and B.1.1.28 variants as well as the 3D-1 and 3D-3 S-GSAS-D614G-ΔFV structures were largely similar to the S-GSAS-D614G structures while the B.1.1.7 and 3D-2 S-GSAS-D614G-ΔFV structures cluster with the u1S2q structure. The S-GSAS-D614G-ΔFV 3D-4 structure differs markedly from all others and may correspond to a pre-protomer S1 dissociated (M1) state. The separation of S-GSAS-D614G-like and u1S2q-like sets is consistent with our structural analysis suggesting the B.1.351 and B.1.1.28 RBD destabilization is mediated by RBD-RBD contacts while the B.1.1.7 and D614G-ΔFV strategy utilizes modified SD1 or SD2 to S2 interaction for the same purpose. We next examined the primary vector contributors to differences observed in these clusters. Consistent with a role for the mobile RBD protomer, the three most variable measures are part of the mobile RBD protomer and included the angle between SD2 and SD1 as well as distances between S2 to SD2 and S2 to NTD′ (**Figure 7B, C**). Finally, the interconnected network of domain interactions suggests the protomer configurations should show correlations and that these might differ among the clusters (**Figure 7D**). We therefore examined angular correlations between SD2, SD1, and NTD′ orientations. As these orientations are label dependent, and since RBD mobility is closely tied to NTD proximity, and therefore to RBD up-state propensity, we assigned the RBD most distant from its adjacent NTD as the anchor for these comparisons. Examination of the correlation results indicated the Protomer3 and Unit1 angles and dihedrals in particular display significant correlated movements (**Figure 7E-F; Supplemental Items 21-25**). In the S-GSAS-D614G cluster structures, these together give rise to a shift in Unit3, drawing RBD3 away from the mobile RBD1 for which the N343-glycan bridging density is absent. The remaining RBD to RBD contact existed between the mobile RBD tip and adjacent RBD, with the B.1.351 and B.1.1.28 variant disordered tip presumably reducing the stability of this contact leading to their observed increase in up-state propensity. Correlations at these sites were often weaker in the u1S2q cluster which showed correlations primarily in the dihedral angles (**Supplemental Items 21-25**). These results show that the spike variants utilized the domain contact network to facilitate shifts in the RBD up state propensity by two different means. The analysis of domain contact differences demonstrates that disparate mutation sites with differing mechanistic impact on S protein conformation can effectively converge on similar increases in up-state RBDs.

**Figure 7.**
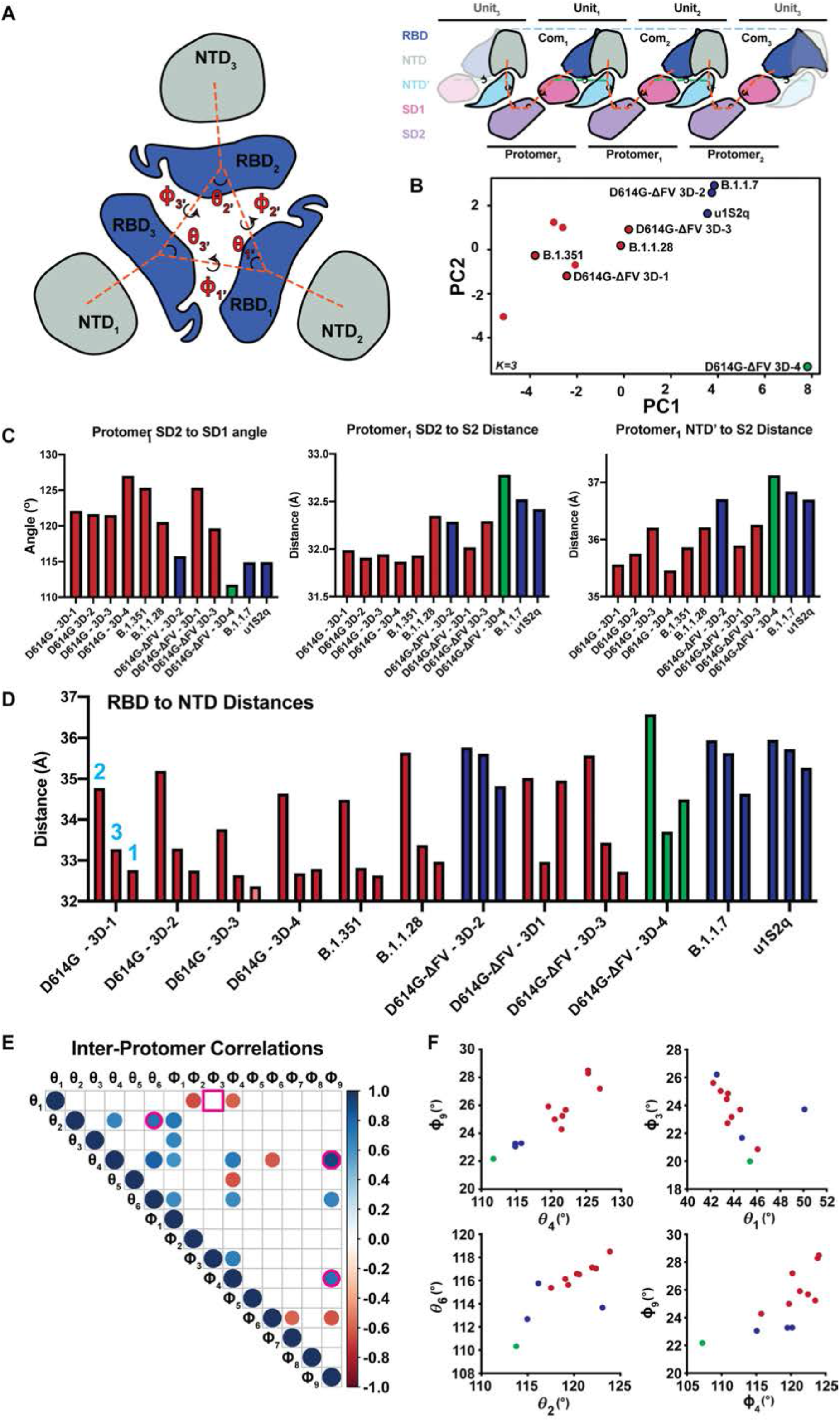
Comparison of inter-protomer network and RBD to RBD vector measures. **A.** (left) RBD and NTD vectors, angles, and dihedrals. (right) Simplified schematic of the SD2, SD1, NTD’ inter-protomer contact network. **B.** Principal components analysis of the inter-protomer network and RBD to RBD vector measures colored according to k-centers (K=3). Clusters correspond to a GSAS-D614G (D614G) like cluster (red), a u1S2q like cluster (blue), and outlier S-GSAS-D614G-ΔFV (D614G-ΔFV) 3D-4 (green). **C.** Top three contributors to PCA component one. **D.** RBD to NTD distance for the variants including D614G-ΔFV and the previously determined D614G. **E.** Significant Correlations between the inter-protomer angle measures (N=12, p<0.05). Pink outlines identify relationships plotted in panel (F). Square outline identifies non-significant correlation in the full structure set that was significant in the D614G cluster only correlations. **F.** Selected vector relationship plots. Colors are according to clusters in the PCA analysis in panel (B).

## Discussion

As multiple COVID-19 vaccines are being rolled out world-wide, the evolution of SARS-CoV-2 and acquisition of resistance to neutralizing antibodies is concerning as they may render current vaccines less effective, and, together with their increased transmissibility, threaten to prolong the pandemic. A different kind of threat is posed by interspecies transmission of SARS-CoV-2 from humans to other animals, with the case of transmission between mink and humans in mink farms. Establishment of a reservoir in a non-human host raises the possibility that the virus will evolve within the non-human host acquiring new mutations, and a transmission back to humans may cause these new variants to be resistant to vaccines and therapies. Since these reports came to light, there have been accelerated efforts to understand the effect of these mutations on viral fitness, transmissibility and resistance to antibodies elicited by current vaccines. Yet, a structural and mechanistic understanding of the effect of the S protein mutations in these variants is thus far lacking.

Here, we studied the S proteins of several SARS-CoV-2 variants in the context of a soluble S ectodomain. We have found similarities as well as fundamental differences between SARS-CoV-2 S protein evolution in the variant associated with inter-species transmission from the variants of interest that evolved in human hosts. All new variants studied showed increased binding to ACE2 receptor. For the variants that arose and evolved in humans, this is consistent with their increased transmissibility. For the mink-associated variant, the increased affinity of the S protein for the ACE2 receptor may play a role in engaging the homologous receptor in minks, helping to establish infection in the new host. While all human-evolved variants studied here showed reduced binding to antibodies at dominant neutralization epitopes, the mink-associated variant retained similar levels of binding to all antibodies tested. Pressure to adapt to a new host may render the virus less fit in the natural host. Consistent with this possibility, we observed evidence for mink-associated variant S protein instability, including identification of a spike population that was partially unfolded. These observations may provide insights into why the mink-associated variant, although it was transmitted back to humans, did not spread very widely.

For the human-evolved variants of interest, we found that the S protein utilized different mechanisms for manipulation of its immunodominant regions, namely the NTD and RBD, and to converge on a common goal of destabilizing the 3-RBD-down state. While in the B.1.1.7 variant this occurred by modifications in SD1 or SD2 to S2 interaction, for the B.1.351 and B.1.1.28 variants RBD destabilization was mediated by RBD-RBD contacts. Indeed, we found intriguing similarities between mutations we had previously engineered to manipulate the S protein (Henderson et al., 2020) with mutations that occur naturally. The CoV spike protein utilizes a network of S1 subunit domain interactions to control the functionally critical disposition of the RBD. The extensive interconnectivity and inherent metastability of the spike render the entire molecular configuration susceptible to relatively minor modifications to domain pairing strength. Our results here show that SARS-CoV-2 variants have taken advantage of this mechanical feature to modify the structural state of the spike.

In summary, our studies provide a structural and mechanistic understanding of the impact of mutations that naturally evolved during the course of a pandemic on the S protein conformation, and in doing so, provides a framework for understanding their functional impact. We demonstrate that convergent S protein evolution to increase SARS-CoV-2 transmissibility and escape neutralization at immunodominant NTD and RBD epitopes can occur via different allosteric communication networks in the spike.

## Acknowledgements

Cryo-EM data were collected at the National Center for Cryo-EM Access and Training (NCCAT) and the Simons Electron Microscopy Center located at the New York Structural Biology Center, supported by the NIH Common Fund Transformative High Resolution Cryo-Electron Microscopy program (U24 GM129539) and by grants from the Simons Foundation (SF349247) and NY State. We thank Ed Eng, Mahira Aragon, Eugene Chua and Joshua Mendez for microscope alignments and assistance with cryo-EM data collection. This work was supported by an administrative supplement to NIH R01 AI145687 for coronavirus research (P.A. and R.C.H.) and NC State funding for COVID research (B.F.H). This study utilized the computational resources offered by Duke Research Computing (http://rc.duke.edu; NIH 1S10OD018164-01) at Duke University. We thank C. Kneifel, M. Newton, V. Orlikowski, T. Milledge, and D. Lane from the Duke Office of Information Technology and Research Computing for assisting with setting up and maintaining the computing environment.

## Author contributions

S.M-C.G. and P.A. designed and led the study, and determined and analyzed cryo-EM structures. S.M-C.G. designed SARS-CoV-2 ectodomain constructs, expressed and purified proteins, and performed SPR assays. K.J., V.S. and M.K. expressed and purified proteins. S.M., K.M. and R.C.H performed structural analysis. K.Mansouri and R.J.E. performed NSEM analysis. R.P. performed ELISA assays. K.O.S. provided key reagents. B.F.H supervised ELISA assays. S.M-C.G., P.A. and R.C.H wrote the manuscript with help from all authors. R.C.H led computational analysis. P.A. supervised the study and reviewed all data.

## STAR METHODS

### RESOURCE AVAILABILITY

#### Lead Contact

Further information and requests for resources and reagents should be directed to and will be fulfilled by the Lead Contact, Priyamvada Acharya.

#### Materials Availability

Further information and requests for resources and reagents should be directed to Priyamvada Acharya. Plasmids generated in this study have will be deposited to Addgene.

#### Data and Code Availability

Cryo-EM reconstructions and atomic models generated during this study are available at wwPDB and EMBD (https://www.rcsb.org; http://emsearch.rutgers.edu) under the accession codes PDB IDs 7LWI, 7LWJ, 7LWK, 7LWL, 7LWM, 7LWN, 7LWO, 7LWP, 7LWQ, 7LWT, 7LWU, 7LWV, 7LWS, 7LWW, 7LYK, 7LYL, 7LYM, 7LYN, 7LYO, 7LYP and 7LYQ and EMDB IDs EMDB-23546, EMD-23547, EMD-23548, EMD-23549, EMD-23550, EMD-23551, EMD-23552, EMD-23553, EMD-23554, EMD-23556, EMD-23557, EMD-23558, EMD-23555, EMD-23559, EMD-23593, EMD-23594, EMD-23595, EMD-23596, EMD-23597, EMD-23598 and EMD-23599. Vector analysis and Markov modelling scripts are available at: https://gitlab.cs.duke.edu/henderson_lab/variant_mar2021 along with information for downloading filtered molecular simulation trajectories.

### EXPERIMENTAL MODEL AND SUBJECT DETAILS

#### Cell culture

Gibco FreeStyle 293-F cells (embryonal, human kidney) were incubated at 37°C and 9% CO2 in a humidified atmosphere. Cells were maintained in FreeStyle 293 Expression Medium (Gibco) with agitation at 120 rpm and 75% humidity. Plasmids were transiently transfected into cells using Turbo293 (SpeedBiosystems) and incubated at 37°C, 9% CO2, 120 rpm for 6 days. On the day following transfection, HyClone CDM4HEK293 media (Cytiva, MA) was added to the cells. Antibodies were produced in Expi293 cells (embryonal, human kidney). Cells were maintained in Expi293 Expression Medium at 37°C, 120 rpm and 8% CO2, 75% humidity. Plasmids were transiently transfected using the ExpiFectamine 293 Transfection Kit and protocol (Gibco).

### METHOD DETAILS

#### Plasmids

Gene synthesis for all plasmids generated by this study were performed and the sequence confirmed by GeneImmune Biotechnology (Rockville, MD). The SARS-CoV-2 spike protein ectodomain constructs comprised the S protein residues 1 to 1208 (GenBank: MN908947) with the D614G mutation, the furin cleavage site (RRAR; residue 682-685) mutated to GSAS, a C-terminal T4 fibritin trimerization motif, a C-terminal HRV3C protease cleavage site, a TwinStrepTag and an 8XHisTag. All spike ectodomains were cloned into the mammalian expression vector pαH (Wrapp et al., 2020). For the ACE2 construct, the C-terminus was fused a human Fc region.

#### Protein purification

On the 6^th^ day post transfection, spike ectodomains were harvested from the concentrated supernatant. The spike ectodomains were purified using StrepTactin resin (IBA) and size exclusion chromatography (SEC) using a Superose 6 10/300 GL Increase column equilibrated in 2mM Tris, pH 8.0, 200 mM NaCl, 0.02% NaN3. All steps of the purification were performed at room temperature and in a single day. The purified proteins were flash frozen and stored at −80 °C in single-use aliquots. Each aliquot were thawed by a 20-minute incubation at 37 °C before use. Antibodies were produced in Expi293F cells and purified by Protein A affinity and digested to their Fab state using LysC. ACE2 with human Fc tag was purified by Protein A affinity chromatography and SEC.

#### SPR

Antibody binding to SARS-CoV-2 spike and RBD constructs was assessed using SPR on a Biacore T-200 (Cytiva, formerly GE Healthcare) with HBS buffer supplemented with 3 mM EDTA and 0.05% surfactant P-20 (HBS-EP+). All binding assays were performed at 25 °C. Spike variants were captured on a Series S Strepavidin (SA) chip coated at 200 nM (60s at 10µL/min). The antibodies Fabs were injected at concentrations ranging from 0.625 nM to 800 nM (prepared in a 2-fold serial dilution manner) over the S proteins using the single cycle kinetics mode with 5 concentration per cycle. For the single injection assay, the Fabs were used at a concentration of 200nM. A contact time of 60 seconds and a dissociation time of 120 seconds (3600 seconds for DH1047) at a flow rate of 50µL/min was used. The surface was regenerated after the last injection with 3 pulses of a 50mM NaoH + 1M NaCl solution for 10 seconds at 100µL/min. For the RBDs, the antibodies were captured on a CM5 chip coated with HumanAntiFc (Cytiva) by their FC region at 100nM using a flowrate of 5µL/min for 120s. The RBDs were then injected at 100nM for 120s at a flowrate of 50µL/min with a dissociation time of 30s. The surface was regenerated by 3 consecutive pulse of 3M MgCl2 for 10s at 100µL/min. Sensogram data were analyzed using the BiaEvaluation software (Cytiva)

#### Negative-stain electron microscopy

Samples were diluted to 100 µg/ml in 20 mM HEPES pH 7.4, 150 mM NaCl, 5% glycerol, 7.5 mM glutaraldehyde and incubated for 5 minutes before quenching the glutaraldehyde by the addition of 1 M Tris (to a final concentration of 75 mM) and 5 minutes incubation. A 5-µl drop of sample was applied to a glow-discharged carbon-coated grid for 10-15 seconds, blotted, stained with 2% uranyl formate, blotted and air-dried. Images were obtained using a Philips EM420 electron microscope at 120 kV, 82,000× magnification, and a 4.02 Å pixel size. The RELION (Scheres, 2012) software was used for particle picking, and 2D and 3D class averaging.

#### ELISA assays

Spike ectodomains tested for antibody- or ACE2-binding in ELISA assays as previously described (Edwards et al., 2020). Assays were run in two formats i.e. antibodies/ACE2 coated or spike coated. For the first format, the assay was performed on 384-well plates coated at 2 µg/ml overnight at 4^°^C, washed, blocked and followed by two-fold serially diluted spike protein starting at 25 µg/mL. Binding was detected with polyclonal anti-SARS-CoV-2 spike rabbit serum (developed in our lab), followed by goat anti-rabbit-HRP and TMB substrate. Absorbance was read at 450 nm. In the second format, serially diluted spike protein was bound in wells of a 384-well plates, which were previously coated with streptavidin at 2 µg/mL and blocked. Proteins were incubated at room temperature for 1 hour, washed, then human mAbs were added at 10 µg/ml. Antibodies were incubated at room temperature for 1 hour, washed and binding detected with goat anti-human-HRP and TMB substrate.

#### Cryo-EM

Purified SARS-CoV-2 spike ectodomains were diluted to a concentration of ∼1.5 mg/mL in 2 mM Tris pH 8.0, 200 mM NaCl and 0.02% NaN3 and 0.5% glycerol was added. A 2.3-µL drop of protein was deposited on a Quantifoil-1.2/1.3 grid that had been glow discharged for 10 seconds using a PELCO easiGlow™ Glow Discharge Cleaning System. After a 30 seconds incubation in >95% humidity, excess protein was blotted away for 2.5 seconds before being plunge frozen into liquid ethane using a Leica EM GP2 plunge freezer (Leica Microsystems). Frozen grids were imaged using a Titan Krios (Thermo Fisher) equipped with a K3 detector (Gatan).

#### Vector Based Structure Analysis

Vector analysis of intra-protomer domain positions was performed as described previously (Henderson et al., 2020) using the Visual Molecular Dynamics (VMD)(Humphrey et al., 1996) software package Tcl interface. For each protomer of each structure, Cα centroids were determined for the NTD (residues 27 to 69, 80 to 130, 168 to 172, 187 to 209, 216 to 242, and 263 to 271), NTD’ (residues 44 to 53 and 272 to 293), RBD (residues 334 to 378, 389 to 443, and 503 to 521), SD1 (residues 323 to 329 and 529 to 590), SD2 (residues 294 to 322, 591 to 620, 641 to 691, and 692 to 696), CD (residues 711 to 716 1072 to 1121), and a S2 sheet motif (S2s; residues 717 to 727 and 1047 to 1071). Additional centroids for the NTD (NTDc; residues 116 to 129 and 169 to 172) and RBD (RBDc; residues 403 to 410) were determined for use as reference points for monitoring the relative NTD and RBD orientations to the NTD’ and SD1, respectively. Vectors were calculated between the following within protomer centroids: NTD to NTD’, NTD’ to SD2, SD2 to SD1, SD2 to CD, SD1 to RBD, CD to S2s, NTDc to NTD, RBD to RBDc. Vector magnitudes, angles, and dihedrals were determined from these vectors and centroids. Inter-protomer domain vector calculations for the SD2, SD1, and NTD’ used these centroids in addition to anchor residue Cα positions for each domain including SD2 residue 671 (SD2r), SD1 residue 575 (SD1r), and NTD’ residue 276 (NTD’r). These were selected based upon visualization of position variation in all protomers used in this analysis via alignment of all of each domain in PyMol(Schrodinger, 2015). Vectors were calculated for the following: NTD’ to NTD’r, NTD’ to SD2, SD2 to SD2r, SD2 to SD1, SD1 to SD1r, and SD1 to NTD’. Angles and dihedrals were determined from these vectors and centroids. Vectors for the RBD to adjacent RBD and RBD to adjacent NTD were calculated using the above RBD, NTD, and RBDc centroids. Vectors were calculated for the following: RBD2 to RBD1, RBD3 to RBD2, and RBD3 to RBD1. Angles and dihedrals were determined from these vectors and centroids. Principal components analysis, K-means clustering, and Pearson correlation (confidence interval 0.95, p<0.05) analysis of vectors sets was performed in R(Team, 2017). Data were centered and scaled for the PCA analyses.

#### Adaptive Sampling Molecular Dynamics

The CHARMM CR3022 bound SARS-CoV-2 RBD crystal structure(Nichols et al., 2020) (PDB ID 6ZLR) model(Jo et al., 2008; Woo et al., 2020) was used for the adaptive sampling simulations. The CR3022 antibody, glycan unit, water, and ions were stripped from the model leaving only the protein portion of the RBD. The final model comprised Spike residues 327 to 529. A single Man5 glycan was added at the N343 position using the CHARMM GUI(Jo et al., 2008) with the B.1.1.28/B.1.351 RBD mutations K417N, E484K, and N501Y prepared in PyMol. Systems for simulation were built using the AmberTools20 Leap(D.A. Case, 2020) program. The unmutated (WT) and B.1.1.28/B.1.351 (Mut) RBDs were immersed in a truncated octahedral TIP3P water box with a minimum edge distance of 15 Å to the nearest protein atom followed by system neutralization with chlorine atoms resulting in systems sizes of 67,508 and 66,894 atoms for the WT and Mut, respectively. The Amber ff14SB protein(Maier et al., 2015) and Glycam(Kirschner et al., 2008) forefields were used throughout. All simulations were performed using the Amber20 pmemd CUDA implementation. The systems were first minimized for 10,000 steps with protein atom restraints followed by minimization of the full system without restraints for an additional 10,000 steps. This was followed by heating of the systems from 0 K to 298 K over a period of 20 ps in the NVT ensemble using a 2 fs timestep using the particle mesh Ewald method for long-range electrostatics and periodic boundary conditions(Essmann et al., 1995). The systems were then equilibrated for 100 ps in the NPT ensemble with the temperature controlled using Langevin dynamics with a frequency of 1.0 ps^-1^ and 1 atm pressure maintained using isotropic position scaling with a relaxation time of 2 ps (Loncharich et al., 1992). A non-bonded cut-off of 8 Å was used throughout and hydrogen atoms were constrained using the SHAKE algorithm (Ryckaert et al., 1977) with hydrogen mass repartitioning (Hopkins et al., 2015) used to allow for a 4 fs timestep. In order to generate an ensemble of RBD tip conformations for initiation of the adaptive sampling routine, we performed one hundred 50 ns simulations in the NVT ensemble with randomized initial velocities for each of the WT and Mut systems. The final frame from each of these simulations was used to initiate the adaptive sampling scheme. Adaptive sampling was performed using the High-Throughput Molecular Dynamics (HTMD v. 1.24.2) package (Doerr et al., 2016). Each iteration consisted of 50-100 independent simulations of 100 ns. Simulations from each iteration were first projected using a dihedral metric with angles split into their sin and cos components for residues 454 to 491. This was followed by a TICA (Pérez-Hernández et al., 2013) projection using a lag time of 5 ns and retaining five dimensions. Markov state models were then built using a lag time of 50 ns for the selection of new states for the next iteration. A total of 29 adaptive iterations were performed yielding total simulation times of 274.8 and 256.8 µs for the WT and Mut systems, respectively. Simulations were visualized in VMD and PyMol.

#### Markov State Modelling

Markov state models (MSMs) were prepared in HTMD with an appropriate coordinate projection selected using PyEMMA (Scherer et al., 2015) (v. 2.5.7). Multiple projections were tested on a 25 µs subset of the Mut simulations that included atomic distance and contact measures between RBD residues as well as backbone torsions of the RBD tip residues using the variational approach to Markov processes score (Wu and Noé, 2020) (Supplementary Table 3). This led to the selection of a Cα pairwise distance metric between residues 471 to 480 and 484 to 488 for MSM construction. MSMs were prepared in HTMD using a TICA lag time of 5 ns retaining five dimensions followed by K-means clustering using 500 cluster centers. The implied timescales (ITS) plots were used to select a lag time of 30 ns for MSM building. Models were coarse-grained via Perron cluster cluster analysis (PCCA++) using 2 states and validated using the Chapman-Kolmogorov (CK) test. A bootstrapping routine without replacement was used to calculate measurement errors retaining 80% of the data per iteration for a total of 100 iterations. State statistics were collected for mean first passage times (MFPT), stationary distributions, and root-mean square deviations (RMSD) for RBD tip residues 470-490. Residue 484 sidechain contacts were calculated from a representative model. A contact was defined as atom pairing within 3.5 Å between either the minimum of either E484 γ-carboxyl O atoms (for WT) or K484 ε-amino N atom (for Mut) and backbone or sidechain O or N atoms for residues 348 to 354, 413 to 425, or 446 to 500. The RMSD and contact metric means were model weighted. Weighted state ensembles containing 250 structures were collected for visualization in VMD.

### QUANTIFICATION AND STATISTICAL ANALYSIS

Principal components analysis, K-means clustering, and Pearson correlation (confidence interval 0.95, p<0.05) analysis of vectors sets was performed in R. Data were centered and scaled for the PCA analyses.

### Key Resources Table

| REAGENT or RESOURCE | SOURCE | IDENTIFIER |
| --- | --- | --- |
| **Antibodies** |  |  |
| ACE2 | (Henderson et al., 2020) | N/A |
| CR3022 | (Henderson et al., 2020) | N/A |
| 2G12 | (Acharya et al., 2020) | N/A |
| DH1041 | (Li et al., 2021) | N/A |
| DH1043 | (Li et al., 2021) | N/A |
| DH1047 | (Li et al., 2021) | N/A |
| DH1050.1 | (Li et al., 2021) | N/A |
| DH1050.2 | (Li et al., 2021) | N/A |
| DH1052 | (Li et al., 2021) | N/A |
| DH1058 | (Li et al., 2021) | N/A |
| Ab82 |  | N/A |
| Goat anti-rabbit-HRP | Abcam | ab97080 |
| Goat anti-human-HRP | Jackson ImmunoResearch Laboratories | 109-035-098 |
| **Bacterial and Virus Strains** |  |  |
| N/A |  |  |
| **Biological Samples** |  |  |
| N/A |  |  |
| **Chemicals, Peptides, and Recombinant Proteins** |  |  |
| RBD | (Saunders et al., 2021) | N/A |
| RBD-K417N | (Saunders et al., 2021) | N/A |
| RBD-N501Y | (Saunders et al., 2021) | N/A |
| RBD-E484K | (Saunders et al., 2021) | N/A |
| FreeStyle 293 Expression Medium | GIBCO | 12338018 |
| Expi293 Expression Medium | GIBCO | A1435101 |
| ExpiFectamine 293 Transfection Kit | GIBCO | A14524 |
| Hyclone SFM4HEK293 | Cytiva | SH30521.02 |
| Opti-MEM I | GIBCO | 31985-070 |
| Turbo293 | Speed BioSystems | PXX1002 |
| 10x Buffer E | IBA | 2-1000 |
| 10x Buffer R | IBA | 2-1002 |
| 10x Buffer W | IBA | 2-1003 |
| Strep-Tactin resin | IBA | 2-1201 |
| 8% Glutaraldehyde | Electron Microscopy Sciences | 16019 |
| 300mesh Cu carbon coated | Electron Microscopy Sciences | CF300-Cu |
| Uranyl formate | Electron Microscopy Sciences | 22450 S-888 |
| Quantifoil R 1.2/1.3 300 mesh | Electron Microscopy Sciences | Q350CR-14 |
| Streptavidin | Thermo Fisher Scientific | S-888 |
| **Critical Commercial Assays** |  |  |
| Superose 6 Increase 10/300 GL | Cytiva | 29091596 |
| NuPage 4-12% | Invitrogen | NP0321 |
| TMB substrate | Sera Care Life Sciences | 5120-0083 |
| Sensor Chip CM5 | Cytiva | 29149603 |
| Sensor Chip SA | Cytiva | BR100531 |
| HBS-EP+ 10x | Cytiva | BR100669 |
| Human Antibody Capture Kit | Cytiva | BR100839 |
| **Deposited Data** |  |  |
| Mink Cluster 5-associated SARS-CoV-2 spike protein (S-GSAS-D614G-delFV) in the 3-RBD down conformation | This paper | PDB 7LWI, EMD-23546 |
| Mink Cluster 5-associated SARS-CoV-2 spike protein (S-GSAS-D614G-delFV) in the 3-RBD down conformation | This paper | PDB 7LWJ, EMD-23547 |
| Mink Cluster 5-associated SARS-CoV-2 spike protein (S-GSAS-D614G-delFV) in the 3-RBD down conformation | This paper | PDB 7LWK, EMD-23548 |
| Mink Cluster 5-associated SARS-CoV-2 spike protein (S-GSAS-D614G-delFV) in the 3-RBD down conformation | This paper | PDB 7LWL, EMD-23549 |
| Mink Cluster 5-associated SARS-CoV-2 spike protein (S-GSAS-D614G-delFV) in the 1-RBD up conformation | This paper | PDB 7LWM, EMD-23550 |
| Mink Cluster 5-associated SARS-CoV-2 spike protein (S-GSAS-D614G-delFV) in the 1-RBD up conformation | This paper | PDB 7LWN, EMD-23551 |
| Mink Cluster 5-associated SARS-CoV-2 spike protein (S-GSAS-D614G-delFV) in the 1-RBD up conformation | This paper | PDB 7LWO, EMD-23552 |
| Mink Cluster 5-associated SARS-CoV-2 spike protein (S-GSAS-D614G-delFV) in the 2-RBD up conformation | This paper | PDB 7LWP, EMD-23553 |
| Mink Cluster 5-associated SARS-CoV-2 spike protein(S-GSAS-D614G-delFV) missing the S1 subunit and SD2 subdomain of one protomer | This paper | PDB 7LWQ, EMD-23554 |
| UK (B.1.1.7) SARS-CoV-2 S-GSAS-D614G variant spike protein in the 3-RBD-down conformation | This paper | PDB 7LWS, EMD-23555 |
| UK (B.1.1.7) SARS-CoV-2 spike protein variant (S-GSAS-B.1.1.7) in the 1-RBD-up conformation | This paper | PDB 7LWT, EMD-23556 |
| UK (B.1.1.7) SARS-CoV-2 spike protein variant (S-GSAS-B.1.1.7) in the 1-RBD-up conformation | This paper | PDB 7LWU, EMD-23557 |
| UK (B.1.1.7) SARS-CoV-2 spike protein variant (S-GSAS-B.1.1.7) in the 1-RBD-up conformation | This paper | PDB 7LWV, EMD-23558 |
| Brazilian (B.1.1.28) SARS-CoV-2 spike protein variant (S-GSAS-B.1.1.28) in the 1-RBD-up conformation | This paper | PDB 7LWW, EMD-23559 |
| South African (B.1.351) SARS-CoV-2 spike protein variant (S-GSAS-B.1.351) in the RBD-down conformation | This paper | PDB 7LYL, EMD-23594 |
| South African (B.1.351) SARS-CoV-2 spike protein variant (S-GSAS-B.1.351) in the RBD-down conformation | This paper | PDB 7LYM, EMD-23595 |
| South African (B.1.351) SARS-CoV-2 spike protein variant (S-GSAS-B.1.351) in the 2-RBD-up conformation | This paper | PDB 7LYK, EMD-23593 |
| South African (B.1.351) SARS-CoV-2 spike protein variant (S-GSAS-B.1.351) in the 1-RBD-up conformation | This paper | PDB 7LYN, EMD-23596 |
| South African (B.1.351) SARS-CoV-2 spike protein variant (S-GSAS-B.1.351) in the 1-RBD-up conformation | This paper | PDB 7LYO, EMD-23597 |
| South African (B.1.351) SARS-CoV-2 spike protein variant (S-GSAS-B.1.351) in the 1-RBD-up conformation | This paper | PDB 7LYP, EMD-23598 |
| South African (B.1.351) SARS-CoV-2 spike protein variant (S-GSAS-B.1.351) in the 1-RBD-up conformation | This paper | PDB 7LYQ, EMD-23599 |
| **Experimental Models: Cell Lines** |  |  |
| Freestyle 293-F cells | GIBCO | R79007 |
| Expi293F cells | GIBCO | A14527 |
| **Experimental Models: Organisms/Strains** |  |  |
| N/A |  |  |
| **Oligonucleotides** |  |  |
| N/A |  |  |
| **Recombinant DNA** |  |  |
| paH-S-GSAS | (Gobeil et al., 2021) | Addgene 164565 |
| paH-S-GSAS-D614G | (Gobeil et al., 2021) | Addgene 164566 |
| paH-S-GSAS-D614G-ΔFV | This paper | Addgene XXXXX |
| paH-S-GSAS-H69/V70del | This paper | Addgene XXXXX |
| paH-S-GSAS-D614G-Y453F | This paper | Addgene XXXXX |
| paH-S-GSAS-D614G-I692V | This paper | Addgene XXXXX |
| paH-S-GSAS-B.1.1.7 | This paper | Addgene XXXXX |
| paH-S-GSAS-D614G-N501Y | This paper | Addgene XXXXX |
| paH-S-GSAS-B.1.351 | This paper | Addgene XXXXX |
| paH-S-GSAS-B.1.1.28 | This paper | Addgene XXXXX |
| paH-S-GSAS-D614G-E484K | This paper | Addgene XXXXX |
| paH-S-GSAS-D614G-K417N | This paper | Addgene XXXXX |
| **Software and Algorithms** |  |  |
| Relion | (Scheres, 2012; Scheres, 2016) | Version 3.1 |
| cryoSPARC | (Punjani et al., 2017) | https://cryosparc.com |
| Phenix | (Afonine et al., 2018; Liebschner et al., 2019) | Version 1.17 |
| Coot | (Emsley et al., 2010) | Version 0.8.9.2 |
| Pymol | Schrodinger<br>The PyMOL Molecular Graphics System (Schrödinger, 2015). | https://www.pymol.org/ |
| UCSF Chimera | (Pettersen et al., 2004) | http://www.cgl.ucsf.edu/chimera/ |
| Chimera X | (Goddard et al., 2018) | https://www.rbvi.ucsf.edu/chimerax/ |
| Image Lab | Bio-Rad | Version 6.0 |
| PRISM 8 | GraphPad Software | Version 8.4.0 |
| R | R Core Team (2014). R: A language and environment for statistical computing. R Foundation for Statistical Computing, Vienna, Austria. URL http://www.R-project.org/ | version 4.0.2 |
| Bio3D | (Grant et al., 2020) | version 2.4-1 |
| ImageJ | (Schneider et al., 2012) | Version 1.53a |
| VMD | (Humphrey et al., 1996) | Version 1.9.4 |
| HTMD | (Doerr et al., 2016) | Version 1.24.2 |
| PyEMMA | (Scherer et al., 2015) | Version 2.5.7 |
| Amber | (D.A. Case, 2020) | Version 20 |
| **Other** |  |  |
| N/A |  |  |

