## Supplemental Data for "Effect of natural mutations of SARS-CoV-2 on spike structure, conformation and antigenicity"

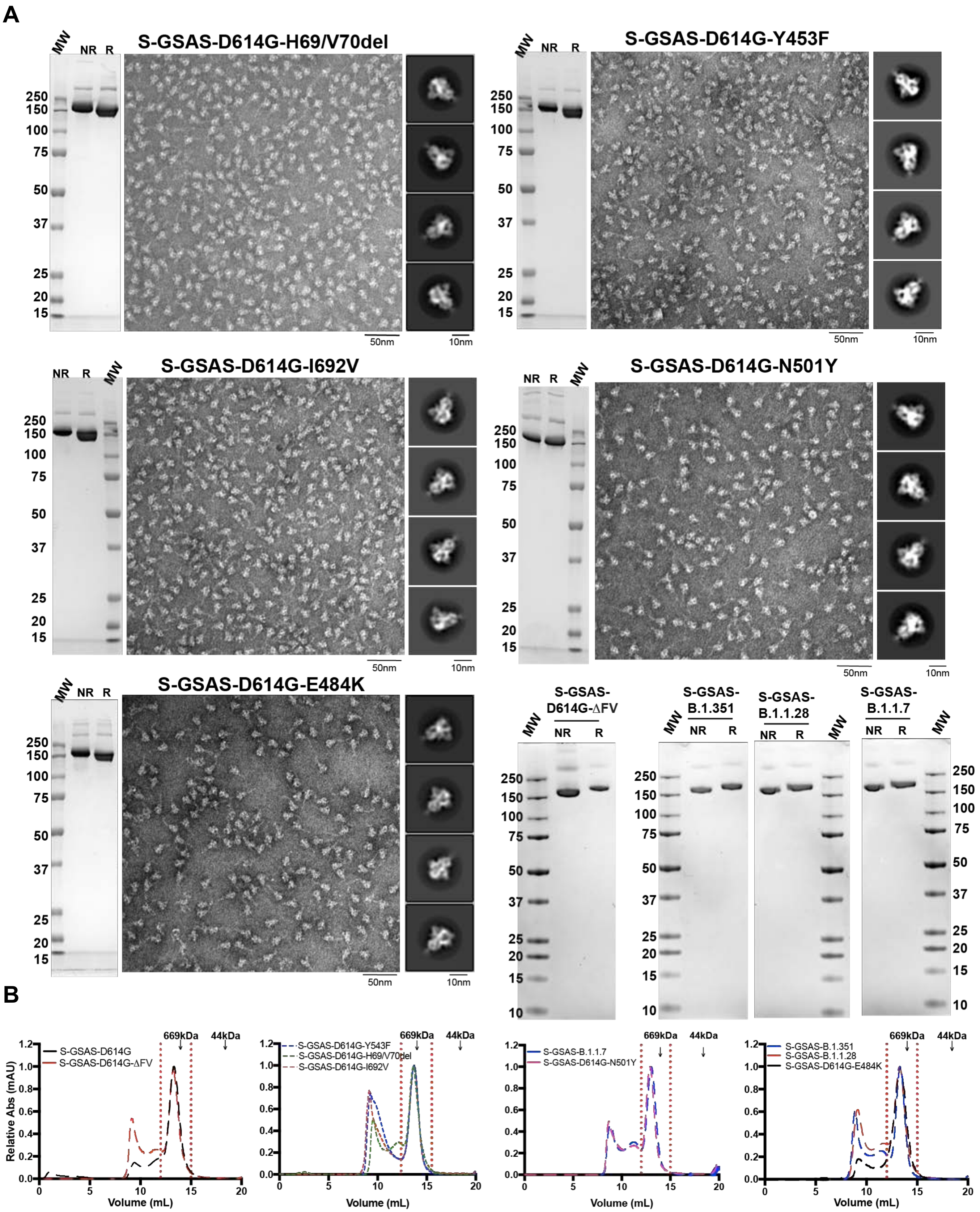


**Supplemental Item 1. Quality control assessment of the spike variants preparations, Related to Figures 1. A**. SDS-PAGE of the SEC purified ectodomains and, representative NSEM micrograph and 2D class averages. **B**. Size-exclusion chromatography (SEC) elution profile on a superose 6 10/300 column of spike variants ectodomains. Fractions isolated for further characterization are indicated by vertical red dotted lines. Elution volumes of molecular weight standards at 669 (thyroglobulin) and 44 kDa (ovalbumin) are labelled for reference.

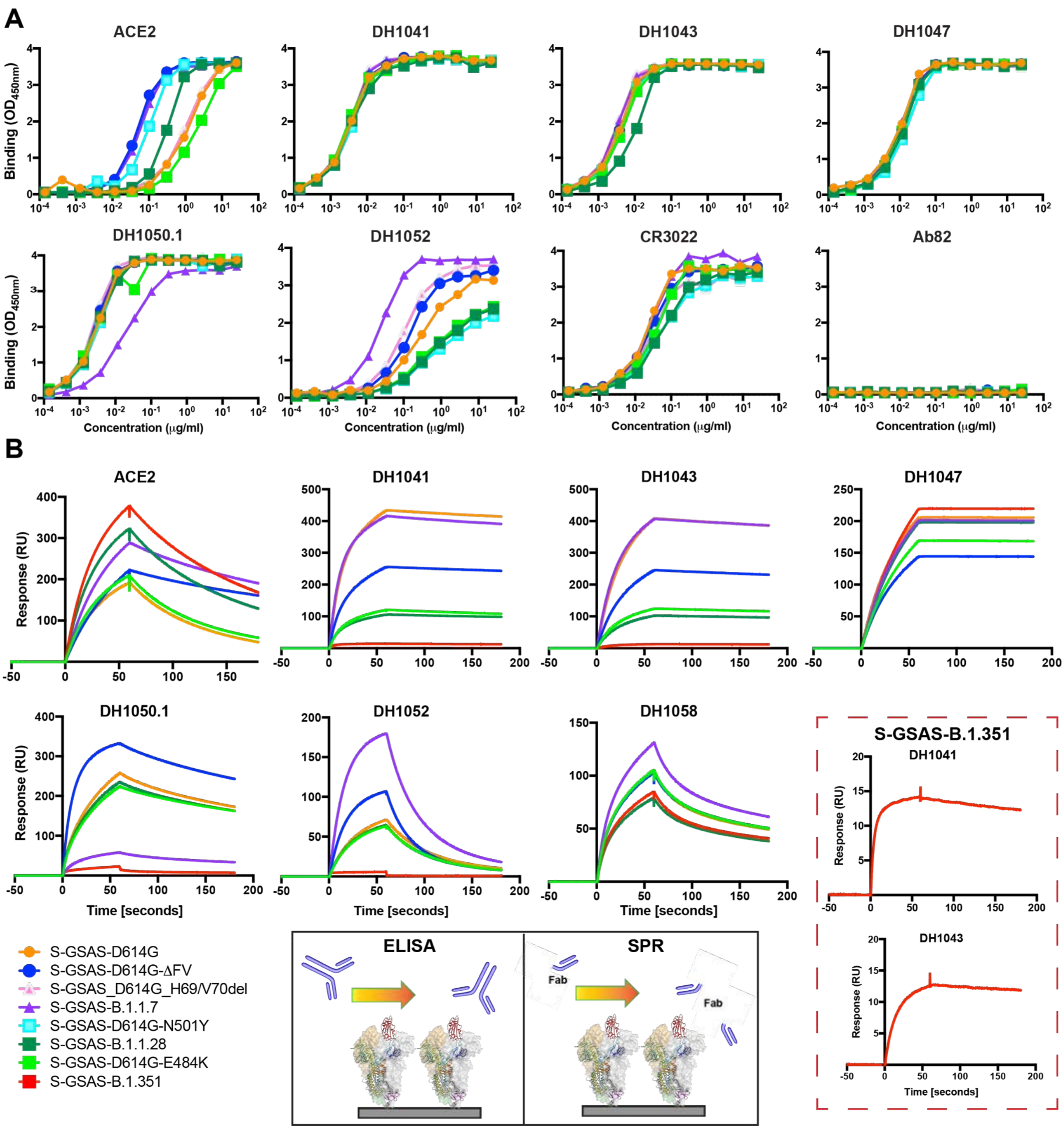


**Supplemental Item 2, Related to figures 2, 3 and 5. Binding of ACE2 receptor ectodomain, DH1041, DH1043 and DH1047 (RBD-directed), DH1050.1 and DH1052 (NTD-directed), and CR3022 (RBD-directed neutralizing antibody) to spike variants measured by (A) ELISA and (B) SPR.** The schematic shows the assay format . For the ELISA, serially diluted spike protein was bound in individual wells of 384-well plates, which were previously coated with streptavidin. Proteins were incubated and washed, then antibodies at 10 µg/ml or ACE2 with a mouse Fc tag at 2 µg/ml were added. Antibodies were incubated, washed and binding detected with goat anti-human-HRP. For the SPR, spike variants were captured on a Series S Strepavidin (SA) chip coated at 200 nM. Fabs were then injected at 200nM with a contact time of 60s and a dissociation time of 120s (50$\mu$L/min). The insets show zoomed-in views of sensorgrams to show low RU binding.

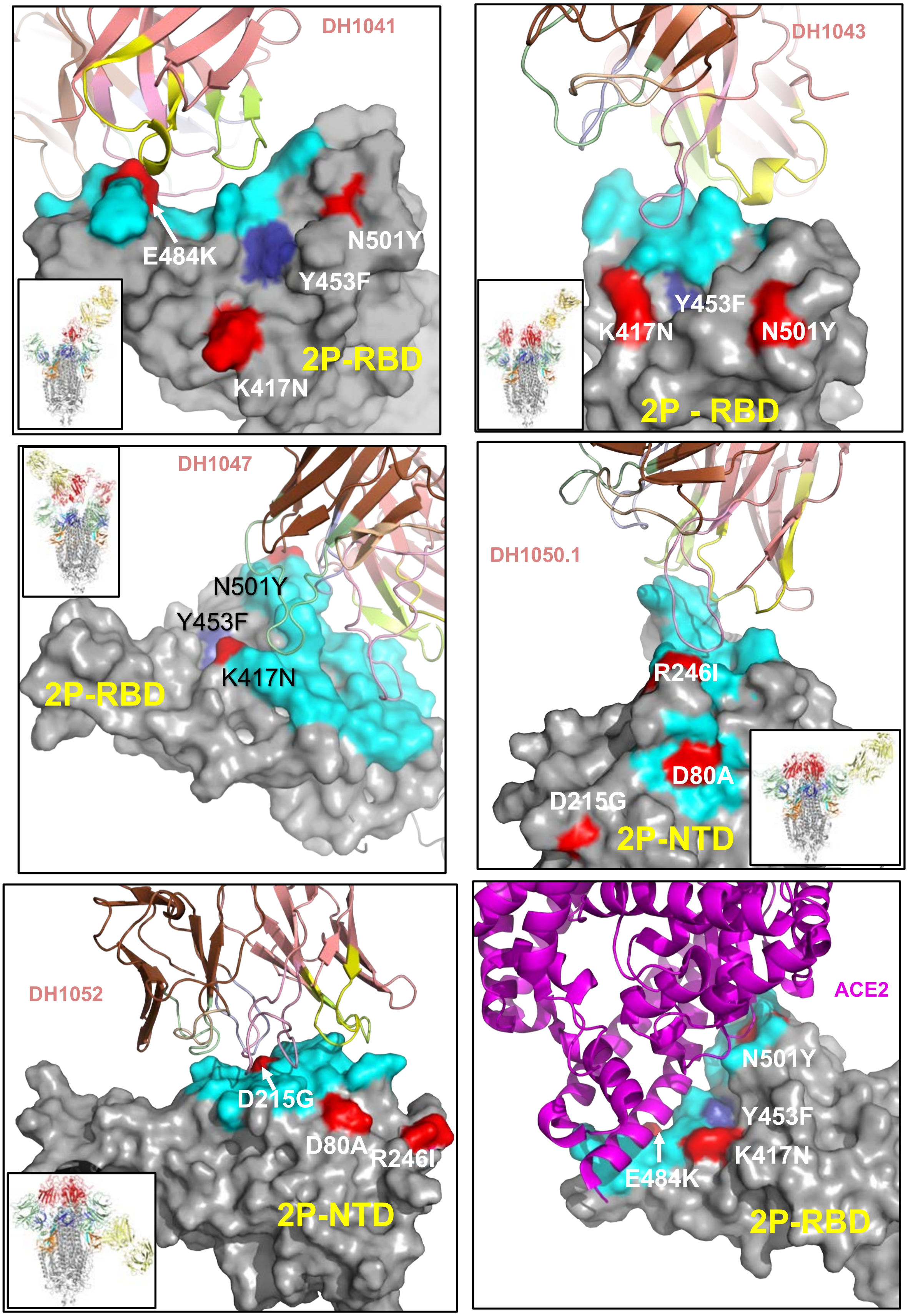
DH1041
E484K
N501Y
Y453F
2P-RBD
K417N
DH1043
Y453F
K417N
N501Y
2P - RBD
DH1047
N501Y
Y453F
K417N
2P-RBD
DH1050.1
R246I
D80A
D215G
2P-NTD
DH1052
D215G
D80A
R246I
2P-NTD
ACE2
N501Y
Y453F
K417N
E484K
2P-RBD

**Supplemental Item 3, Related to Figures 2, 3 and 5. Binding footprints of ACE2 and antibodies on SARS-CoV-2 S RBD and NTD**. RBD and NTD are shown as grey surface. Bound antibodies and ACE2 are shown in cartoon representation. ACE2 is colored purple and antibodies heavy chain are colored salmon and light chain brown. The HCDR1 are yellow, HCDR2 limon green, HCDR3 pink, LCDR1 pale green, LCDR2 wheat and LCDR3 light blue. Mutations from the B.1.351 spike variant are colored red, and Y453F from the mink cluster 5 associated mutation is colored blue. PDB entries 7LAA (DH1041), 7LD1 (DH1047), 6M0J (ACE2), 7LCN (DH1050.1), 7LAB (DH1052) and 7LJR (DH1043) for the complexes with the S-GSAS/PP spike protein were used.

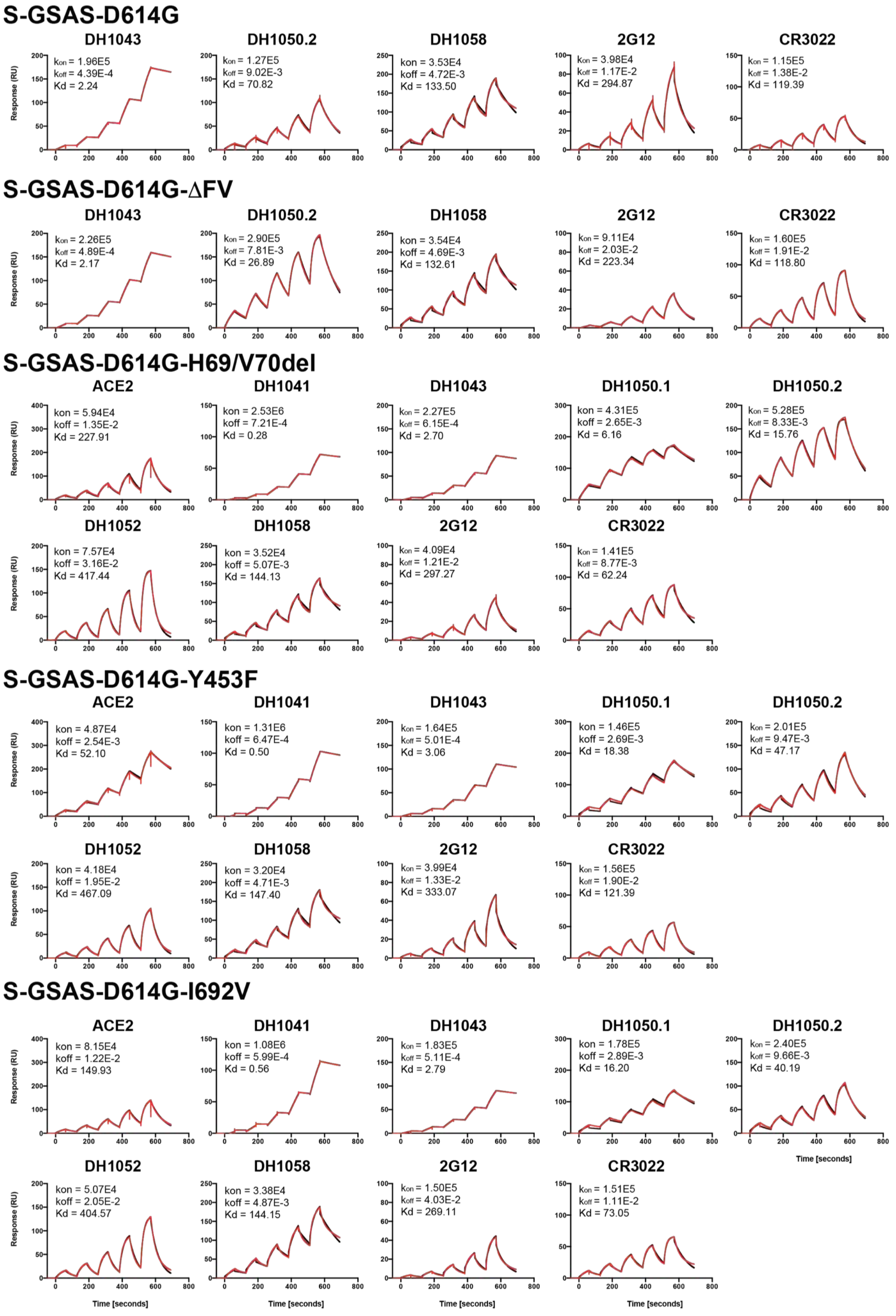


**Supplemental Item 4. Kinetics of RBD, NTD and S2 directed antibody Fabs binding to spike variants measured by SPR, Related to Figures 2, 3 and 5.** The kinetic data is in red and the fitted data is in black.

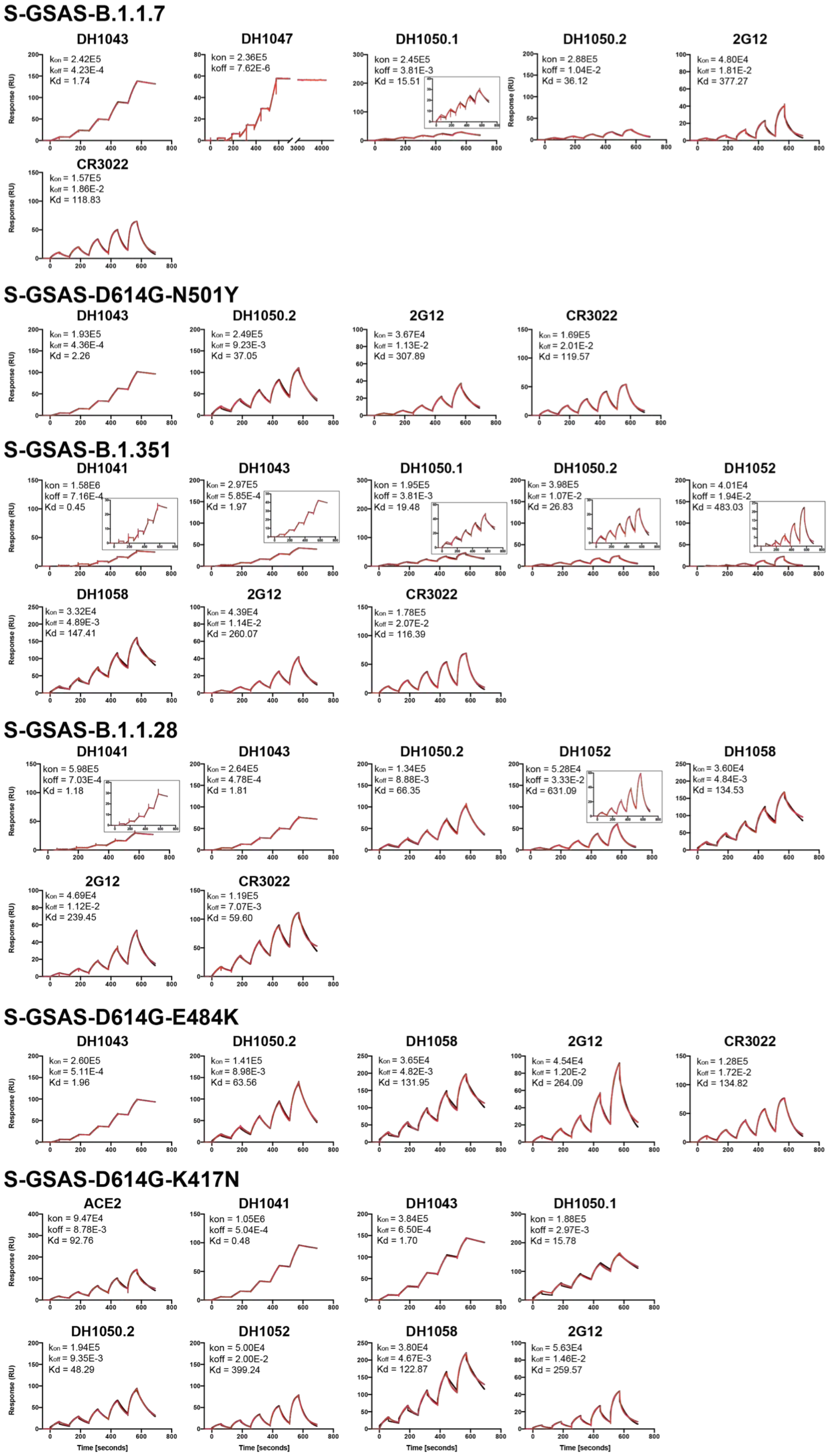


**Supplemental Item 4, continued. Kinetics of RBD, NTD and S2 directed antibody Fabs binding to spike variants measured by SPR, Related to Figures 2, 3 and 5.** The kinetic data is in red and the fitted data is in black.

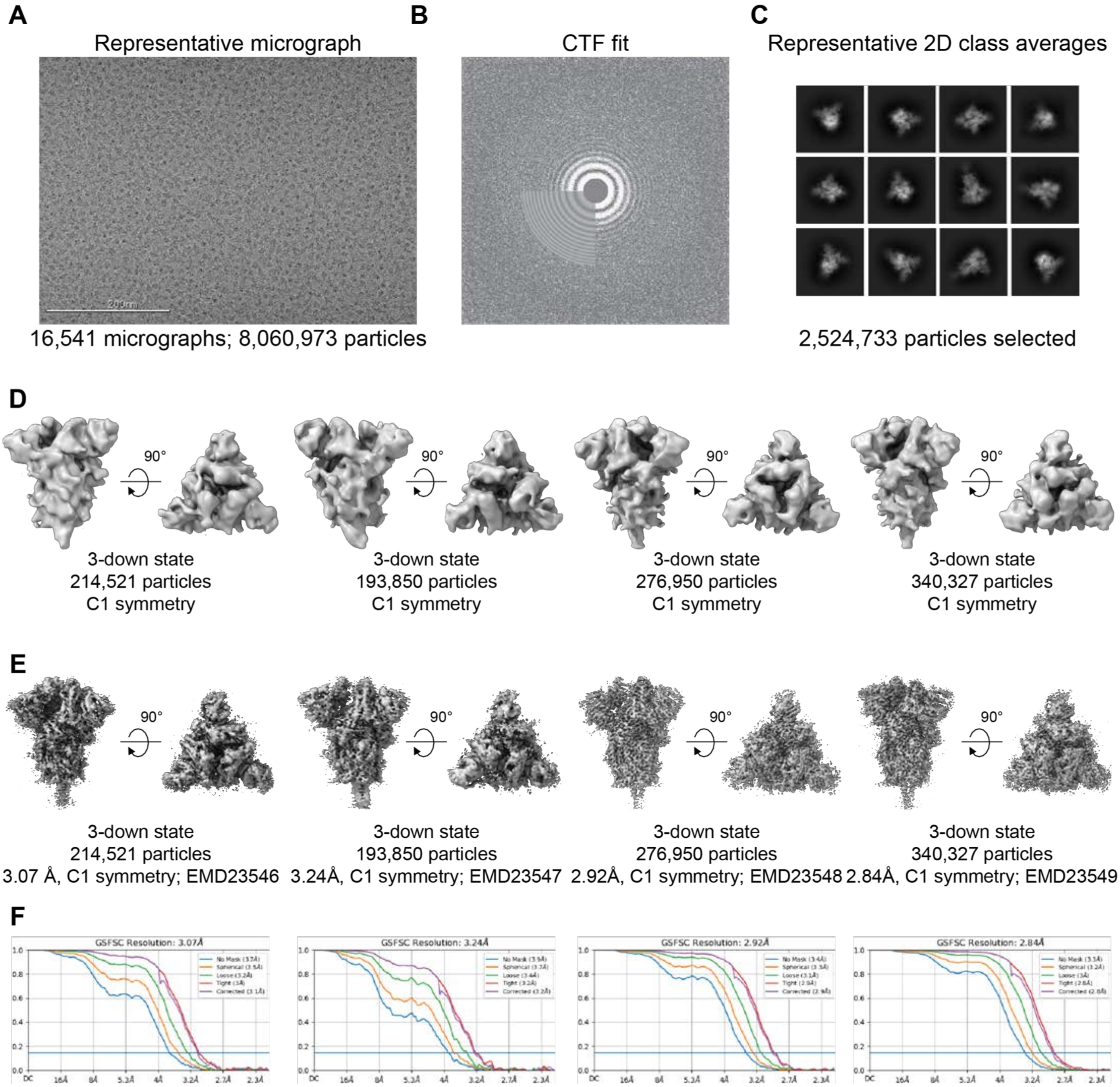


**Supplemental Item 5. Cryo-EM data processing for the mink-associated S-GSAS-D614G-ΔFV SARS-CoV-2 S ectodomain, Related to Figure 2. A**. Representative cryo-EM micrograph. **B**. Cryo-EM CTF fit. **C**. Representative 2D class averages from Cryo-EM dataset. **D.** *Ab initio* reconstructions for the cryo-EM 3-down states. **E**. Refined maps for the cryo-EM 3-down states. **F.** Fourier shell correlation curves for the cryo-EM 3-down states. **G.** *Ab initio* reconstructions for the cryo-EM 1-up, 2-up and new states. **H**. Refined maps for the cryo-EM 1-up, 2-up and new states. **I.** Fourier shell correlation curves for the cryo-EM 1-up, 2-up and new states.

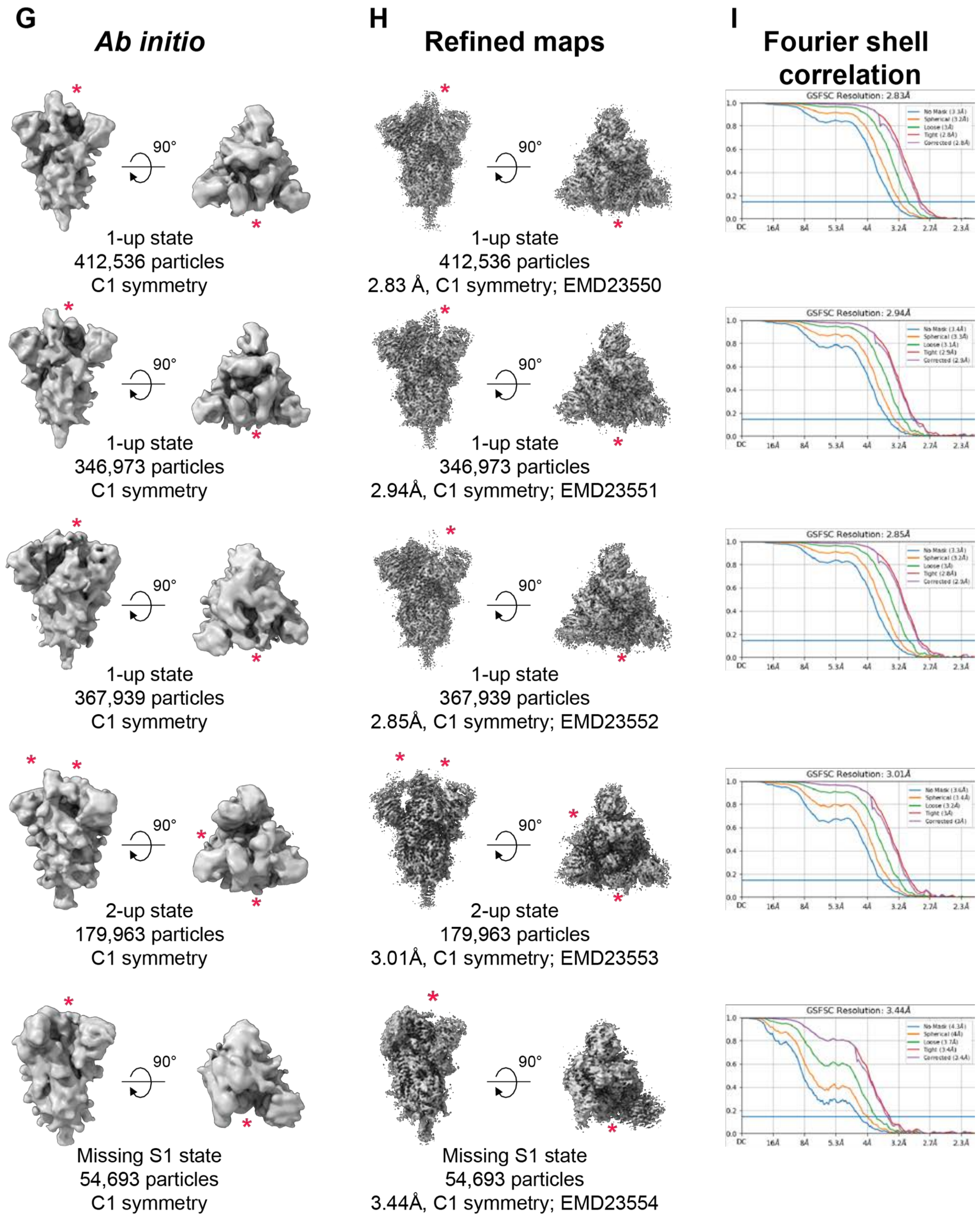


**Supplemental Item 5 continued. Cryo-EM data processing for the mink-associated S-GSAS-D614G-ΔFV SARS-CoV-2 S ectodomain, Related to Figure 2. A**. Representative cryo-EM micrograph. **B**. Cryo-EM CTF fit. **C**. Representative 2D class averages from Cryo-EM dataset. **D.** *Ab initio* reconstructions for the cryo-EM 3-down states. **E**. Refined maps for the cryo-EM 3-down states. **F.** Fourier shell correlation curves for the cryo-EM 3-down states. **G.** *Ab initio* reconstructions for the cryo-EM 1-up, 2-up and new states. **H**. Refined maps for the cryo-EM 1-up, 2-up and new states. **I.** Fourier shell correlation curves for the cryo-EM 1-up, 2-up and new states.

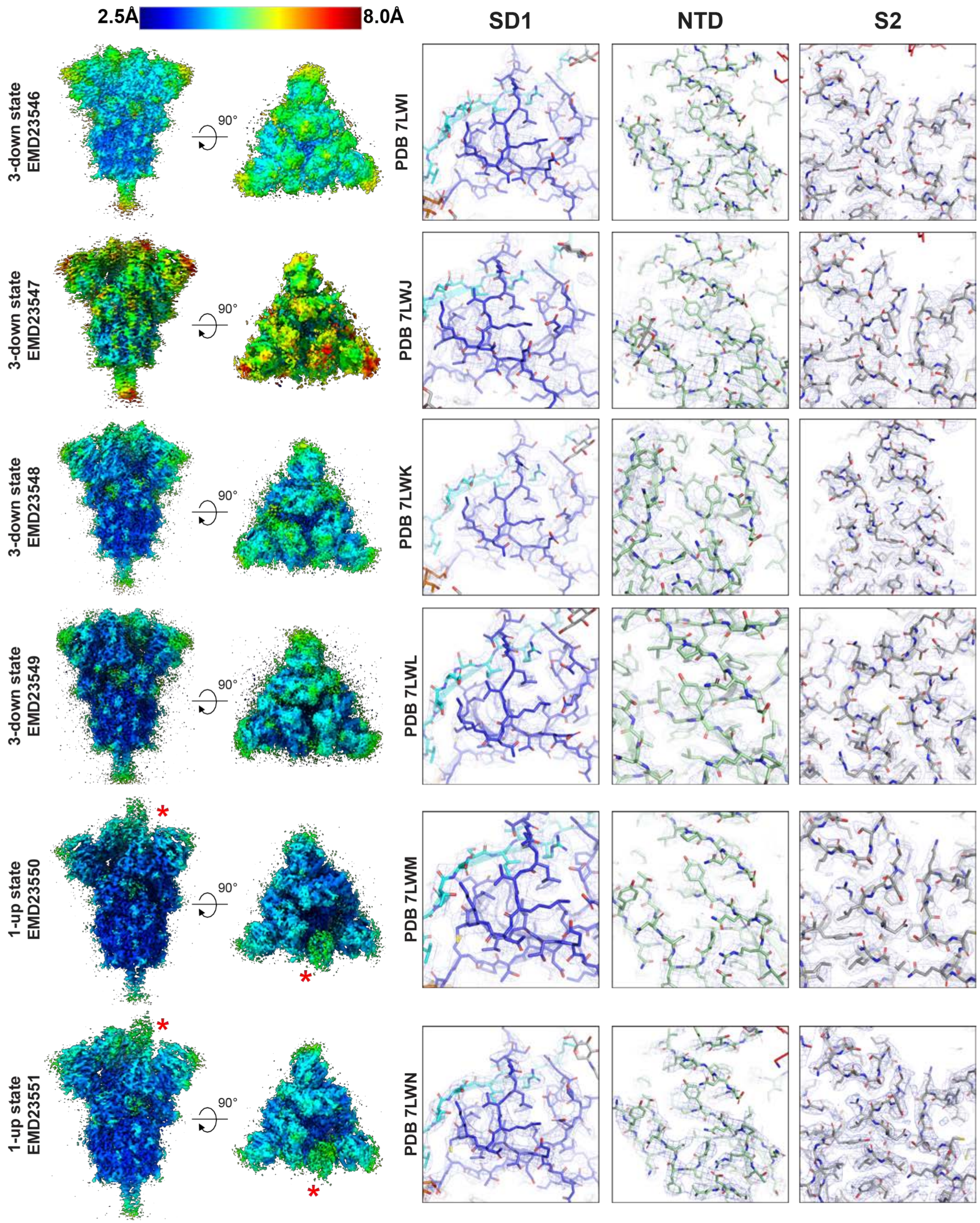


**Supplemental Item 6. Details of the mink-associated S-GSAS-D614G-ΔFV SARS-CoV-2 S ectodomain cryo-EM map and model fitting, Related to Figure 2. Left.** Refined cryo-EM maps colored by local resolution **Right.** Zoom-in images showing the SD1, NTD and S2 regions in the structures. The cryo-EM map is shown as a mesh surface and the fitted model is in cartoon representation, with residues shown as sticks.

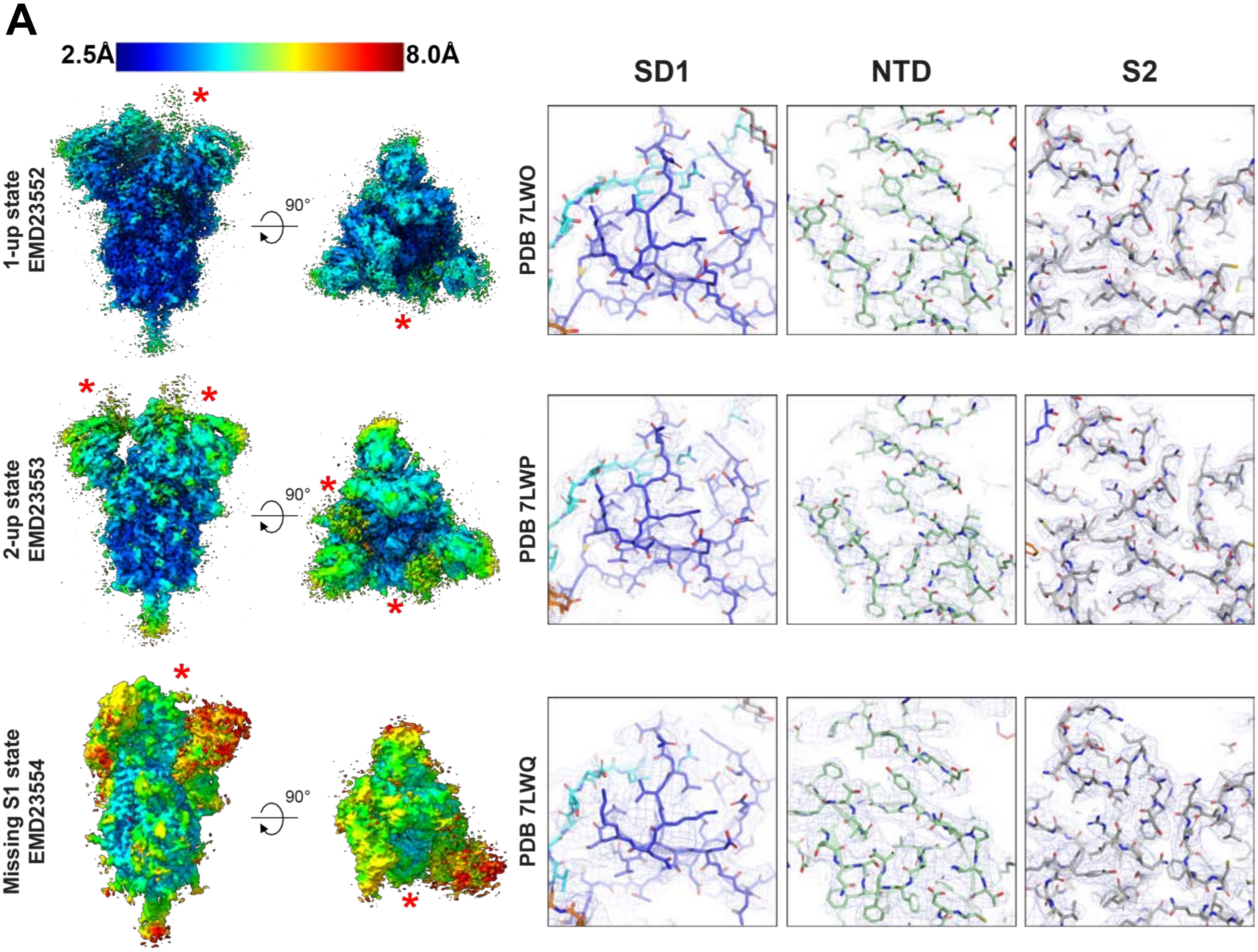


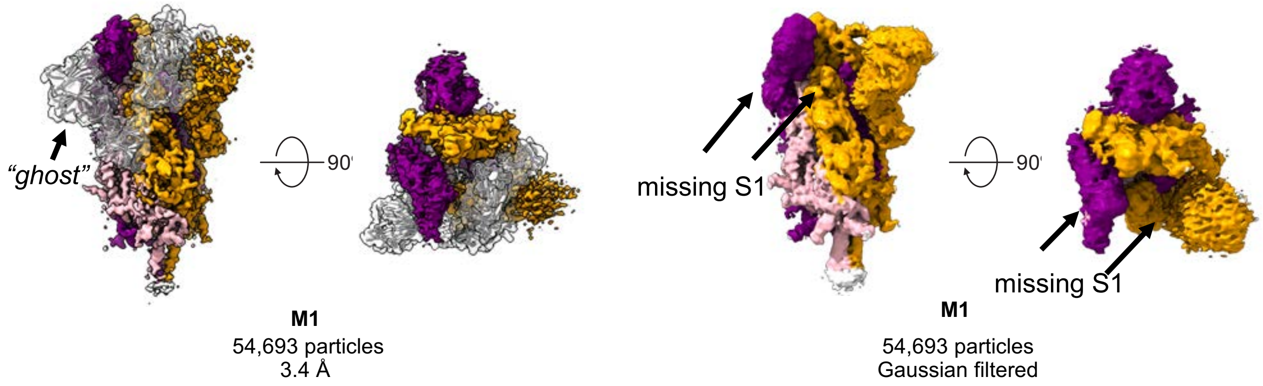


**Supplemental Item 6 continued. Details of the mink-associated S-GSAS-D614G-ΔFV SARS-CoV-2 S ectodomain cryo-EM map and model fitting, Related to Figure 2. Top. Left.** Refined cryo-EM maps colored by local resolution Top. **Right.** Zoom-in images showing the SD1, NTD and S2 regions in the structures. The cryo-EM map is shown as a mesh surface and the fitted model is in cartoon representation, with residues shown as sticks. **Bottom. Left.** Refined cryo-EM reconstruction of M1, colored by protomer in orange, purple and pink. The missing S1 domain of the protomer in pink is depicted by grey/white structure ("ghost") to show the missing regions. **Bottom. Right.** Gaussian filtered M1 map with arrows showing the missing regions.

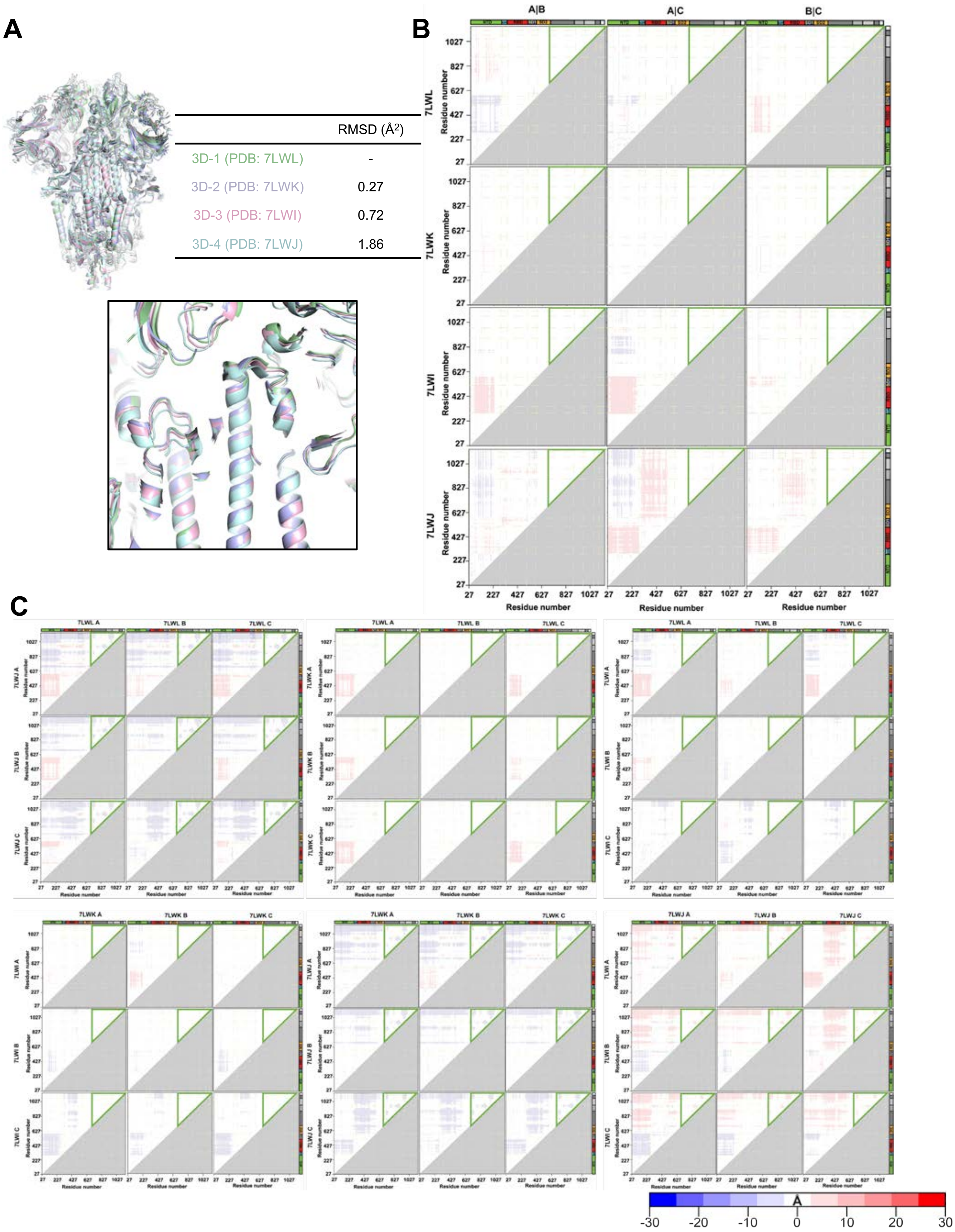
A
RMSD (Å2)
3D-1 (PDB: 7LWL) -
3D-2 (PDB: 7LWK) 0.27
3D-3 (PDB: 7LWI) 0.72
3D-4 (PDB: 7LWJ) 1.86
B
A|B
A|C
B|C
7LWL
7LWK
7LWI
7LWJ
Residue number
27 227 427 627 827 1027
C
7LWL A
7LWL B
7LWL C
7LWJ A
7LWJ B
7LWJ C
7LWK A
7LWK B
7LWK C
7LWI A
7LWI B
7LWI C
-30 -20 -10 0 10 20 30
Å

**Supplemental Item 7. Variability between in the mink-associated S-GSAS-D614G-ΔFV SARS-CoV-2 S ectodomain 3-RBD-down structures, Related to Figure 2.** **A.** Superposition of structures aligned on residues 908-1035 spanning the HR1-CH regions. Table lists RMSD of each structure superposed on 3D-1 (PDB: 7LWL). Inset shows zoomed-in view of the 908-1035 region. **B.** DDM of the 3-RBD down structures comparing each chain of a structure to the other chains of the same structure. **C.** DDM of the 3-RBD-down structures comparing each chain to each chains of the other 3-down structures.

A

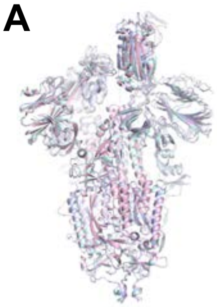

| | RMSD (Å$^2$) |
|---|---|
| 1U-1 (PDB: 7LWM) | - |
| 1U-2 (PDB: 7LWN) | 0.33 |
| 1U-3 (PDB: 7LWO) | 0.18 |

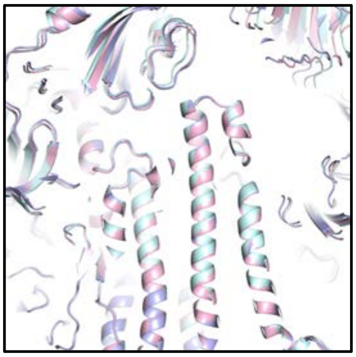

B

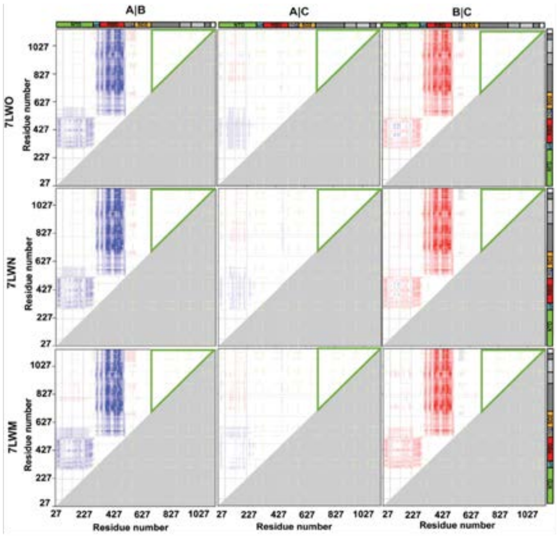


C

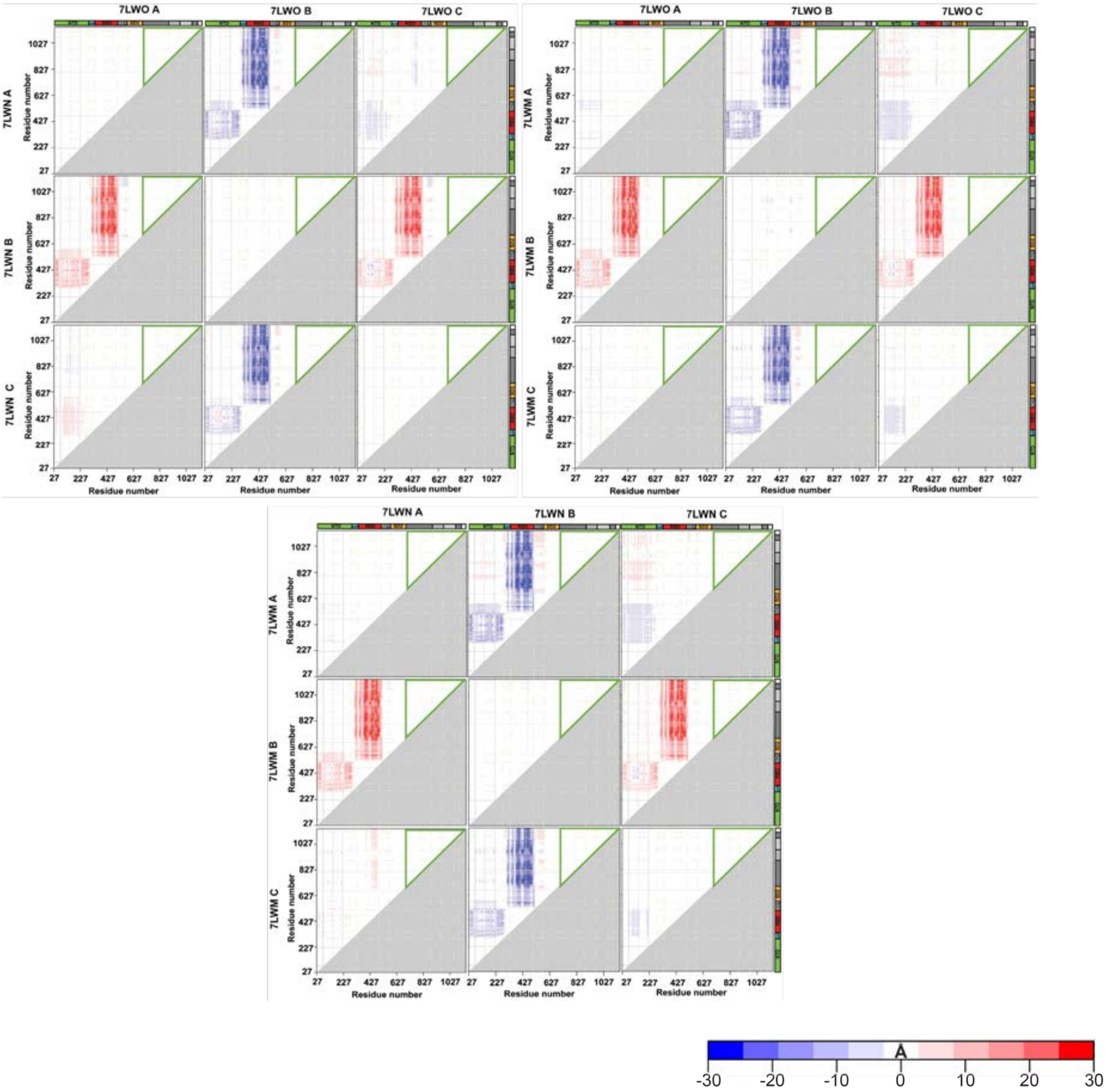

**Supplemental Item 8. Variability between in the mink-associated S-GSAS-D614G-ΔFV SARS-CoV-2 S ectodomain 3-RBD-down structures, Related to Figure 2. A.** Superposition of structures aligned on residues 908-1035 spanning the HR1-CH regions. Table lists RMSD of each structure superposed on 1U-1 (PDB: 7LWM). Inset shows zoomed-in view of the 908-1035 region. **B.** DDM of the 1-RBD-up structures comparing each chain of a structure to the other chains of the same structure. **C.** DDM of the 1-RBD-up structures comparing each chain to each chains of the other 1-RBD-up structures.

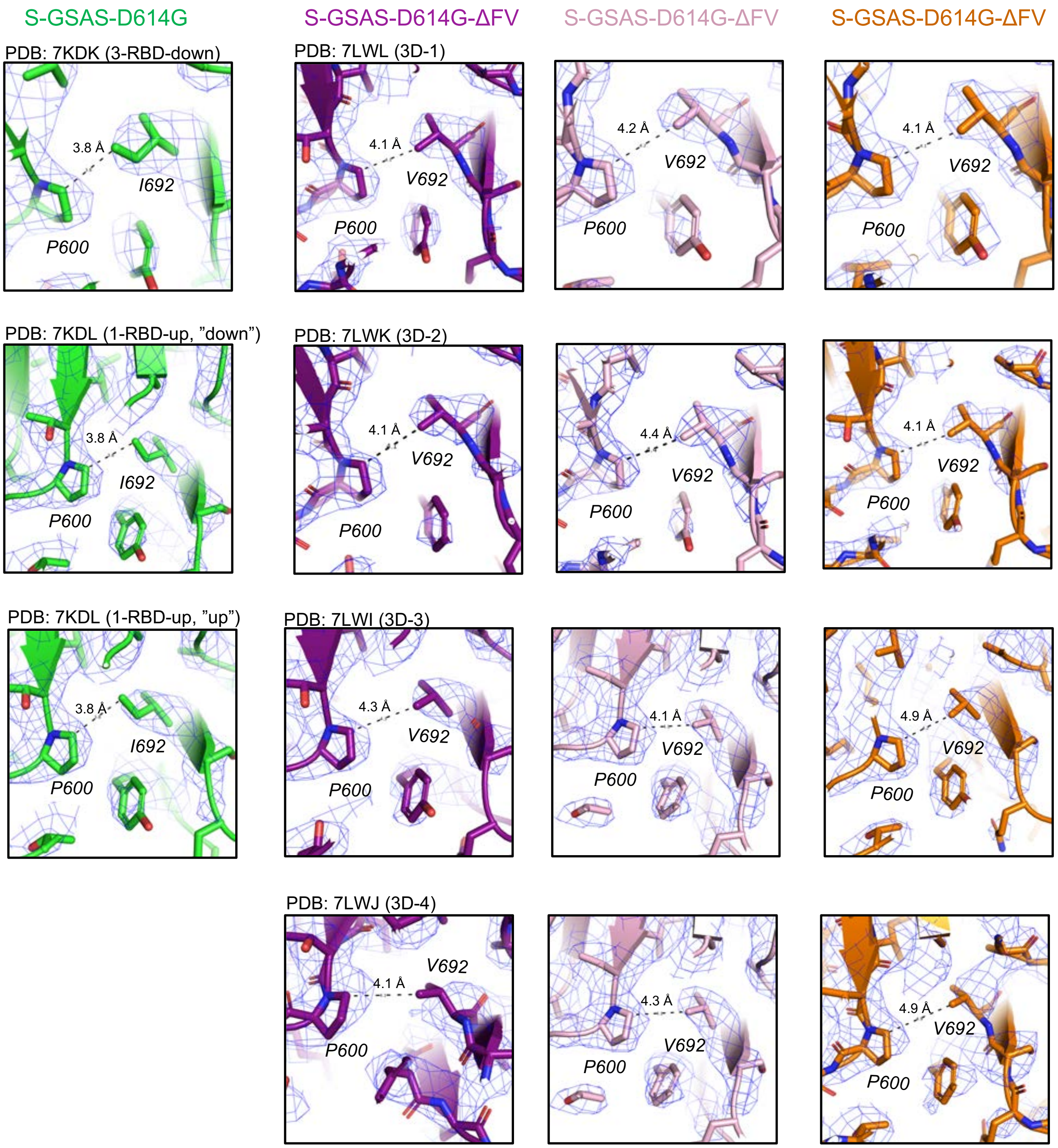


**Supplemental Item 9. The I692V mutation in the mink-associated S-GSAS-D614G-ΔFV SARS-CoV-2 S ectodomain, Related to Figure 2. Left.** Zoomed-in views of the region around residue I692 in S-GSAS-D614G. **Right.** Zoomed-in views of the region around the I692V mutation the in the S-GSAS-D614G-ΔFV SARS-CoV-2 S ectodomain structures. The cryo-EM map is shown as a mesh and the fitted model is in cartoon representation, with residues shown as sticks.

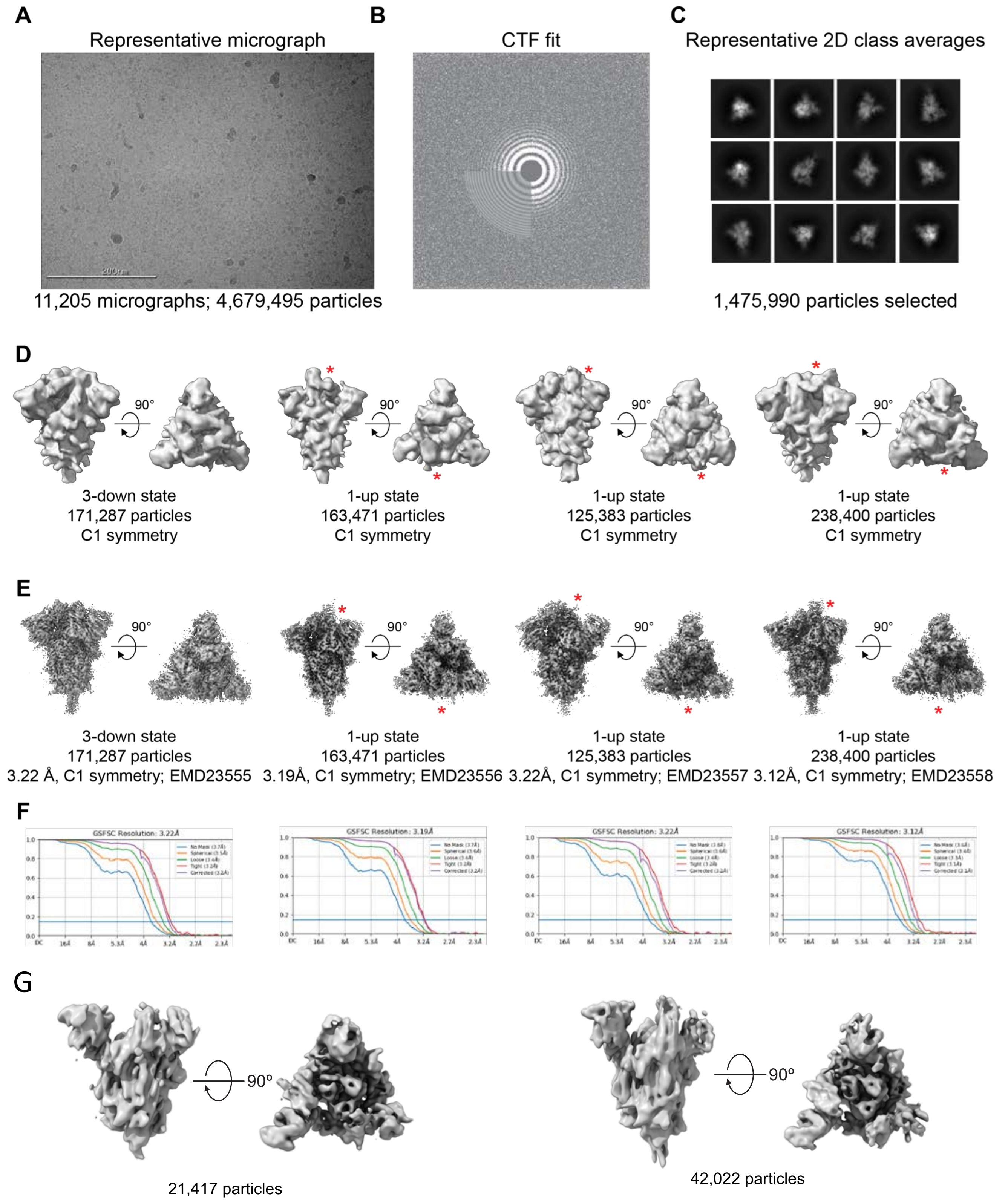


**Supplemental Item 10. Cryo-EM data processing for the S-GSAS-B.1.1.7 SARS-CoV-2 S ectodomain variant, Related to Figure 3. A**. Representative cryo-EM micrograph. **B**. Cryo-EM CTF fit. **C**. Representative 2D class averages from Cryo-EM dataset. **D.** *Ab initio* reconstructions for the cryo-EM 3-down and 1-up states. **E**. Refined maps for the cryo-EM 3-down and 1-up states. **F.** Fourier shell correlation curves for the cryo-EM 3-down and 1-up states. **G.** Additional populations of RBD-up states..

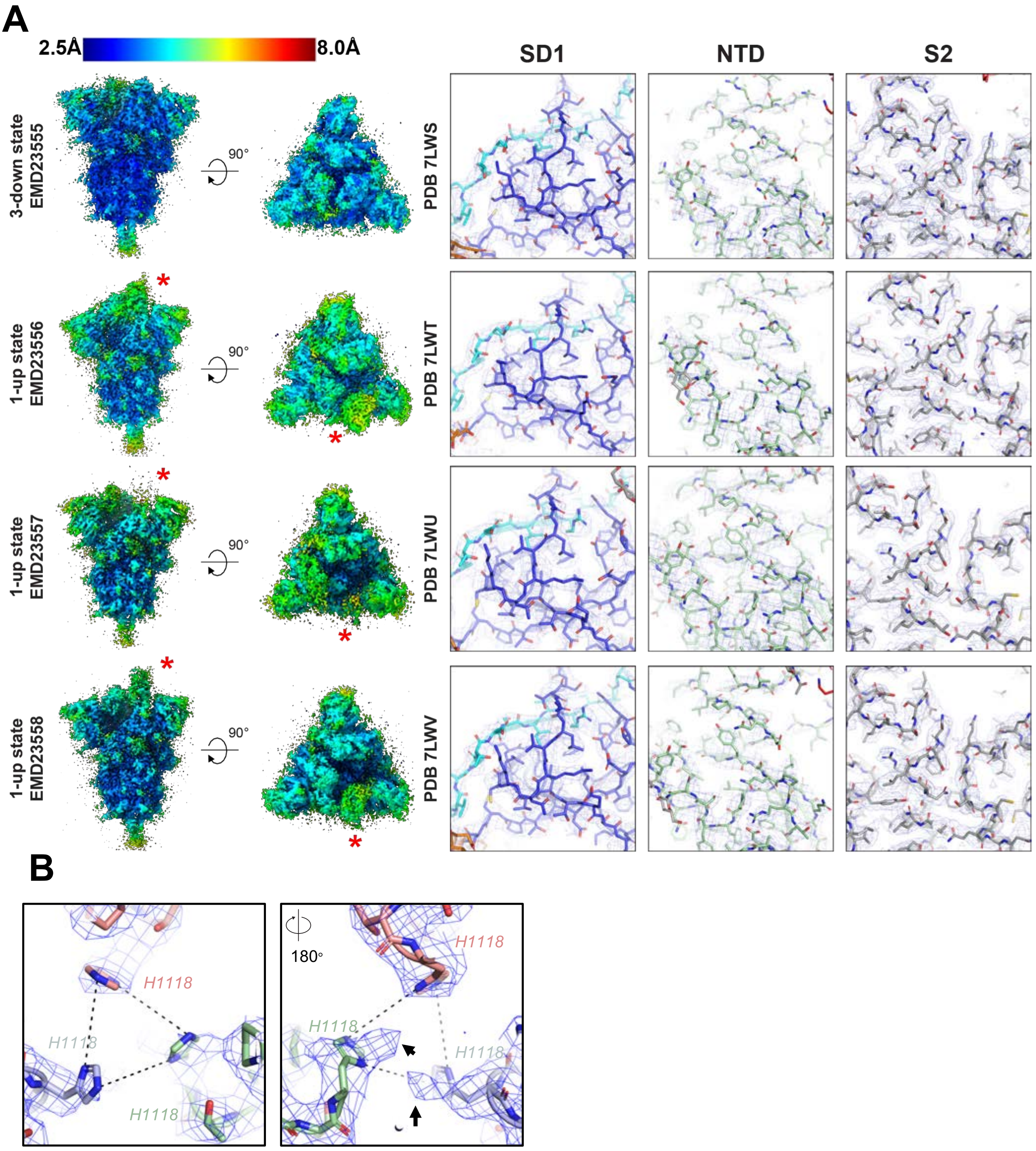


**Supplemental Item 11. Details of the S-GSAS-B.1.1.7 SARS-CoV-2 S ectodomain cryo-EM map and model fitting, Related to Figure 3. A. Left.** Refined cryo-EM maps colored by local resolution **Right.** Zoom-in images showing the SD1, NTD and SD2 regions in the structures. The cryo-EM map is shown as a mesh surface and the fitted model is in cartoon representation, with residues shown as sticks. **B.** Zoomed-in view of the H1118 histidine triad, shown in two different orientations. The arrows point to the additional visible density for alternate conformations of the H1118 residue.

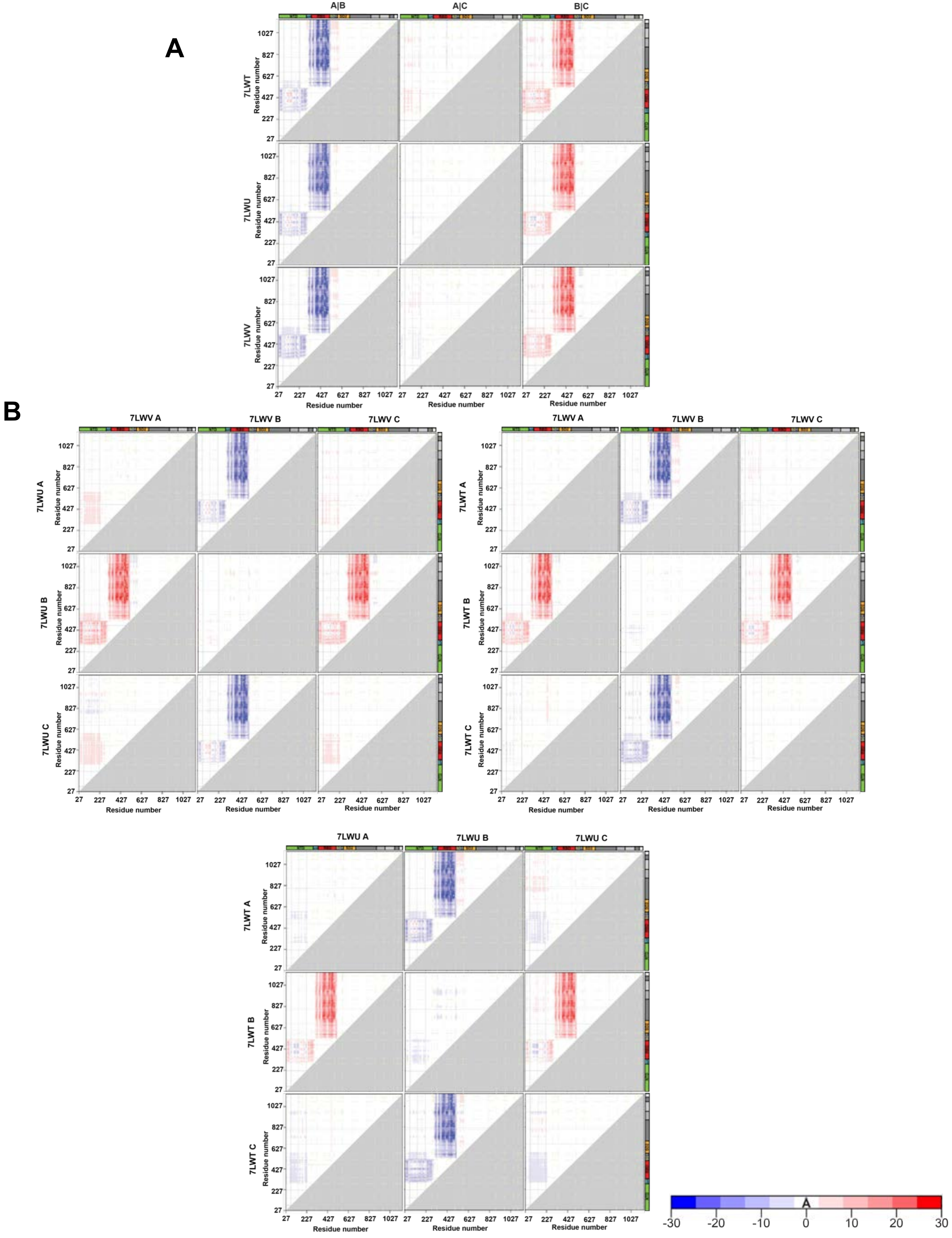


**Supplemental Item 12. Difference distance matrices (DDM) showing structural changes between different protomers for the S-GSAS-B.1.1.7 SARS-CoV-2 S ectodomain structures. Related to Figure 3 A.** DDM of the 1-RBD up structures comparing each chain of a structure to the other chains of the same structure. **B.** DDM of the 1-RBD up structures comparing each chain to each chains of the other 1-up structures.

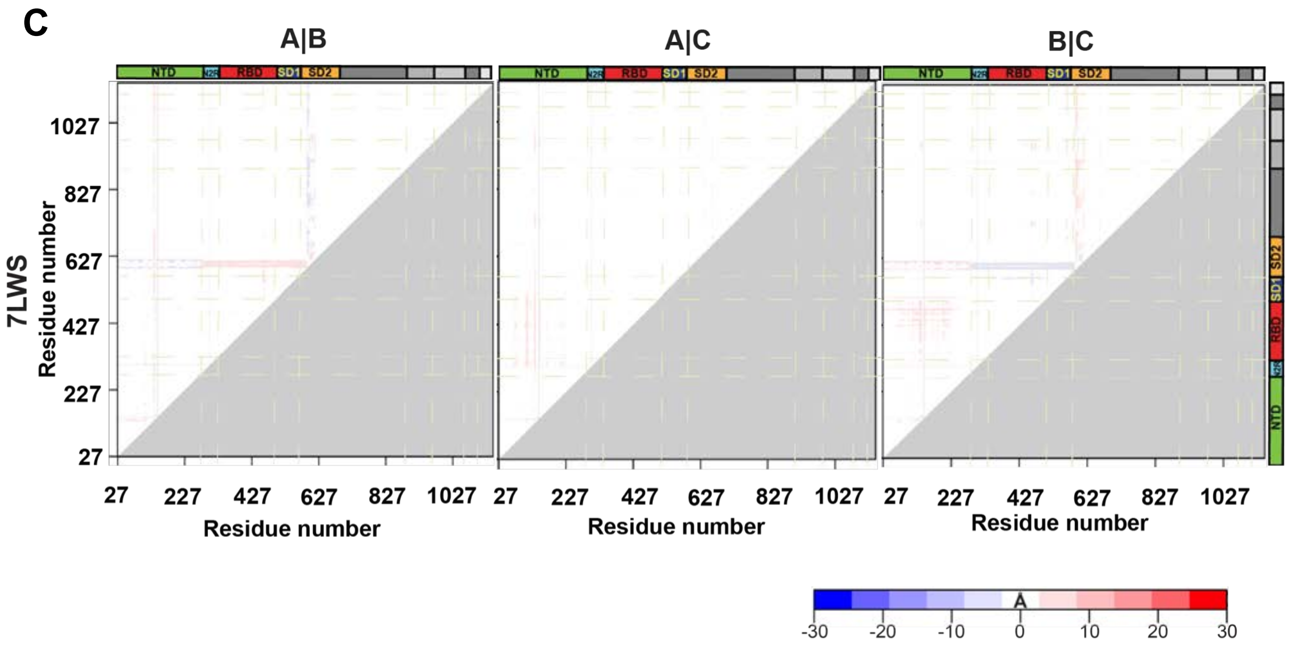


**Supplemental Item 12 continued. Difference distance matrices (DDM) showing structural changes between different protomers for the S-GSAS-B.1.1.7 SARS-CoV-2 S ectodomain structures. Related to Figure . C.** DDM of the 3-RBD down structures comparing each chain of a structure to the other chains of the same structure..

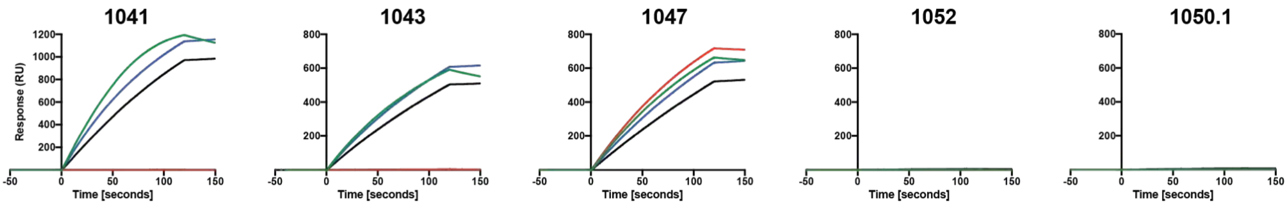


**Supplemental Item 13. Binding of RBD, NTD and S2 directed antibody to the RBD of spike variants measured by SPR, Related to Figure 5.**

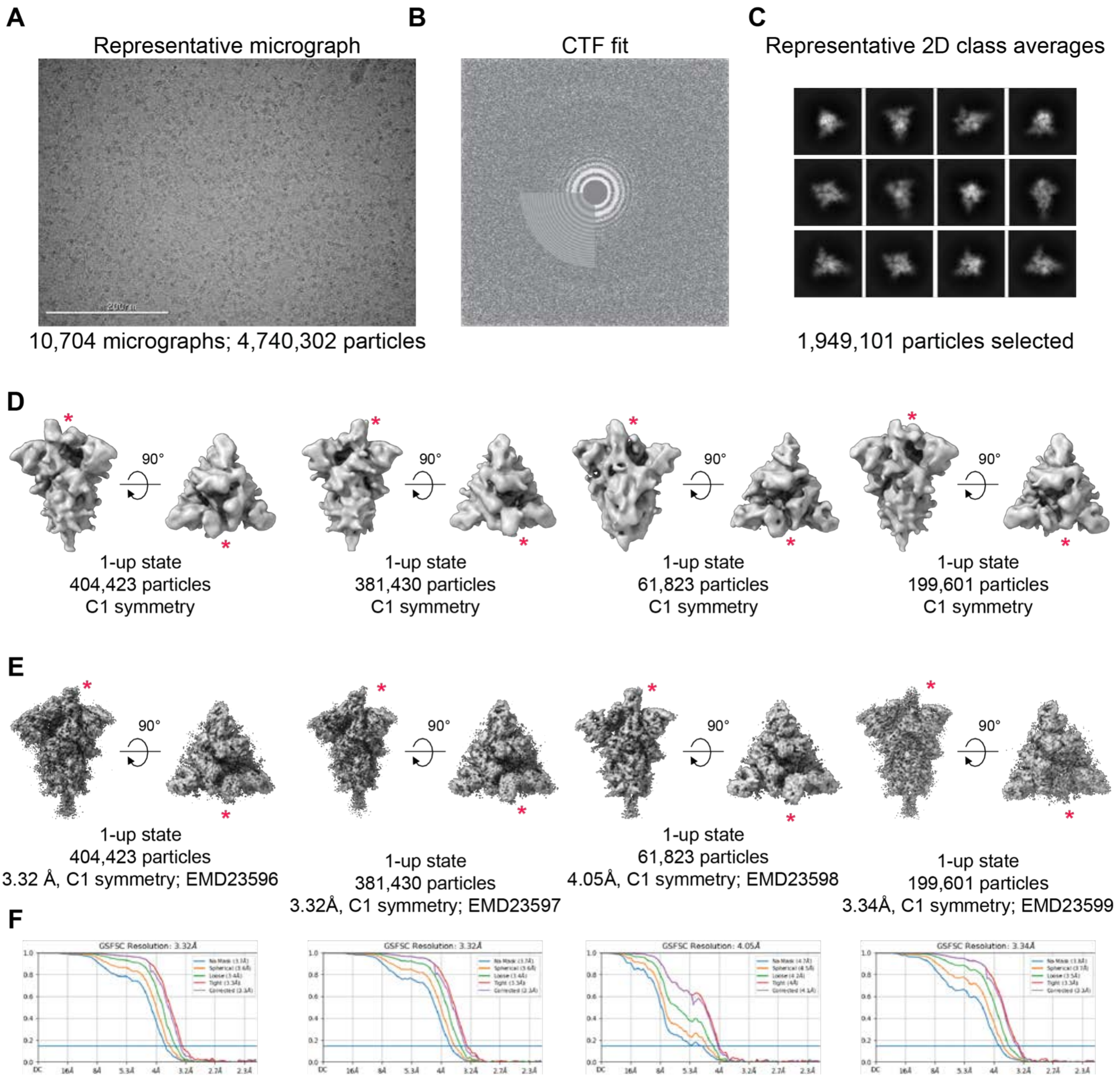


**Supplemental Item 14. Cryo-EM data processing for the S-GSAS-B.1.351 SARS-CoV-2 S ectodomain variant, Related to Figure 6. A**. Representative cryo-EM micrograph. **B**. Cryo-EM CTF fit. **C**. Representative 2D class averages from Cryo-EM dataset. **D.** *Ab initio* reconstructions for the cryo-EM 1-up states. **E.** Refined maps for the cryo-EM 1-up states. **F.** Fourier shell correlation curves for the cryo-EM1-up states. **G.** *Ab initio* reconstructions for the cryo-EM 2-up and 3-down states. **E.** Refined maps for the cryo-EM 2-up and 3-down states. **F.** Fourier shell correlation curves for the cryo-EM 2-up and 3-down states.

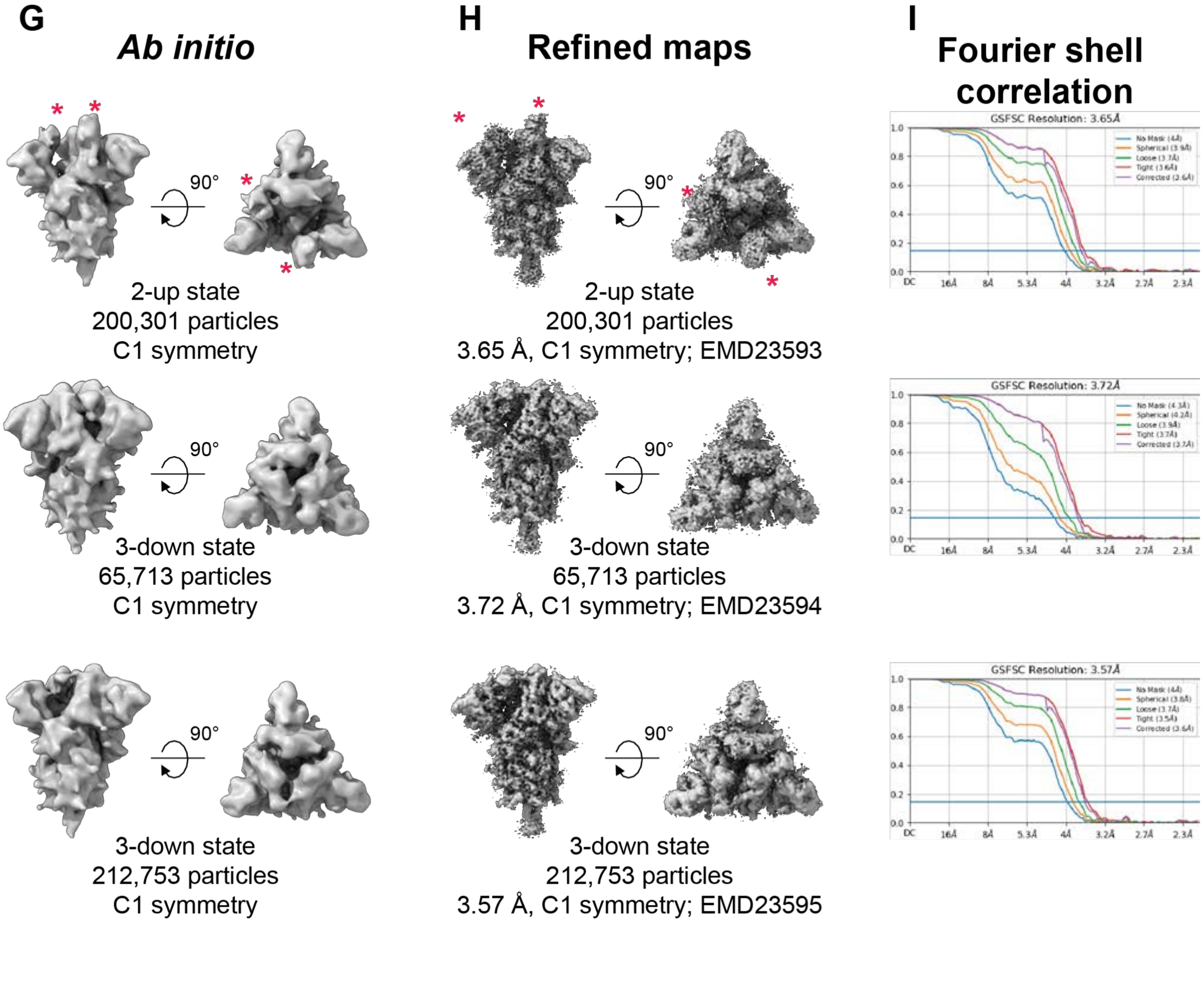


**Supplemental Item 14 continued. Cryo-EM data processing for the S-GSAS-B.1.351 SARS-CoV-2 S ectodomain variant, Related to Figure 6. A**. Representative cryo-EM micrograph. **B**. Cryo-EM CTF fit. **C**. Representative 2D class averages from Cryo-EM dataset. **D.** *Ab initio* reconstructions for the cryo-EM 1-up states. **E.** Refined maps for the cryo-EM 1-up states. **F.** Fourier shell correlation curves for the cryo-EM1-up states. **G.** *Ab initio* reconstructions for the cryo-EM 2-up and 3-down states. **E.** Refined maps for the cryo-EM 2-up and 3-down states. **F.** Fourier shell correlation curves for the cryo-EM 2-up and 3-down states.

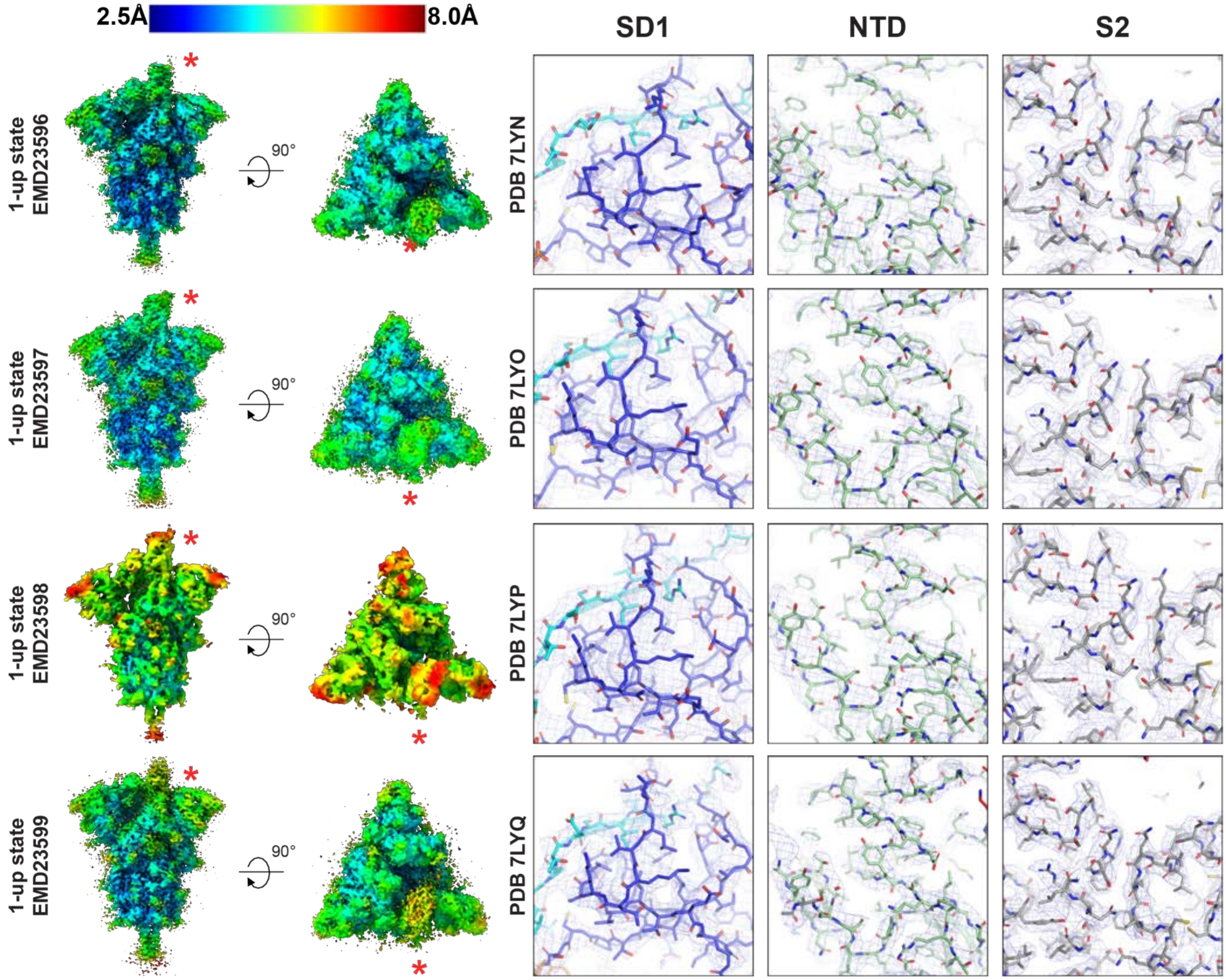


**Supplemental Item 15. Quality assessment of the S-GSAS-B.1.351 SARS-CoV-2 S ectodomain cryo-EM map and model fitting, Related to Figure 6. Left.** Refined cryo-EM maps colored by local resolution **Right.** Zoom-in images showing the SD1, NTD and S2 regions in the structures. The cryo-EM map is shown as a mesh surface and the fitted model is in cartoon representation, with residues shown as sticks.

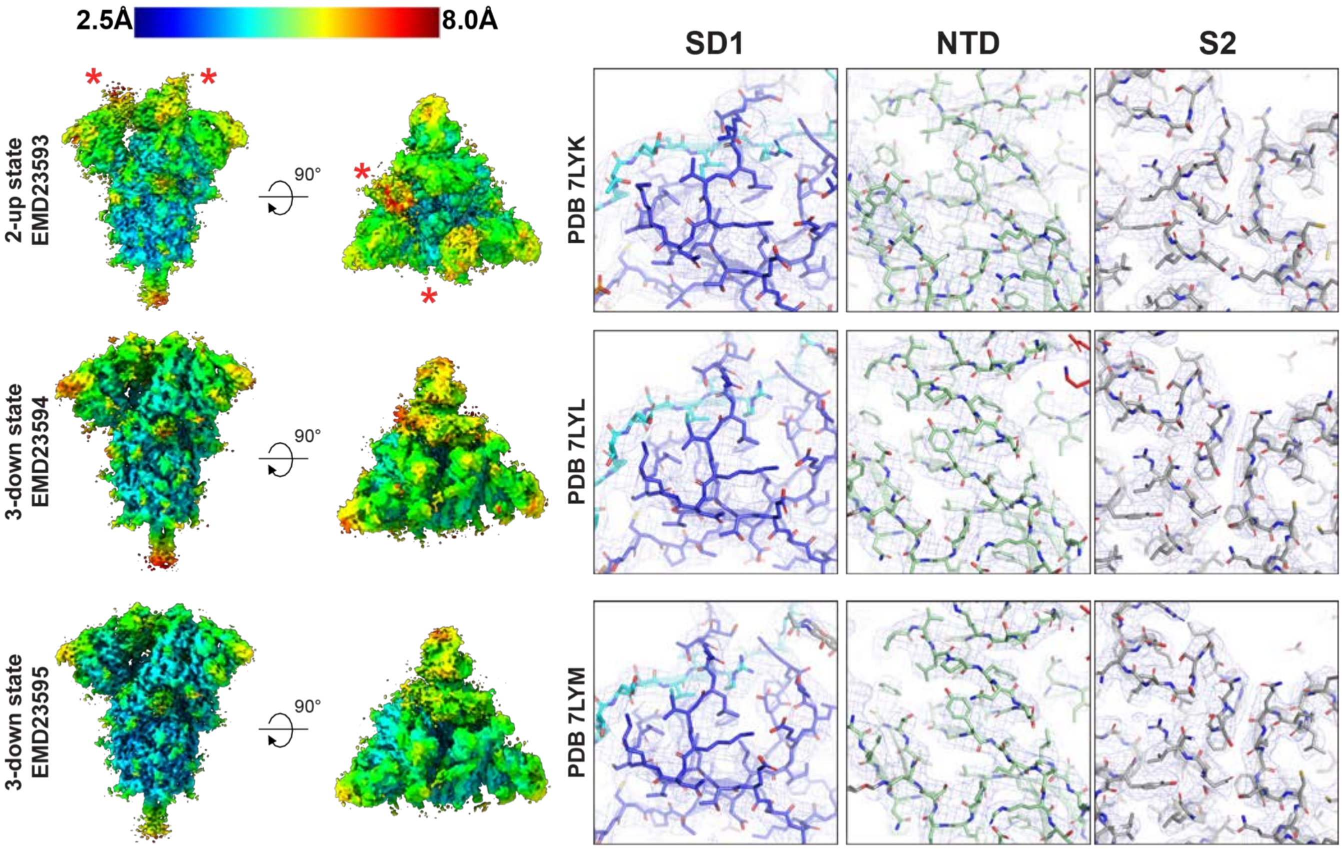


**Supplemental Item 15, continued. Quality assessment of the S-GSAS-B.1.351 SARS-CoV-2 S ectodomain cryo-EM map and model fitting, Related to Figure 6. Left.** Refined cryo-EM maps colored by local resolution **Right.** Zoom-in images showing the SD1, NTD and S2 regions in the structures. The cryo-EM map is shown as a mesh surface and the fitted model is in cartoon representation, with residues shown as sticks.

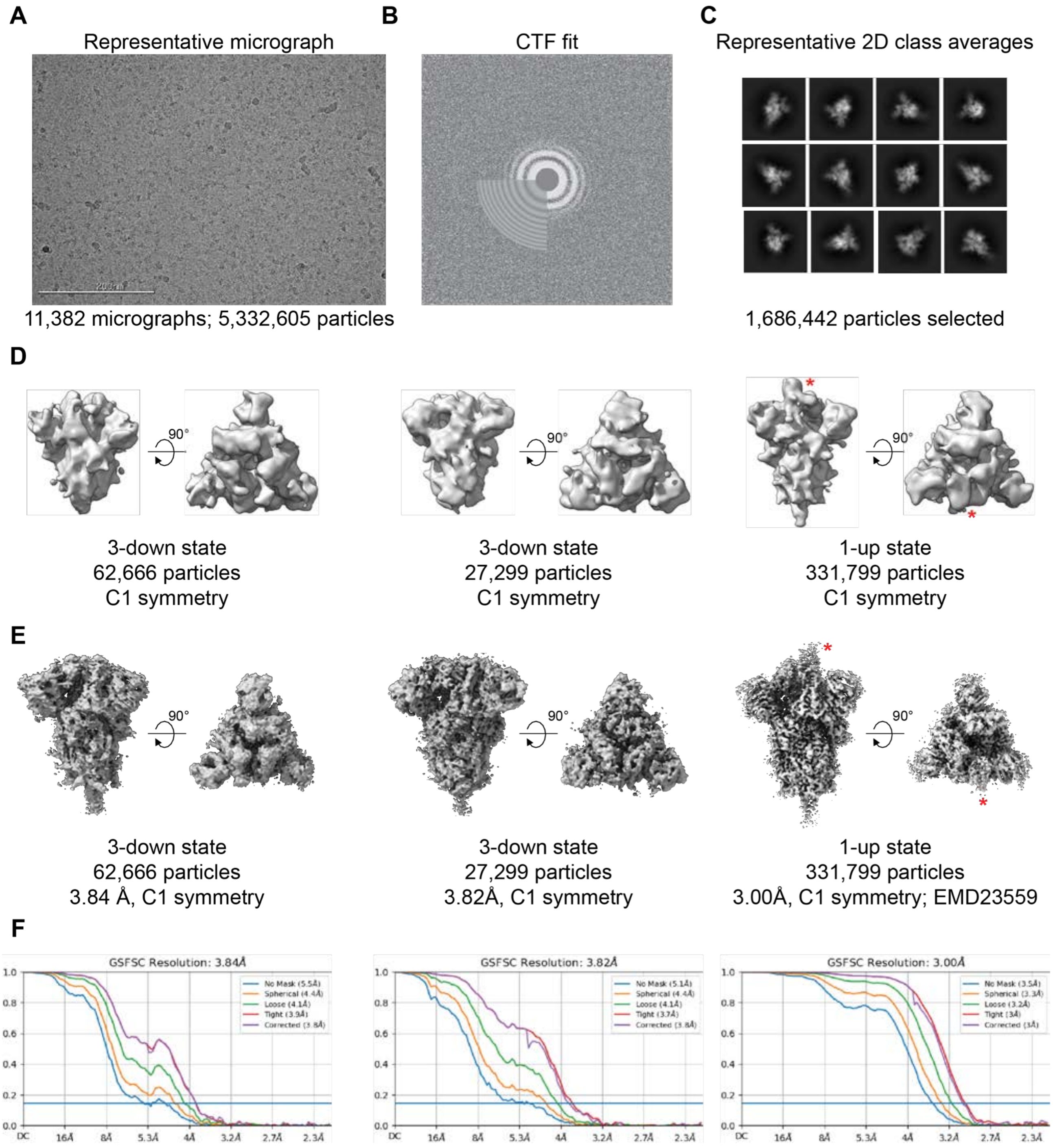


**Supplemental Item 16. Cryo-EM data processing for the S-GSAS-B.1.1.28 SARS-CoV-2 S ectodomain variant, Related to Figure 6. A**. Representative cryo-EM micrograph. **B**. Cryo-EM CTF fit. **C**. Representative 2D class averages from Cryo-EM dataset. **D.** *Ab initio* reconstructions for the cryo-EM 3-down and 1-up states. **E**. Refined maps for the cryo-EM 3-down and 1-up states. **F.** Fourier shell correlation curves for the cryo-EM 3-down and 1-up states.

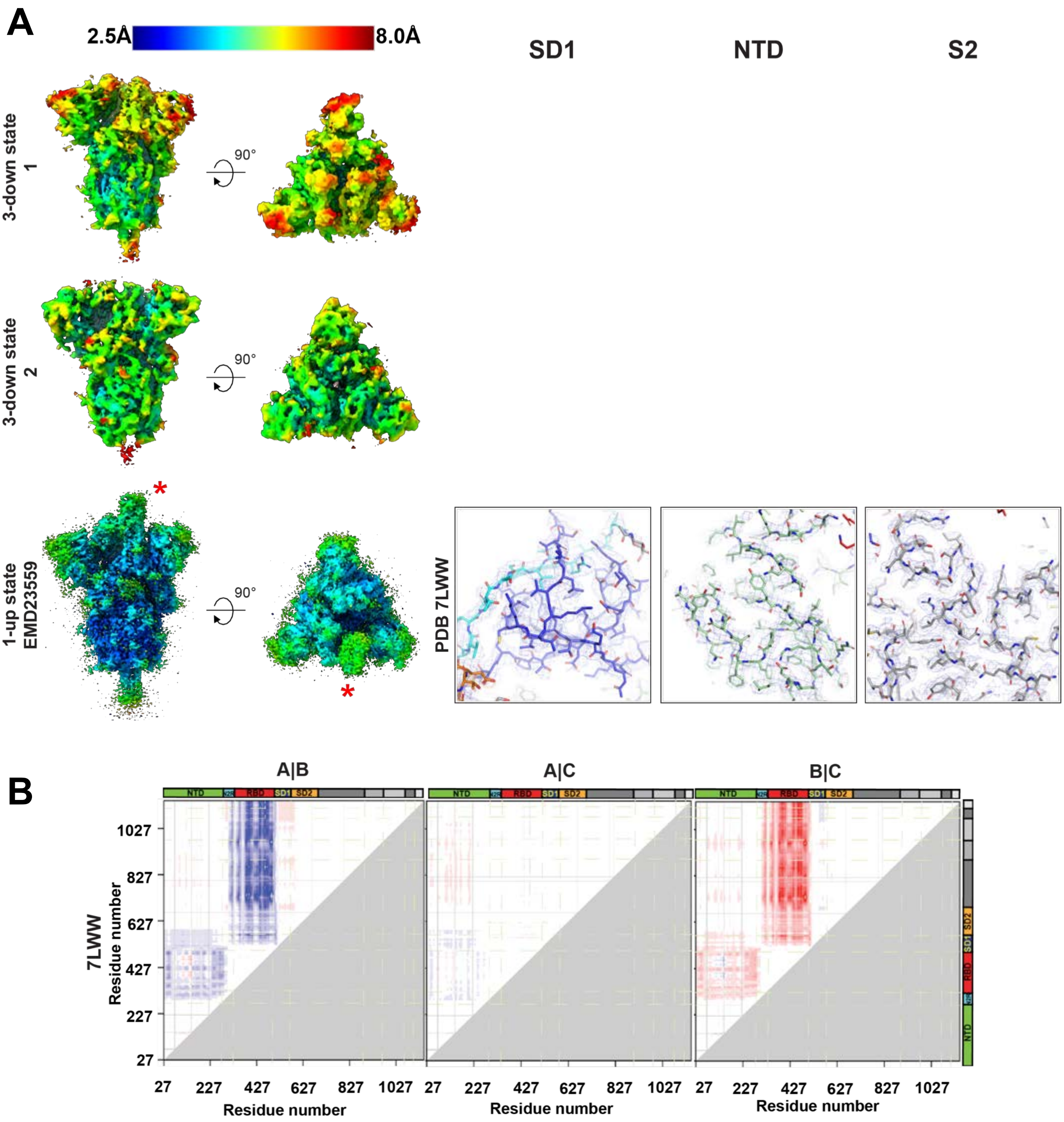


**Supplemental Item 16, continued. Quality assessment of the cryo-EM map and model fitting and difference distance matrices (DDM) showing structural changes between different protomers of the S-GSAS-B.1.1.28 SARS-CoV-2 S ectodomain, Related to Figure 6. A. Left.** Refined cryo-EM maps colored by local resolution **Right.** Zoom-in images showing the SD1, NTD and S2 regions in the structures. The cryo-EM map is shown as a mesh surface and the fitted model is in cartoon representation, with residues shown as sticks. **B.** DDM of the 1-RBD-up structure.

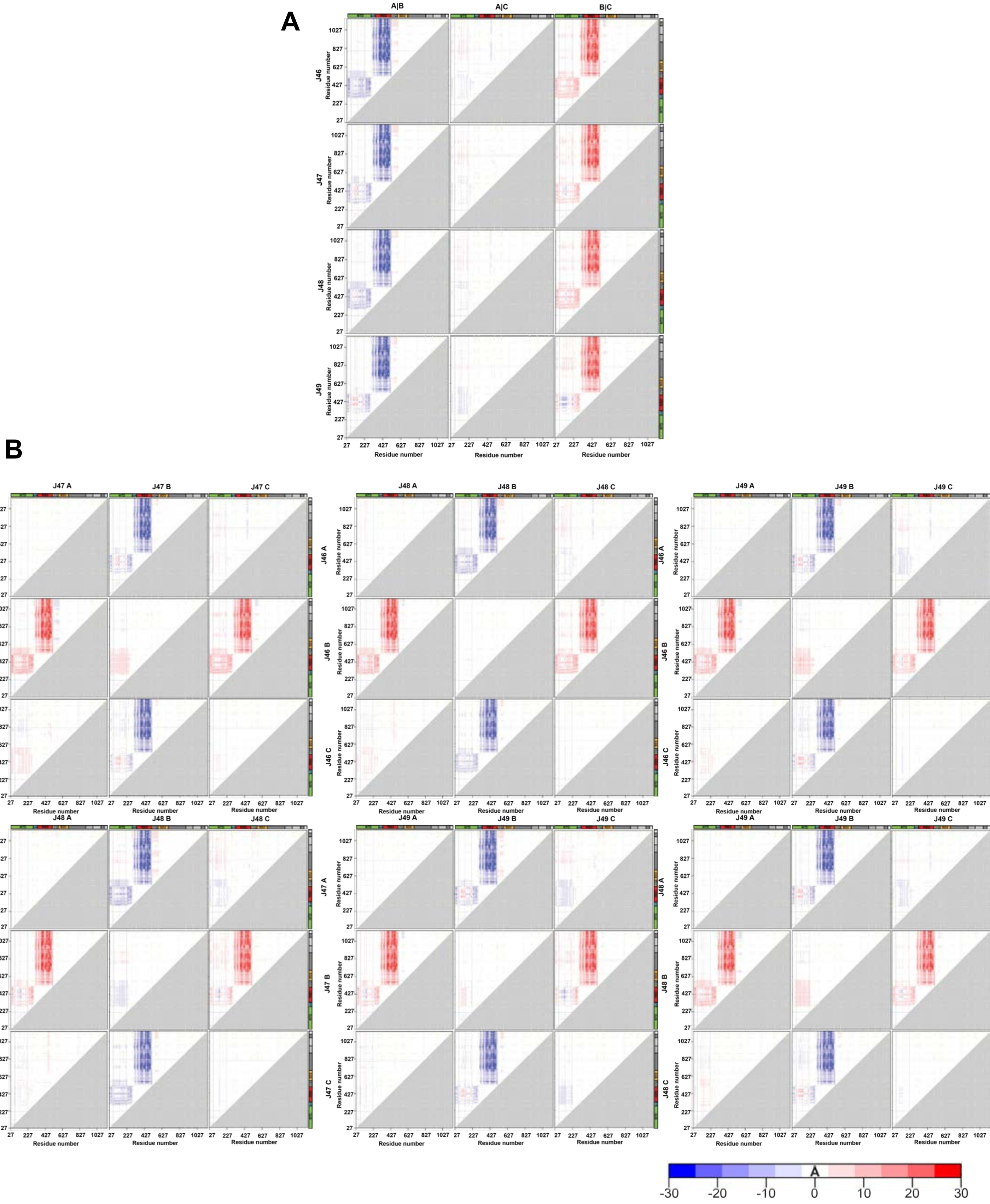


**Supplemental Item 17. Difference distance matrices (DDM) showing structural changes between different protomers for the S-GSAS-B.1.351 SARS-CoV-2 S ectodomain structures. Related to Figure 6. A.** DDM of the 1-RBD up structures comparing each chain of a structure to the other chains of the same structure. **B.** DDM of the 1-RBD up structures comparing each chain to each chains of the other 1-up structures.

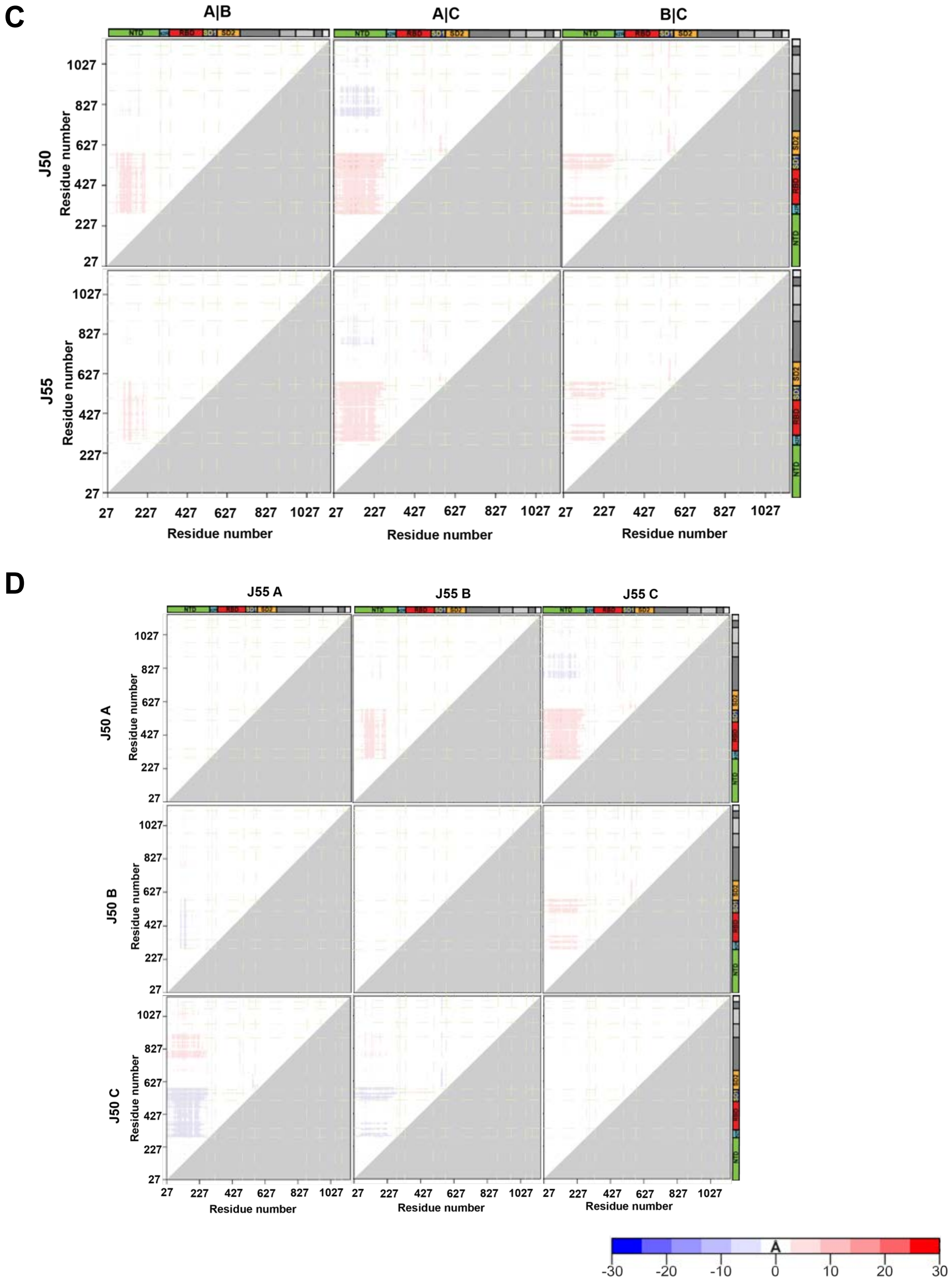


**Supplemental Item 17 continued. Difference distance matrices (DDM) showing structural changes between different protomers for the S-GSAS-B.1.351 SARS-CoV-2 S ectodomain structures. Related to Figure 6. C.** DDM of the 3-RBD down structures comparing each chain of a structure to the other chains of the same structure. **D.** DDM of the 3-RBD-down structures comparing each chain to each chains of the other 3-down structures.

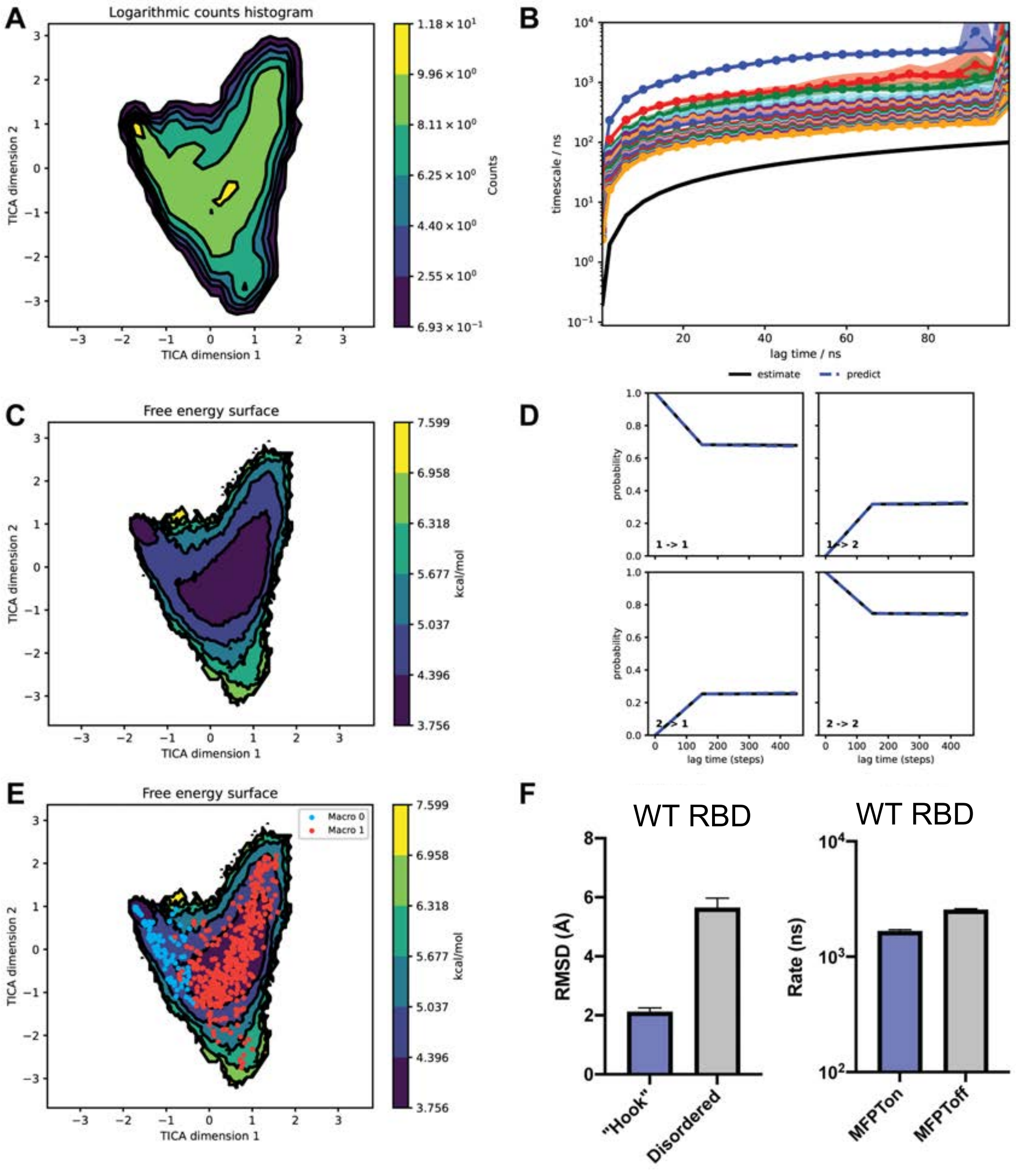


**Supplemental Item 18, Related to Figure 6. Markov model validation and results for WT RBD simulations. A.** Histogram of counts plotted along TICA components 1 and 2. **B.** Implied timescales (ITS) plot. Shaded region represents 95% confidence interval. **C.** Free energy surface plotted along TICA components 1 and 2. **D.** Chapman-Kolmogorov test for the Markov model. **E.** Free energy surface with the cluster state assignments. **F.** (left) Weighted RMSD values for the RBD "Hook" in each state. (right) Mean first passage times for transitions to ($MFPT_{on}$) and from ($MFPT_{off}$) the Disordered state.

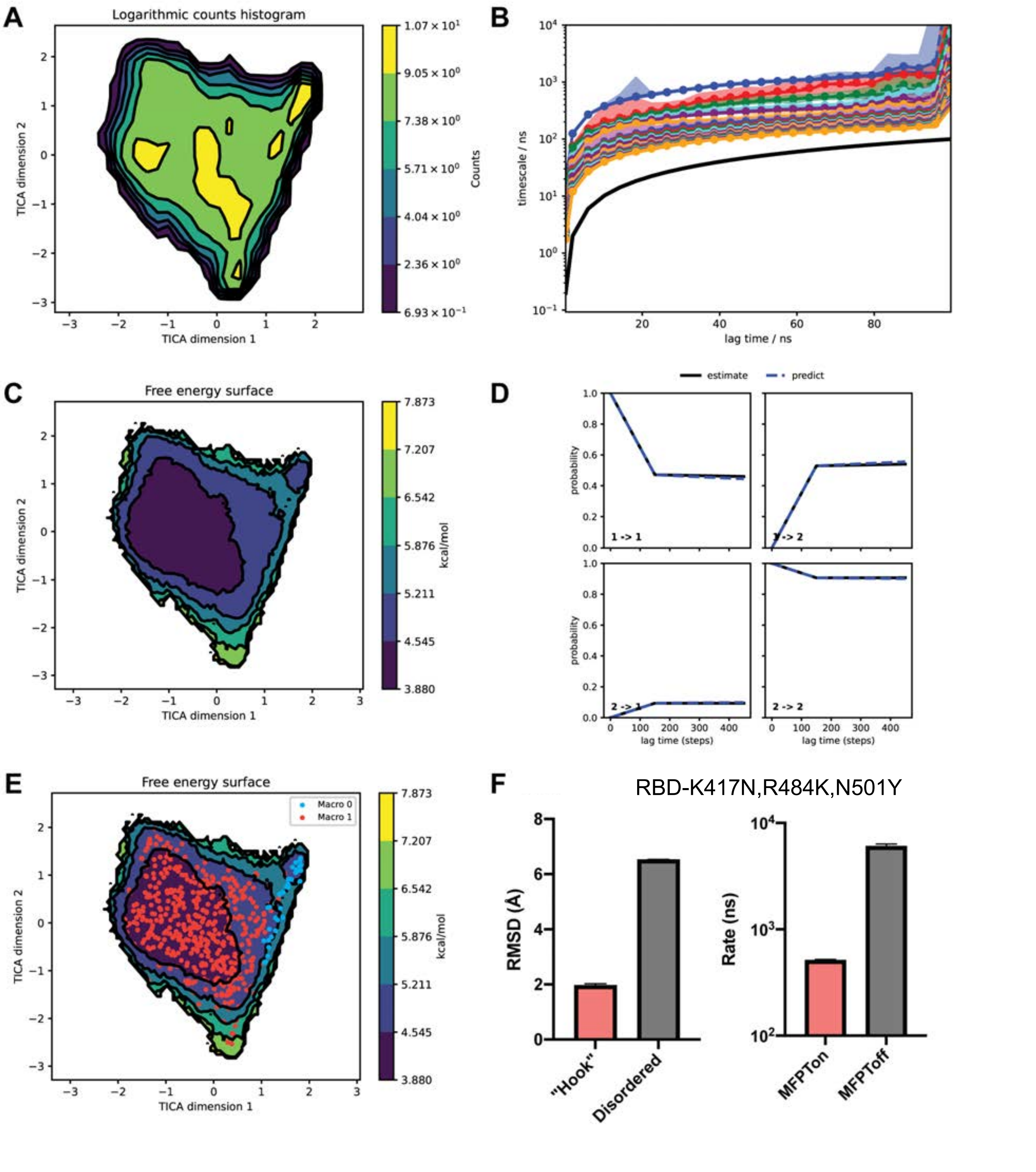


**Supplemental Item 19, Related to Figure 6. Markov model validation and results for K417N, E484K, N501Y RBD simulations. A.** Histogram of counts plotted along TICA components 1 and 2. **B.** Implied timescales (ITS) plot. Shaded region represents 95% confidence interval. **C.** Free energy surface plotted along TICA components 1 and 2. **D.** Chapman-Kolmogorov test for the Markov model. **E.** Free energy surface with the cluster state assignments. **F.** (left) Weighted RMSD values for the RBD "Hook" in each state. (right) Mean first passage times for transitions to ($MFPT_{on}$) and from ($MFPT_{off}$) the disordered state.

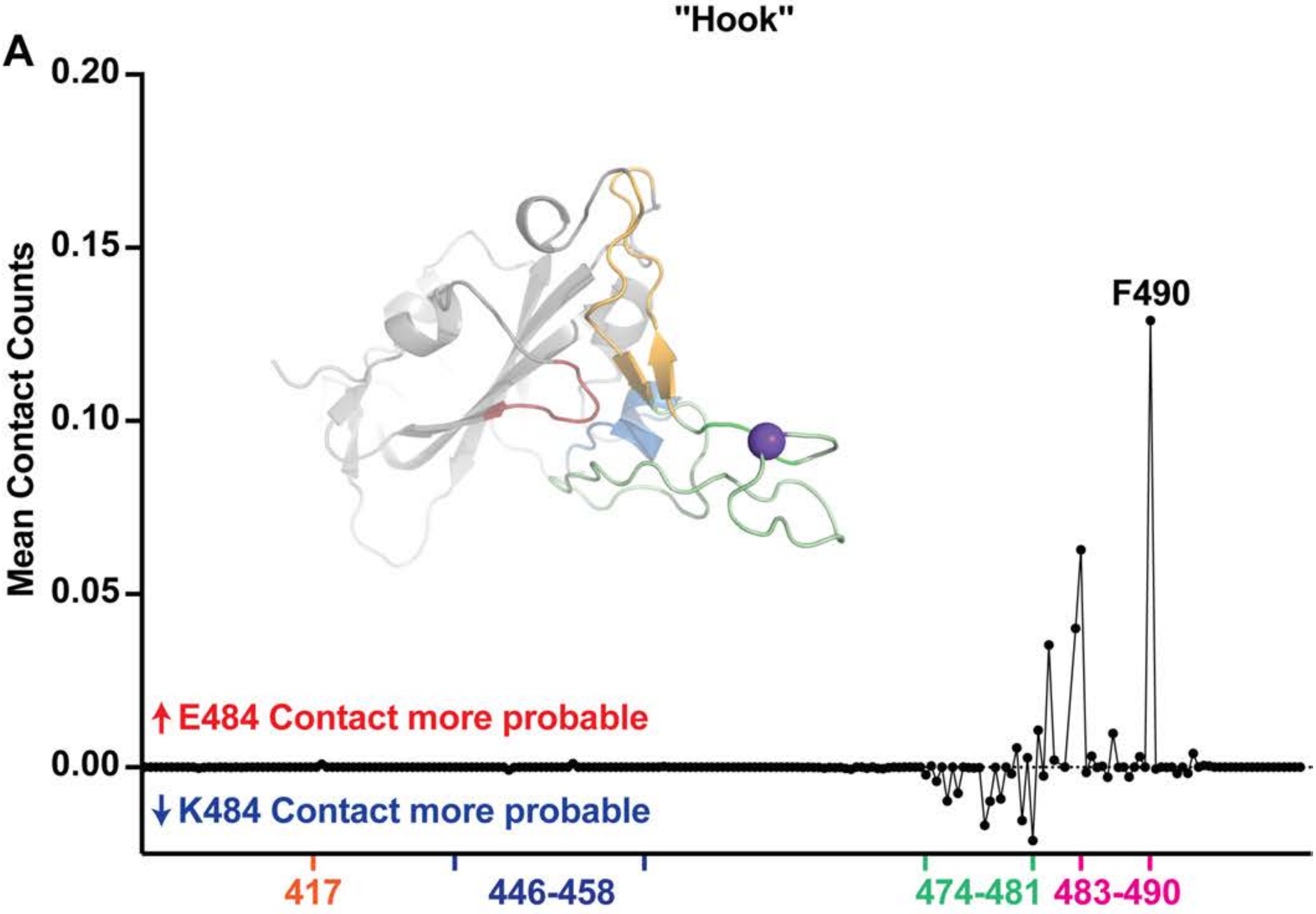
"Hook"
A
0.20
0.15
0.10
0.05
0.00
Mean Contact Counts
F490
↑E484 Contact more probable
↓K484 Contact more probable
417
446-458
474-481
483-490

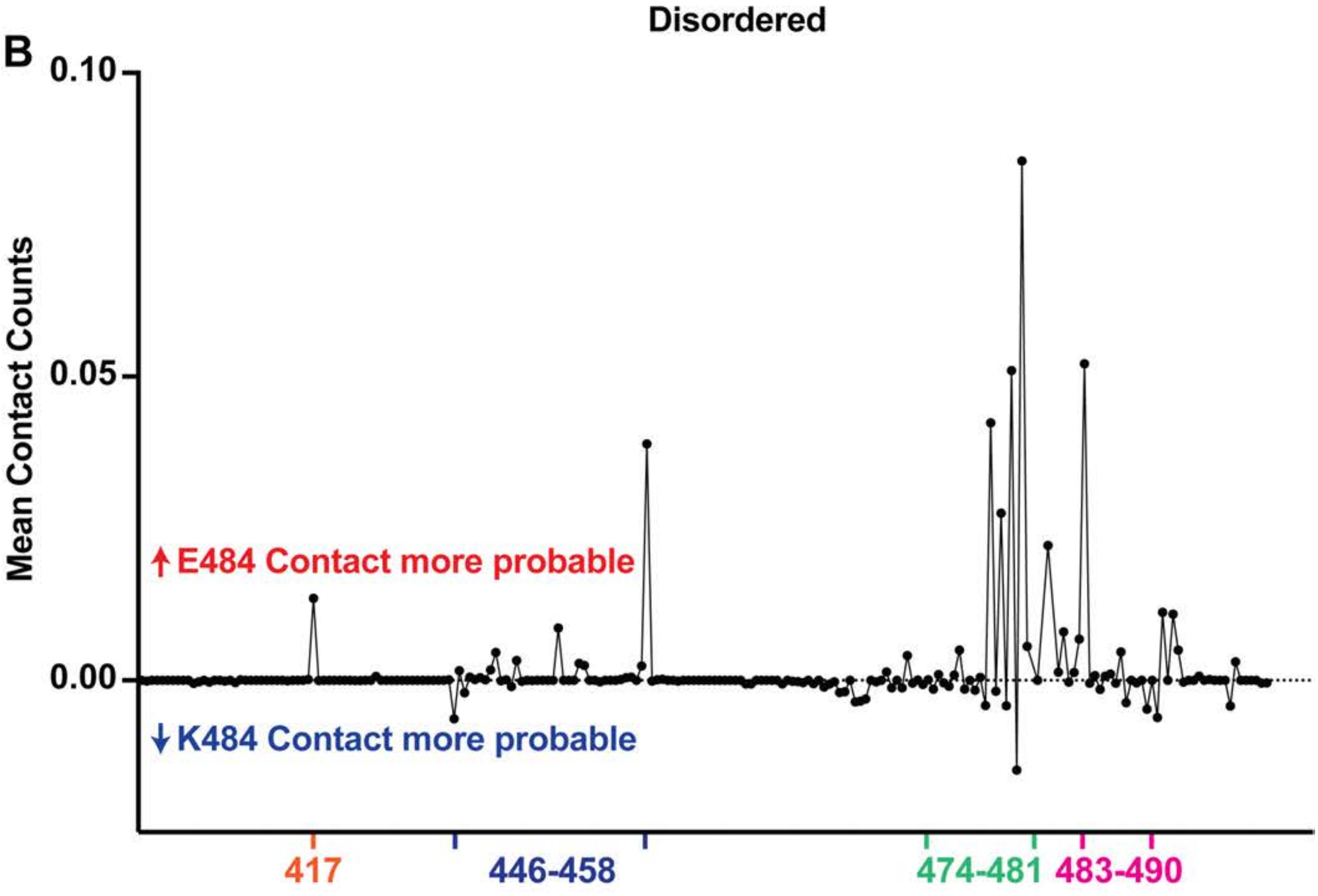
Disordered
B
0.10
0.05
0.00
Mean Contact Counts
↑E484 Contact more probable
↓K484 Contact more probable
417
446-458
474-481
483-490

**Supplemental Item 20, Related to figure 6. Residue 484 Markov model contact counts. A.** Mut subtracted WT "Hook" state contact counts for the residue 484 interacting with RBD residues. (inset) Structure of the RBD highlighting the probed interaction surface including residues 384 to 354 (red), 413 to 425 (blue), 454 to 491, 446 to 453/492 to 500 (light orange ). **B.** Mut subtracted WT Disordered state contact counts for the residue 484 interacting with RBD residues.

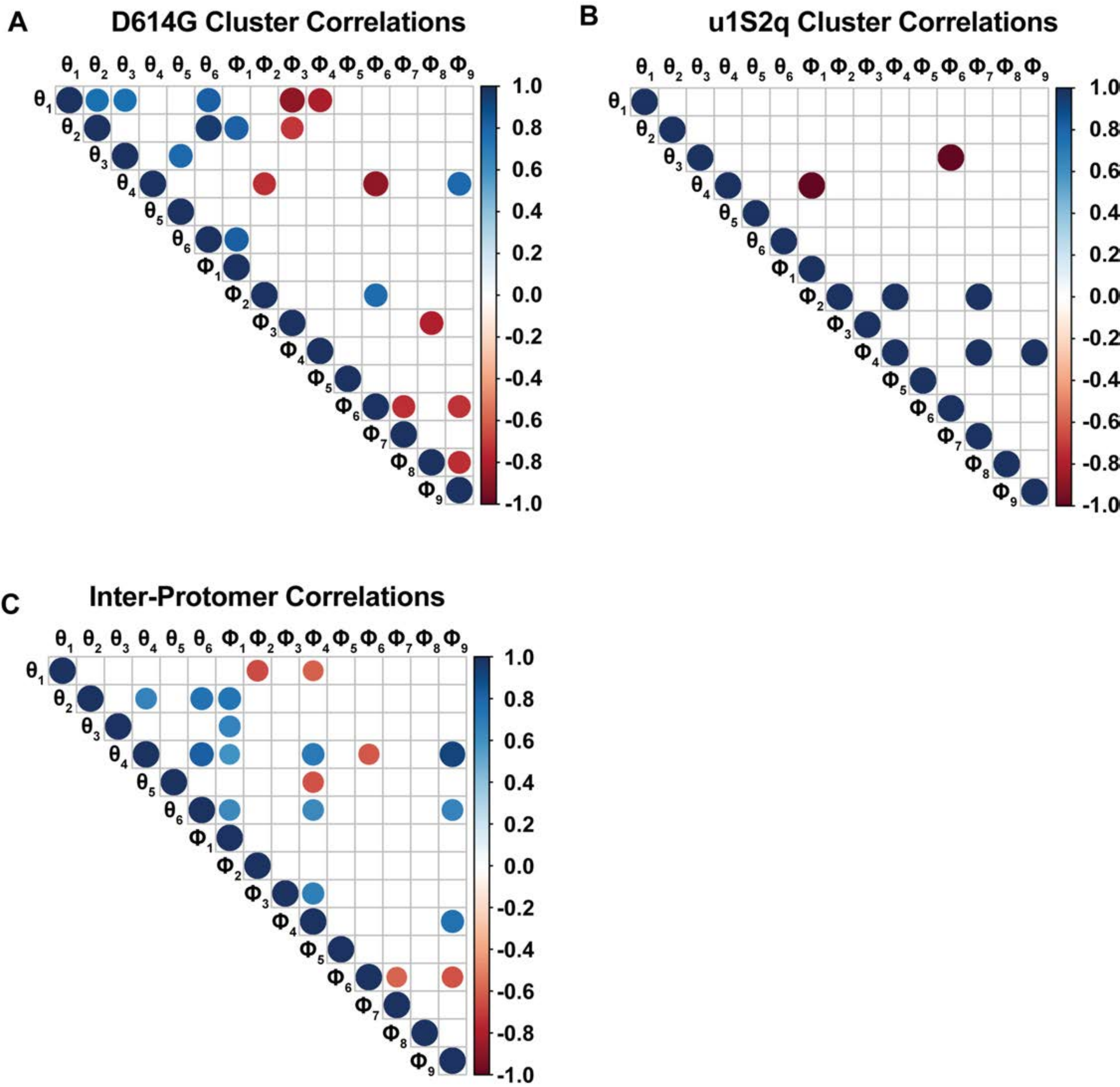


**Supplemental Item 21, Related to Figure 7. Cluster Correlations. A.** Significant Correlations between the inter-protomer D614G cluster structure angle measures (N=9, $p<0.05$). **B.** Significant Correlations between the inter-protomer D614G cluster structure angle measures (N=3, $p<0.05$). **C.** Significant Correlations between all inter-protomer structure angle measures (N=12, $p<0.05$).

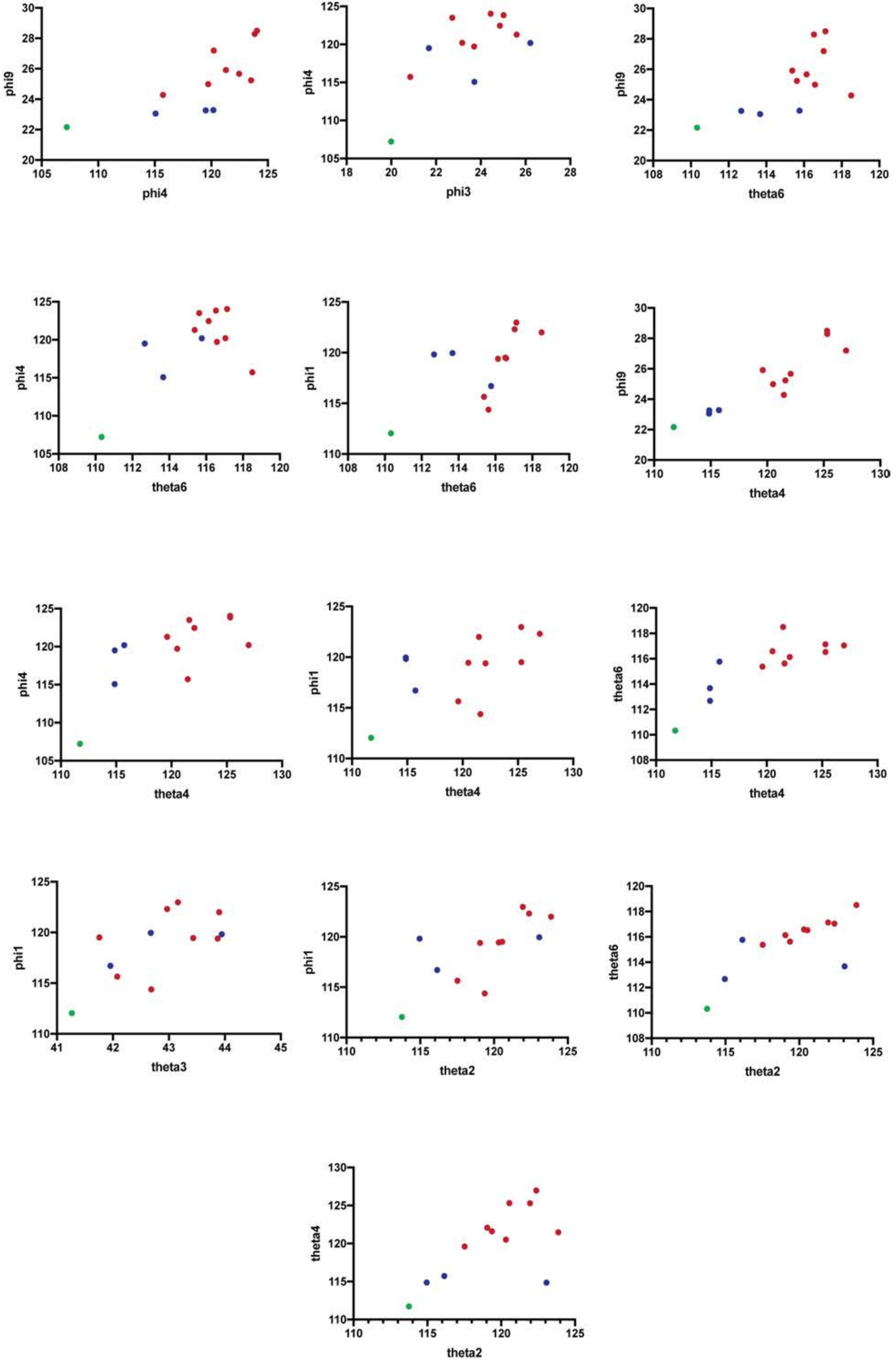

phi9
phi4
phi4
phi3
phi9
theta6
phi4
theta6
phi1
theta6
phi9
theta4
phi4
theta4
phi1
theta4
theta6
theta4
phi1
theta3
phi1
theta2
theta6
theta2
theta4
theta2

**Supplemental Item 22. Related to Figure 7. Significantly positively correlated vector relationship plots in the full inter-protomer set.** Colors are according to clusters in the PCA analysis in Figure 7 panel B.

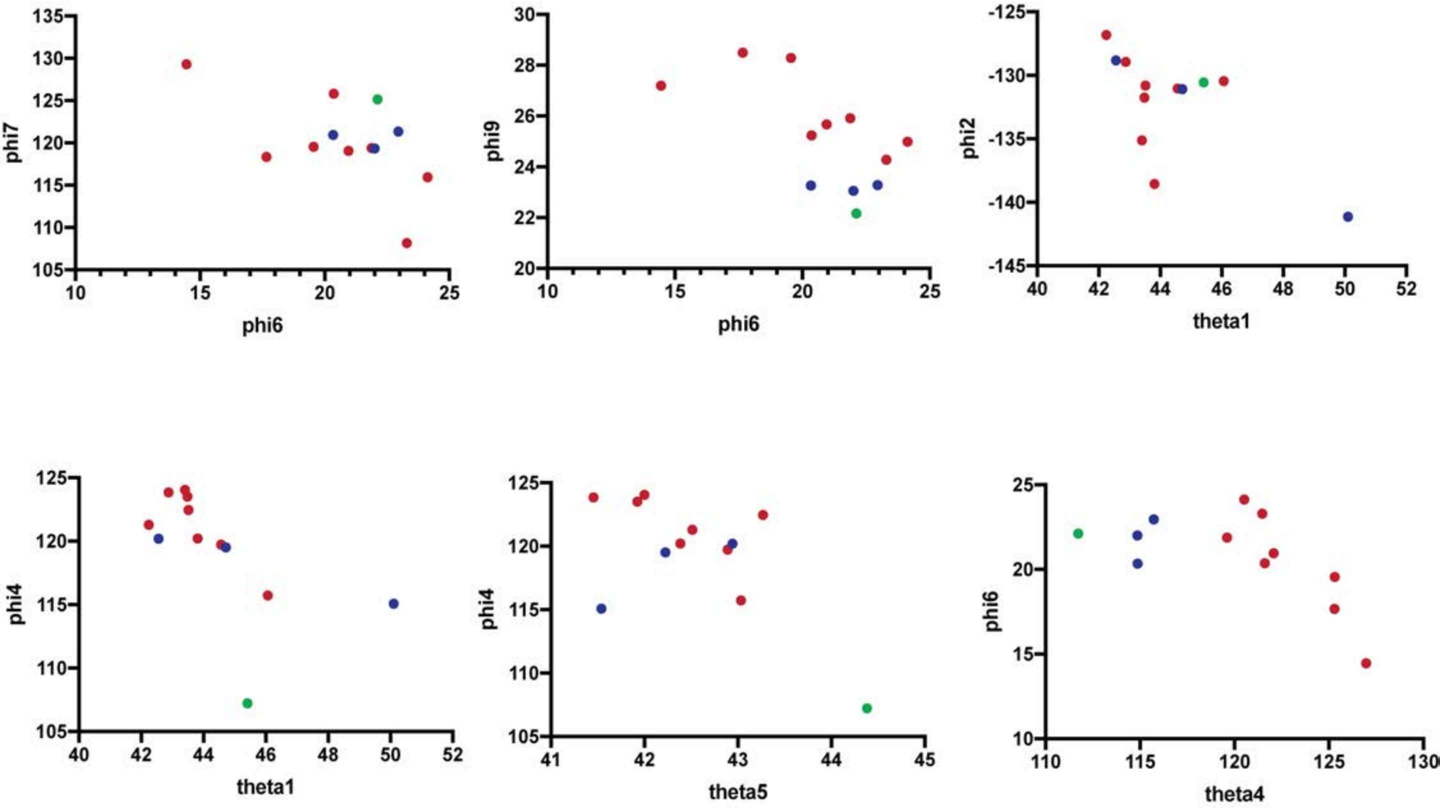


**Supplemental Item 23. Related to Figure 7. Significantly negatively correlated vector relationship plots in the full inter-protomer set.** Colors are according to clusters in the PCA analysis in Figure 7 panel B.

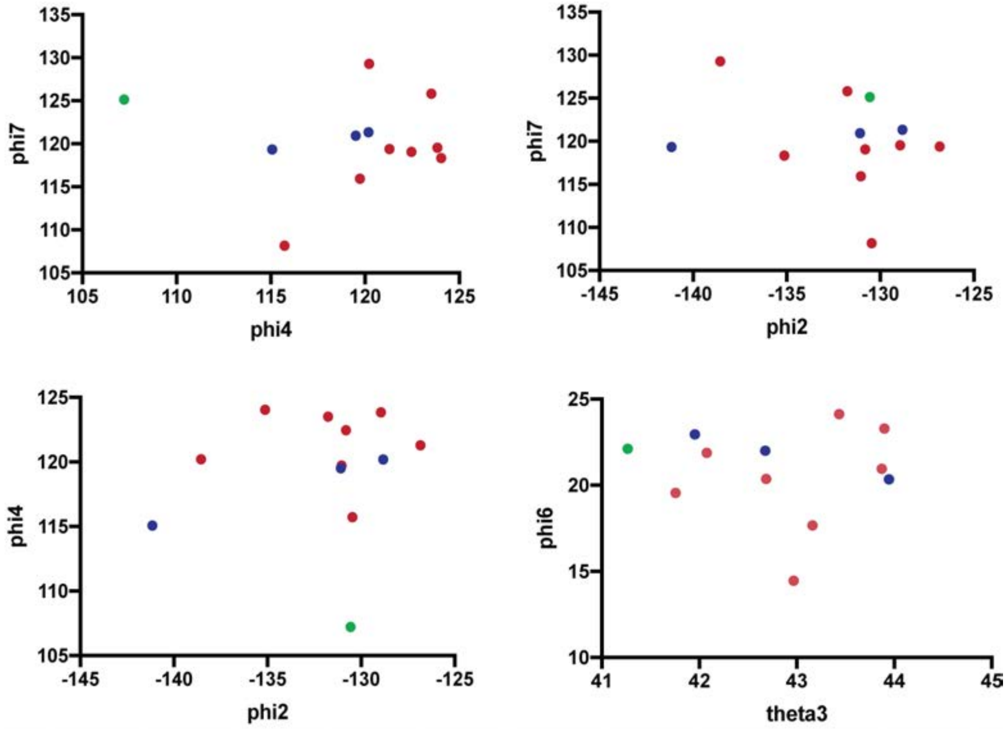


**Supplemental Item 24. Related to Figure 7. Significantly correlated vector relationship plots in the u1S2q cluster inter-protomer set that were not significant in the full set.** Colors are according to clusters in the PCA analysis in Figure 7 panel B.

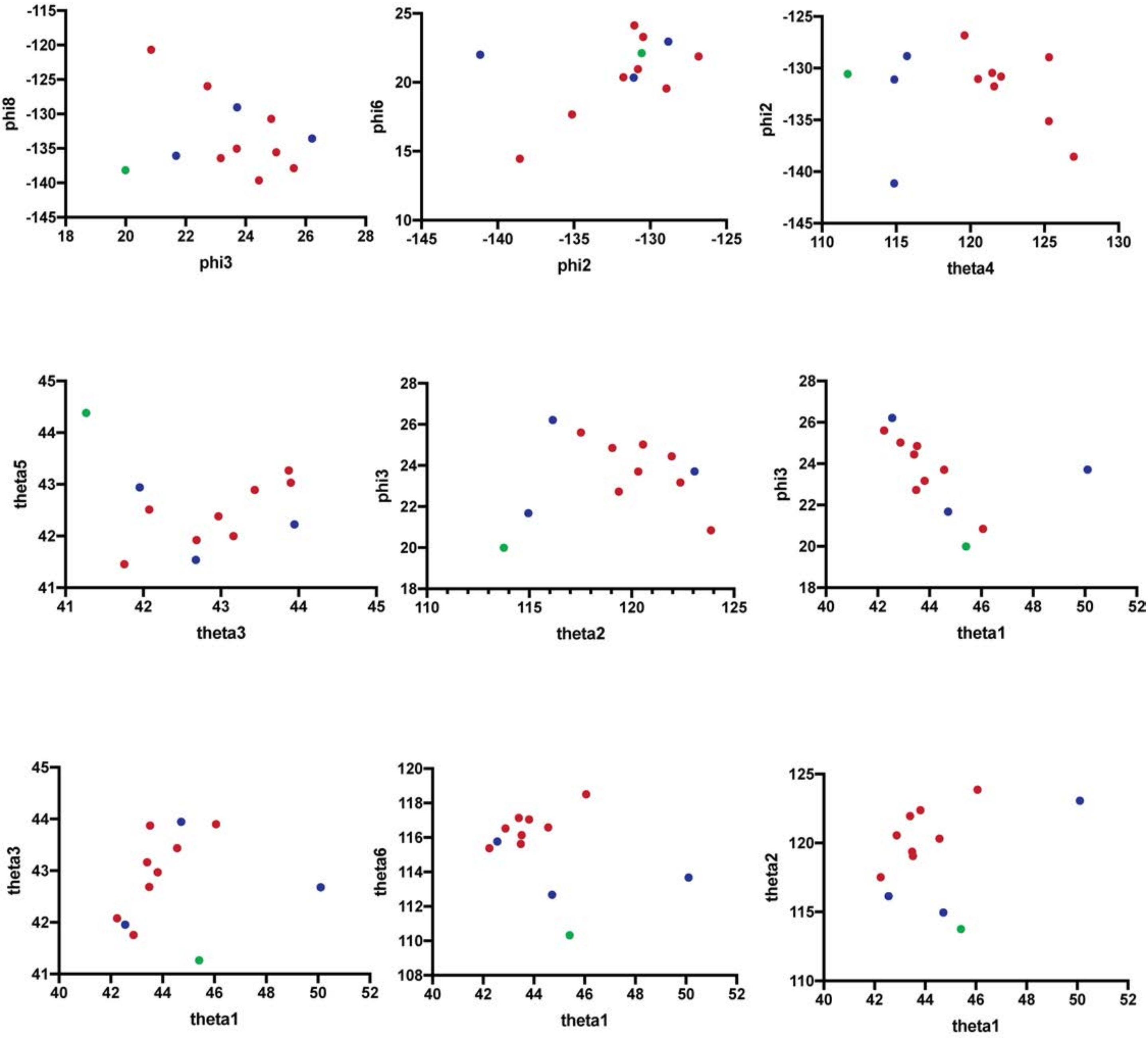


**Supplemental Item 25. Related to Figure 7. Significantly correlated vector relationship plots in the D614G cluster inter-protomer set that were not significant in the full set.** Colors are according to clusters in the PCA analysis in Figure 7 panel B.

**Supplemental Table 1. KD extracted from the kinetics of RBD, NTD and S2 directed antibody Fabs on the spike variants measured by SPR, Related to Figure 2, 3 and 5.**

| | KD (nM) | | | | | | | | |
|---|---|---|---|---|---|---|---|---|---|
| | **ACE2** | **DH1041** | **DH1043** | **DH1050.1** | **DH1050.2** | **DH1052** | **DH1058** | **2G12** | **CR3022** |
| **S-GSAS/PP** | 177.73 | 0.46 | 1.79 | 21.89 | 76.93 | 502.78 | 129.12 | 209.55 | 132.66 |
| **S-GSAS** | 189.53 | 0.07 | 2.04 | 18.63 | 72.33 | 377.40 | 129.94 | 237.34 | 52.27 |
| **S-GSAS-D614G** | 218.29 | 0.76 | 2.24 | 20.74 | 70.82 | 576.94 | 133.50 | 294.87 | 119.39 |
| **S-GSAS-∆FV** | 63.00 | 1.05 | 2.17 | 5.90 | 26.89 | 693.76 | 132.61 | 223.34 | 118.80 |
| **S-GSAS-D614G-H69/V70del** | 227.91 | 0.28 | 2.70 | 6.16 | 15.76 | 417.44 | 144.13 | 297.27 | 62.24 |
| **S-GSAS-D614G-Y453F** | 52.10 | 0.50 | 3.06 | 18.38 | 47.17 | 467.09 | 147.40 | 333.07 | 121.39 |
| **S-GSAS-D614G-I692V** | 149.93 | 0.56 | 2.79 | 16.20 | 40.19 | 404.57 | 144.15 | 269.11 | 73.05 |
| **S-GSAS-B.1.1.7** | 45.20 | 1.04 | 1.74 | 15.51 | 36.12 | 356.49 | 132.06 | 377.27 | 118.83 |
| **S-GSAS-D614G-N501Y** | 51.80 | 0.47 | 2.26 | 17.48 | 37.05 | 624.84 | 130.13 | 307.89 | 119.57 |
| **S-GSAS-B.1.351** | 72.63 | 0.45 | 1.97 | 19.48 | 26.83 | 483.03 | 147.41 | 260.07 | 116.39 |
| **S-GSAS-B.1.1.28** | 93.72 | 1.18 | 1.81 | 18.93 | 66.35 | 631.09 | 134.53 | 239.45 | 59.60 |
| **S-GSAS-D614G-E484K** | 181.43 | 1.08 | 1.96 | 17.50 | 63.56 | 590.51 | 131.95 | 264.09 | 134.82 |
| **S-GSAS-D614G-K417N** | 92.76 | 0.48 | 1.70 | 15.78 | 48.29 | 399.24 | 122.87 | 259.57 | - |

**Supplemental Table 2 continued. Cryo-EM data collection and refinements statistics for the S-GSAS-ΔFV, S-GSAS-B.1.1.7, S-GSAS-B.1.1.28 and S-GSAS-B.1.351 spike ectodomain variants, Related to Figure 2, 3, 4 and 6.**

| | S-GSAS-ΔFV | | | | | | | | |
|---|---|---|---|---|---|---|---|---|---|
| | 3 down | | | | 1up | | | 2up | New State |
| **PDB ID** | 7LWI | 7LWJ | 7LWK | 7LWL | 7LWM | 7LWN | 7LWO | 7LWP | 7LWQ |
| **EMDB ID** | 23546 | 23547 | 23548 | 23549 | 23550 | 23551 | 23552 | 23553 | 23554 |
| **Data collection and processing** | | | | | | | | | |
| Microscope | FEI Titan Krios | | | | | | | | |
| Detector | Gatan K3 | | | | | | | | |
| Magnification | 81,000 | | | | | | | | |
| Voltage (kV) | 300 | | | | | | | | |
| Electron exposure (e–/Å2) | 52.9 | | | | | | | | |
| Defocus range (μm) | 2.32-0.77 | | | | | | | | |
| Pixel size (Å) | 1.069 | | | | | | | | |
| Reconstruction software | cryoSparc | | | | | | | | |
| Symmetry imposed | C1 | C1 | C1 | C1 | C1 | C1 | C1 | C1 | C1 |
| Initial particle images (no.) | 8,060,973 | | | | | | | | |
| Final particle images (no.) | 214,521 | 193,850 | 276,950 | 340,327 | 412,536 | 346,973 | 367,939 | 179,963 | 54,693 |
| Map resolution (Å) | 3.07 | 3.24 | 2.92 | 2.84 | 2.83 | 2.94 | 2.85 | 3.01 | 3.44 |
| FSC threshold | 0.143 | 0.143 | 0.143 | 0.143 | 0.143 | 0.143 | 0.143 | 0.143 | 0.143 |
| **Refinement** | | | | | | | | | |
| Initial model used | 7JMO, 7KDK | | | | 7JMO, 7KDL | | | 7JMO, 6X2B | 7JMO, 7KDK |
| Model resolution (Å) | 3.07 | 3.24 | 2.92 | 2.84 | 2.83 | 2.94 | 2.85 | 3.01 | 3.44 |
| FSC threshold | 0.143 | 0.143 | 0.143 | 0.143 | 0.143 | 0.143 | 0.143 | 0.143 | 0.143 |
| **Model composition** | | | | | | | | | |
| Nonhydrogen atoms | 23,847 | 23,847 | 23,847 | 23,847 | 23,825 | 23,825 | 23,825 | 23,069 | 19,190 |
| Protein residues | 3,003 | 3,003 | 3,003 | 3,003 | 2,980 | 2,980 | 2,980 | 2,982 | 2,425 |
| **R.m.s. deviations** | | | | | | | | | |
| Bond lengths (Å) | 0.012 | 0.013 | 0.012 | 0.012 | 0.013 | 0.013 | 0.013 | 0.012 | 0.013 |
| Bond angles (°) | 1.873 | 1.902 | 1.867 | 1.885 | 1.866 | 1.865 | 1.870 | 1.897 | 1.780 |
| **Validation** | | | | | | | | | |
| MolProbity score | 0.95 | 1.09 | 0.89 | 0.88 | 1.00 | 0.99 | 0.96 | 0.98 | 0.97 |
| Clashscore | 0.15 | 0.28 | 0.21 | 0.17 | 0.28 | 0.38 | 0.30 | 0.15 | 0.05 |
| Poor rotamers (%) | 0.31 | 0.38 | 0.27 | 0.23 | 0.35 | 0.31 | 0.20 | 0.82 | 0.85 |
| EM ringer score | 3.32 | 1.86 | 3.57 | 3.92 | 3.98 | 3.93 | 3.55 | 3.06 | 2.56 |
| **Ramachandran plot** | | | | | | | | | |
| Favored (%) | 94.18 | 92.05 | 95.57 | 95.43 | 94.17 | 94.86 | 94.94 | 93.70 | 93.09 |
| Allowed (%) | 5.62 | 7.75 | 4.37 | 4.30 | 5.48 | 4.87 | 4.73 | 5.82 | 6.53 |
| Disallowed (%) | 0.20 | 0.20 | 0.07 | 0.27 | 0.34 | 0.27 | 0.34 | 0.48 | 0.38 |

**Supplemental Table 2 continued. Cryo-EM data collection and refinements statistics for the S-GSAS-∆FV, S-GSAS-B.1.1.7, S-GSAS-B.1.1.28 and S-GSAS-B.1.351 spike ectodomain variants, Related to Figure 2, 3, 4 and 6.**

| | S-GSAS-B.1.1.7 | | | | u1S2q |
|---|---|---|---|---|---|
| | **3down** | | **1up** | | **3 down** |
| **PDB ID** | 7LWS | 7LWT | 7LWU | 7LWV | 7M0J |
| **EMDB ID** | 23555 | 23556 | 23557 | 23558 | 23613 |
| **Data collection and processing** | | | | | |
| Microscope | | | FEI Titan Krios | | |
| Detector | | | Gatan K3 | | |
| Magnification | | | 81,000 | | |
| Voltage (kV) | | | 300 | | |
| Electron exposure (e–/Å2) | | 52.65 | | | 66.82 |
| Defocus range (μm) | | 2.32-0.78 | | | 0.55-2.94 |
| Pixel size (Å) | | 1.069 | | | 1.058 |
| Reconstruction software | | | cryoSparc | | |
| Symmetry imposed | C1 | C1 | C1 | C1 | C1 |
| Initial particle images (no.) | | 4,679,495 | | | 906,517 |
| Final particle images (no.) | 171,287 | 163,471 | 125,383 | 238,400 | 139,267 |
| Map resolution (Å) | 3.22 | 3.19 | 3.22 | 3.12 | 3.52 |
| FSC threshold | 0.143 | 0.143 | 0.143 | 0.143 | 0.143 |
| **Refinement** | | | | | |
| Initial model used | 7JMO, 7KDK | | 7JMO, 7KDL | | 6VXX |
| Model resolution (Å) | 3.22 | 3.19 | 3.22 | 3.12 | 3.52 |
| FSC threshold | 0.143 | 0.143 | 0.143 | 0.143 | 0.143 |
| **Model composition** | | | | | |
| Nonhydrogen atoms | 23,439 | 23,760 | 23,578 | 23,564 | 22,827 |
| Protein residues | 3,000 | 2,980 | 2,980 | 2,980 | 2,916 |
| **R.m.s. deviations** | | | | | |
| Bond lengths (Å) | 0.012 | 0.012 | 0.012 | 0.012 | 0.012 |
| Bond angles (°) | 1.879 | 1.851 | 1.830 | 1.845 | 1.954 |
| **Validation** | | | | | |
| MolProbity score | 0.86 | 0.95 | 1.04 | 0.95 | 1.21 |
| Clashscore | 0.06 | 0.23 | 0.45 | 0.15 | 0.13 |
| Poor rotamers (%) | 0.50 | 0.79 | 0.90 | 0.47 | 1.68 |
| EM ringer score | 3.58 | 3.87 | 3.59 | 3.99 | 2.38 |
| **Ramachandran plot** | | | | | |
| Favored (%) | 95.18 | 94.84 | 94.43 | 94.29 | 92.16 |
| Allowed (%) | 4.41 | 4.65 | 5.23 | 5.40 | 7.38 |
| Disallowed (%) | 0.41 | 0.51 | 0.34 | 0.31 | 0.46 |

**Supplemental Table 2 continued. Cryo-EM data collection and refinements statistics for the S-GSAS-ΔFV, S-GSAS-B.1.1.7, S-GSAS-B.1.1.28 and S-GSAS-B.1.351 spike ectodomain variants, Related to Figure 2, 3, 4 and 6.**

| | **S-GSAS-B.1.1.28 (Brazilian)** | **S-GSAS-B.1.351 (South African)** | | | | | | |
|---|---|---|---|---|---|---|---|---|
| | **1up** | **3 down** | **3down-Consensus** | **2up** | **1up** | | | |
| **PDB ID** | 7LWW | 7LYL | 7LYM | 7LYK | 7LYN | 7LYO | 7LYP | 7LYQ |
| **EMDB ID** | 23559 | 23594 | 23595 | 23593 | 23596 | 23597 | 23598 | 23599 |
| **Data collection and processing** | | | | | | | | |
| Microscope | FEI Titan Krios | FEI Titan Krios | | | | | | |
| Detector | Gatan K3 | Gatan K3 | | | | | | |
| Magnification | 81,000 | 81,000 | | | | | | |
| Voltage (kV) | 300 | 300 | | | | | | |
| Electron exposure (e–/Å2) | 52.9 | 51.16 | | | | | | |
| Defocus range (μm) | 2.33-0.77 | 2.46-0.64 | | | | | | |
| Pixel size (Å) | 1.069 | 1.069 | | | | | | |
| Reconstruction software | cryoSparc | cryoSparc | | | | | | |
| Symmetry imposed | C1 | C1 | C1 | C1 | C1 | C1 | C1 | C1 |
| Initial particle images (no.) | 5,332,605 | 4,740,302 | | | | | | |
| Final particle images (no.) | 331,799 | 65,713 | 212,753 | 200,301 | 404,423 | 381,430 | 61,823 | 199,601 |
| Map resolution (Å) | 3.00 | 3.72 | 3.57 | 3.65 | 3.32 | 3.32 | 4.05 | 3.34 |
| FSC threshold | 0.143 | 0.143 | 0.143 | 0.143 | 0.143 | 0.143 | 0.143 | 0.143 |
| **Refinement** | | | | | | | | |
| Initial model used | 7JMO, 7KDL | 7JMO, 7KDK | | 7JMO, 6X2B | 7JMO, 7KDL | | | |
| Model resolution (Å) | 3.00 | 3.72 | 3.57 | 3.65 | 3.32 | 3.32 | 4.05 | 3.34 |
| FSC threshold | 0.143 | 0.143 | 0.143 | 0.143 | 0.143 | 0.143 | 0.143 | 0.143 |
| **Model composition** | | | | | | | | |
| Nonhydrogen atoms | 23,803 | 23,828 | 23,912 | 23,444 | 23,642 | 23,649 | 23,506 | 23,642 |
| Protein residues | 2,982 | 3,003 | 3,005 | 2,983 | 2,985 | 2,982 | 2,985 | 2,985 |
| **R.m.s. deviations** | | | | | | | | |
| Bond lengths (Å) | 0.012 | 0.012 | 0.012 | 0.012 | 0.012 | 0.012 | 0.012 | 0.012 |
| Bond angles (°) | 1.869 | 1.787 | 1.799 | 1.792 | 1.811 | 1.828 | 1.784 | 1.818 |
| **Validation** | | | | | | | | |
| MolProbity score | 1.03 | 1.06 | 0.97 | 1.10 | 1.10 | 1.11 | 1.26 | 1.05 |
| Clashscore | 0.4 | 0.55 | 0.21 | 0.58 | 0.56 | 0.58 | 0.95 | 0.39 |
| Poor rotamers (%) | 0.75 | 0.80 | 0.57 | 0.43 | 0.59 | 0.47 | 1.02 | 0.55 |
| EM ringer score | 3.93 | 2.37 | 2.67 | 2.88 | 3.64 | 3.68 | 1.82 | 3.4 |
| **Ramachandran plot** | | | | | | | | |
| Favored (%) | 94.33 | 94.51 | 94.28 | 94.00 | 93.91 | 93.68 | 92.44 | 93.74 |
| Allowed (%) | 5.29 | 5.32 | 5.52 | 5.70 | 5.92 | 5.87 | 6.84 | 5.82 |
| Disallowed (%) | 0.38 | 0.17 | 0.20 | 0.31 | 0.17 | 0.44 | 0.72 | 0.44 |

***Supplemental Table 3.* Tested projections for Markov model building.**

| **Identifier** | **Projection** | **Cut-off** | **Atoms** | **Residues** |
|---|---|---|---|---|
| **A** | Distances | - | C-α | 452-494 |
| **B** | Contacts | 3.5 Å | All | 452-494 |
| **C** | Contacts | 3.5 Å | N and O | 471-480, 484-488 |
| **D** | Torsions | - | Backbone | 452-494 |
| **E** | Distances | - | C-α | 452-494 |
| **F** | Distances | - | C-α | 471-480, 484-488 |
| **G** | Torsions | - | Backbone | 471-480, 484-488 |
| **H** | Distances | - | N, O, C-β | 471-480, 484-488 |
| **I** | Distances | - | C-α | 347-353, 413-425, 446-500 |
